# Immunogenomic, single-cell and spatial dissection of CD8^+^T cell exhaustion reveals critical determinants of cancer immunotherapy

**DOI:** 10.1101/2021.11.22.468617

**Authors:** Stefan Naulaerts, Daniel M Borras, Asier Antoranz Martinez, Julie Messiaen, Yannick Van Herck, Lendert Gelens, Tom Venken, Isaure Vanmeerbeek, Sanket More, Jenny Sprooten, Oliver Bechter, Gabriele Bergers, Adrian Liston, Steven De Vleeschouwer, Benoit J Van Den Eynde, Diether Lambrechts, Jannie Borst, Francesca Bosisio, Sabine Tejpar, Frederik De Smet, Abhishek D Garg

## Abstract

Tumoural-CD8^+^T cells exhibit exhausted or dysfunctional states. Contrary to immunotherapy-responsive exhausted-CD8^+^T cells, the clinical features of dysfunctional-CD8^+^T cells are disputed. Hence, we conducted large-scale multi-omics and multi-dimensional mapping of CD8^+^T cell-states across multiple cancer patient-cohorts. This identified tumour-specific continuum of CD8^+^T cell-states across 6 human cancers, partly imprinted by organ-specific immuno-modulatory niches. Herein, melanoma and glioblastoma enriched prototypical exhausted (CD8^+^T_EXT_) and severely-dysfunctional (CD8^+^T_SDF_) states, respectively. Contrary to CD8^+^T_EXT_, CD8^+^T_SDF_ displayed transcriptomic and epigenetic effector/cytolytic dysfunctions, and dysregulated effector/memory single-cell trajectories, culminating into maladaptive prodeath stress and cell-cycle defects. Suboptimal antigen-priming underscored CD8^+^T_SDF_, which was distinct from immune-checkpoints “rich” CD8^+^T_EXT_, reflecting chronic antigen-stimulation. Continuum variation also existed on tumour spatial-level, with convergent (CD8^+^T_EXT_-supportive vascular regions) and divergent features (dysfunctional CD4^+^T::CD8^+^T_SDF_cell-to-cell interactions) between melanoma and glioblastoma. Globally, IFNγ-IL2 disparities, paucity of intra-tumoural CD4^+^/CD8^+^T cells, and myeloid TGFβ/wound healing responses, distinguished CD8^+^T_SDF_-landscape. Within immuno-oncology clinical-trials, anti-PD1 immunotherapy failed to “reinvigorate” CD8^+^T_SDF_-landscape, and instead facilitated effector-dysfunction and TGFβ/wound healing. However, cellular immunotherapies (dendritic cell-vaccines, adoptive T-cell therapy) ameliorated assorted CD8^+^T_SDF_-landscape disparities, highlighting a roadmap for anti-glioblastoma multimodal-immunotherapy. Collectively, our study comprehensively expands clinical-knowledge on CD8^+^T cell-exhaustion and suggests that tumour-specific, pre-existing CD8^+^T_EXT_/T_SDF_-states, determine immunotherapy-responses.

## INTRODUCTION

Immune-checkpoint blocking (ICB)-immunotherapy has revolutionized oncology.^1–4^ However, not all patients or cancer-types respond to ICBs.^5–7^ Benefits of ICBs are amplified by guiding their application through biomarkers,^1,7–10^ connected to density/diversity of intratumoural-CD8^+^T cells.^1,7,11^ The *modus operandi* of ICBs (especially those targeting PDl/PDLl-axis) entails “reinvigorating” anti-tumour effector/cytolytic functions of pre/early-exhausted CD8^+^T cells.^12–14^ Tumour-induced CD8^+^T cell-exhaustion/dysfunction^12–14^ entails multi-scale remodelling that causes loss of effector/cytolytic functions, paralleling the up-regulation of inhibitory-receptors (IRs), like PD1 or CTLA4.^2,10,12,13^

Tumoural-CD8^+^T cells exhibit heterogenous chronically-sensitised states, that can be broadly classified as differentially exhausted (CD8^+^T_EXT_), or severely dysfunctional (CD8^+^T_SDF_).^2,10,12,15^ Originally defined CD8^+^T_EXT_ include ICB-responsive, pre/early-exhausted CD8^+^T cell-populations, produced under chronic antigen-stimulation.^2,10,16^ Tumoural-CD8^+^T_EXT_ show threshold effector/cytolytic features, memory-differentiation, transcriptional/epigenetic ‘plasticity’, and positive prognostic impact in patient-context.^10,12,17,18^ However, the characteristics and ICB-responsiveness of CD8^+^T_SDF_ population is disputed. Studies with ICB-responsive cancers identify CD8^+^T_SDF_ as late/terminal-CD8^+^T_EXT_ (derived from early-CD8^+^T_EXT_) exhibiting low effector-function but threshold immunotherapy-responsiveness.^2,10,14,16,17,19^ However, studies with ICB-nonresponsive cancers suggest permanently-dysfunctional CD8^+^T_SDF_, proposed to exhibit transcriptional/epigenetic disparities, lack of effector/cytolytic activity, and null-to-negative prognostic impact.^3,9,15,16,18^ Since most ICB-nonresponsive cancers exhibit low antigenicity/immunogenicity, CD8^+^T_SDF_ herein might be distinct from late-CD8^+^T_EXT_, owing to former’s origin from suboptimally (rather than chronically) antigen-stimulated CD8^+^T cells. These proposed CD8^+^T_EXT_/T_SDF_ definitions haven’t been sufficiently verified via cross-omics clinical-approaches thereby creating a paucity of tangible biomarkers for differentiating them.

Current CD8^+^T cell-exhaustion biomarkers mainly comprise of multiple-IRs, and interferon-γ (IFNγ)-signalling or memory-differentiation determinants.^1,2,9,17^ Besides being limited in scope and inconsistently applied across studies,^2^ majority of these cannot discriminate CD8^+^T_EXT_from CD8^+^T_SDF_. Moreover, most clinical studies interpret these biomarkers in a pan-cancer manner, based on non-spatial omics.^2,14,15,17^ As such, these approaches underestimate the impact of divergent cancer ecosystems and their spatial tissue-characteristics. Thus, there is an urgent need to fully dissect clinical-characteristics of CD8^+^T_SDF_ (and their divergence from CD8^+^T_EXT_), to guide future immunotherapy-regimen.

Hence, we conducted deep exploration of clinical CD8^+^T cell-states, integrating diverse tumour-contexts (>4000 patients across 7 cancer-types, spanning 24 distinct clinical-cohorts). We used a literature-derived ‘consensus’ CD8^+^T cell-signature, complemented by other interlocking signatures/scores, to drive a multi-scale (transcriptomic, epigenomic, proteins) and multi-dimensional mapping (tumour-bulk vs. single-cell; non-spatial vs. spatial), of existing large-scale patient datasets. Broad conclusions derived herewith were further verified in our own original cancer patient-cohorts, via tumour-tissue’s multiplex protein analyses at singlecell resolution. Schematic overview of our study-design is depicted in **Fig.1a.**

**Figure 1.**
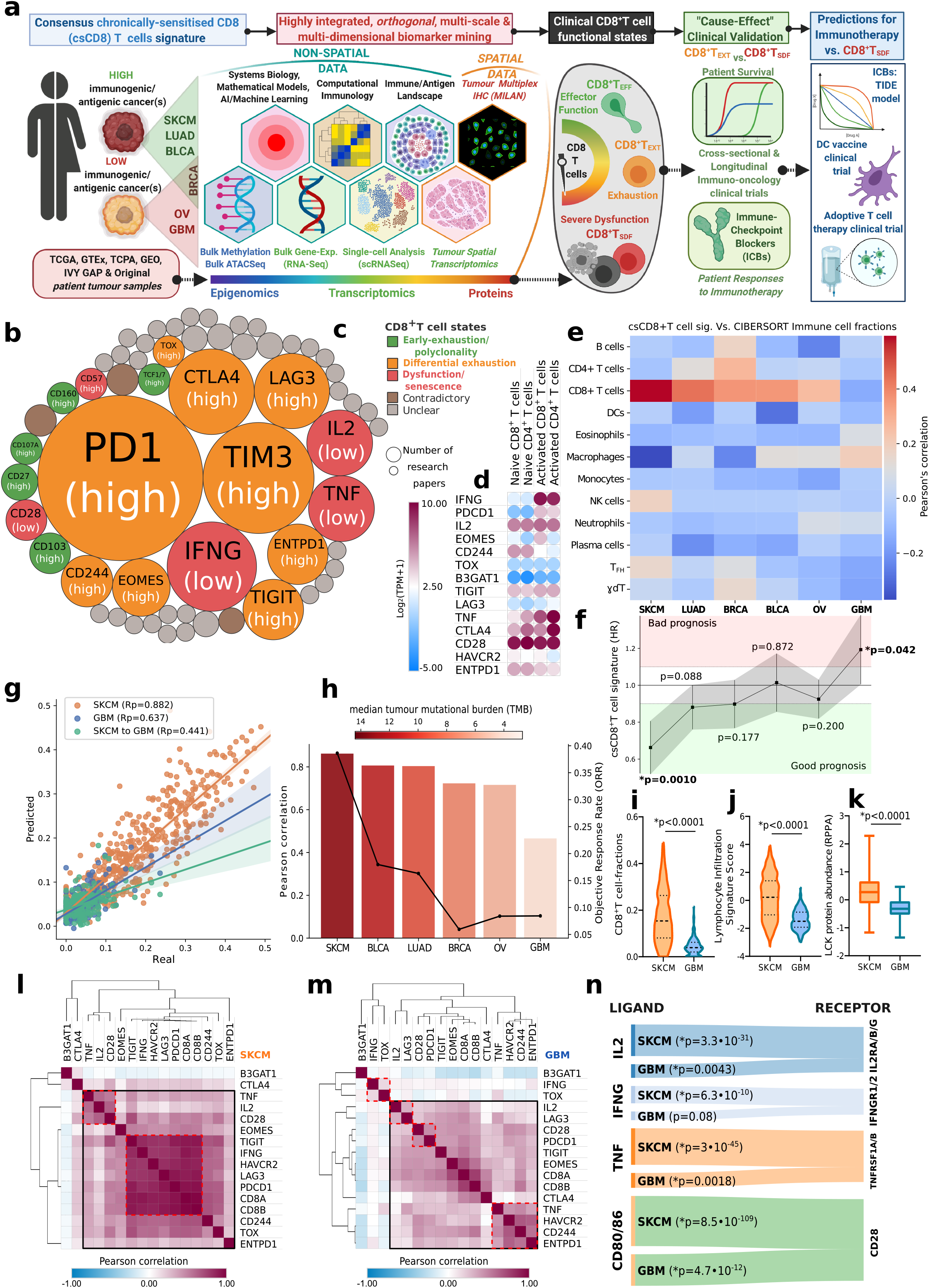
Tumour bulk-RNAseq-based immunogenomics. **(a)** Schematics of our large-scale mapping pipeline to differentiate, differentially exhausted tumoural-CD8^+^T_EXT_ from severely dysfunctional tumoural-CD8^+^T_SDF_; spanning >4000 patients across 7 cancer-types, integrating 24 distinct clinical-cohorts (both publicly available and in-house), with non-spatial (bulk/single-cell transcriptomic, epigenomic) or spatial-data (multiplex-protein, transcriptomic). Finally, we validated our main conclusions with immuno-oncology clinical trials’ data. For details, see “Study design” Methods-section. **(b,c)** Literature-wide meta-analysis of CD8^+^T cell-exhaustion/anergy biomarkers in cancer-patients. Circle-size indicates number of researchpapers per biomarker **(b);** colours indicate differential-association with CD8^+^T cell-states across studies (name of genes/proteins with contradictory/unclear associations aren’t mentioned) **(c). (d)** Dot-plot of indicated gene-expressions from human-peripheral CD4^+^/CD8^+^T-cells, untreated/naive, or *in-vitro* activated via CD3/CD28-dynabeads for 4h. Pearson’s correlation of csCD8^+^T cell-signature vs. 12 myeloid/lymphoid cell-fractions (CIBERSORT-derived) **(e),** or multi-variate CoxPh hazard-ratio/HR estimates for csCD8^+^T cell-signature’s prognostic-impact on overall patient survival (f) in The Cancer Genome-Atlas (TCGA)-datasets (BLCA, n=426; BRCA, n=1212; GBM, n=171; LUAD, n = 574; OV, n=427, SKCM, n=470). (g) Pearson-correlation for predicted values vs. real observations after TCGA-SKCM/GBM-specific XGBoost-models were trained (LOOCV evaluation). The TCGA-SKCM model was also tested on TCGA-GBM data (green) (line±95% confidence intervals; for statistics, see Fig.S1b). (h) Bar-plots indicating LOOCV-performance for csCD8^+^T cell-signature-trained ML-models across TCGA-datasets [trend-line: objective-response rates from anti-PD(L)1 immunotherapy clinical-trial metaanalysis; color-scale: median-tumour mutational-burden (TMB)]. (i-j) Violin-plots for indicated scores/fractions (CIBERSORT-derived) across TCGA-SKCM/GBM-datasets (Mann-Whitney test). **(k)** Box-plot for TCGA tumours-derived LCK protein-levels from The Cancer Proteome-Atlas (TCPA)-SKCM (n = 353) or TCPA-GBM (n=205) patient-datasets (median±S.D.; Mann-Whitney test). (l,m) Correlation-matrix for csCD8^+^T cell-signature across TCGA-SKCM/GBM-datasets. **(n)** Sankey-plots indicating Spearman-correlations between ligand-receptor genes across TCGA-SKCM/GBM-datasets. Statistical-significance threshold: *p<0.05. In **e,** DC: dendritic cells, NK: natural-killer, T_FH_: T follicular-helper, T_GD_: γδT cell.

Since every omics-approach presents specific limitations, we opted for a cross-omics method integrated via computational immunology and mathematical modelling. Herein, we first benchmarked the tumoural CD8^+^T cell-landscape via bulk-RNAseq based immunogenomics, and the predictions derived herewith were verified via analyses of single-cell CD8^+^T cell states/trajectories and spatial configurations. This enabled more precise interrogation of bulk-tumour transcriptomics data from cross-sectional and longitudinal immuno-oncology clinical-trials to deliver a “cause-effect” holistic validation, thereby providing a future roadmap for optimising immunotherapy against low antigenic/immunogenic tumours.

## RESULTS

### A continuum of functionally diverse CD8^+^T cell-states exist across human cancers

To construct an experimentally-validated CD8^+^T cell-signature **(Fig.1a),** we performed a metaanalysis of research studies on cancer patient-associated CD8^+^T cell-exhaustion/anergy **(Supplementary-Table.**1). Herein, we derived two features i.e., number of studies per CD8^+^T cell-marker **(Fig.1b;** indicated by circle-size), and the type of (broad) CD8^+^T cell-states observed per marker, per study **(Fig.1c;** indicated by colour-codes). This revealed that most studies used multiple-IRs and/or memory differentiation-markers to define exhaustion **(Fig.1b).** However, these markers were reported to simultaneously associate with polyclonality/early-exhaustion, differential exhaustion and senescence/dysfunction **(Fig.1c).**

This promiscuity was further outlined by transcriptomic-profiling of early stage, *in vitro-* activated, human peripheral-CD4^+^/CD8^+^T cells,^20^ exhibiting increase in multiple-IRs and memory differentiation-markers, along with effector-function markers (*IFNG, IL2, TNF*), despite no chronic tumour-exposure **(Fig.1d).** Importantly, some studies captured specific senescence/dysfunction features e.g., depleted effector-function markers **(Fig.1b).** Interrogation of experimental/perturbation datasets from ImmuneSigDB^21^ with genes linked to differential exhaustion or memory-differentiation (*PDCD1/PD1, CTLA4, HAVCR2/TIM3, ENTPD1/CD39, TIGIT, LAG3, TOX, EOMES, CD244*), effector-function (IFNG, *TNF, IL2*), costimulation (*CD28*), replicative senescence (B3GAT1/CD57) and CD8^+^T cell-lineage (*CD8A/CD8B*), simultaneously enriched for diverse T cell-specific features: activation/stimulation, effector/memory function, chronic viral-infection and IRs-linked exhaustion **(Fig.S1a).** Considering these genes’ wide-ranging coverage of differential T cell-functions/states, we exploited them to form a “chronically-sensitised” CD8^+^T (csCD8^+^T)-cell signature **(Supplementary-Table.2),** that can help in qualitative differentiation of CD8^+^T cellstates in multi-omics datasets.

Next, we pursued tumour-bulk RNAseq-driven immunogenomics centred around csCD8^+^T cell-signature within The Cancer Genome Atlas (TCGA)-dataset, to exploit its large-scale cohort-sizes and omics depth/diversity/standardization. To allow differential-mapping of CD8^+^T cellstates, we selected 6 TCGA-cancers exhibiting variable immunogenicity/antigenicity: highly immunogenic/antigenic (skin-cutaneous melanoma/SKCM, lung-adenocarcinoma/LUAD or bladder-carcinoma/BLCA), marginally immunogenic but low antigenic (breast-carcinoma/BRCA), and low immunogenic/antigenic (glioblastoma/GBM or ovarian-cancer/OV).^5,8,11^ Herein, we probed the correlation of csCD8^+^T cell-signature with 12 diverse myeloid/lymphoid immune-cell fractions, computed via CIBERSORT immune-deconvolution. Remarkably, this arranged the 6 TCGA-cancers as a “continuum” **(Fig.1e),** centred around a gradient of correlations between csCD8^+^T cell-signature and CD8^+^T cell-fractions, rather than other immune-cells, going from highly-positive (SKCM) to negative (GBM) **(Fig.1e).** Since csCD8^+^T cell-signature and CD8^+^T cell-fraction are supposed to mark the same transcriptomic entity (i.e., CD8^+^T cells), any discrepancy in their correlation possibly indicates transcriptional dysregulation in CD8^+^T cells. Indeed, operationally, this translated into a similar gradient of positive-to-negative prognostic impacts for csCD8^+^T cell-signature on patient’s overall survival (OS; multi-variate CoxPh-analyses) **(Fig.1f).**

These observations were intriguing from functional-transcriptome perspective; thus, we wondered whether machine learning (ML) algorithms trained to utilize this approach can further shed light on transcriptional-states of CD8^+^T cells. Indeed, our ensemble model successfully predicted CD8^+^T cell-fractions, using csCD8^+^T cell-signature, across SKCM-patients **(Fig.1g; Fig.S1b),** but its prediction-efficiency was significantly reduced across GBM-patients **(Fig.1g; Fig.S1b),** which kept reducing when models trained on the SKCM-patients were further tested on GBM-patients **(Fig.1g).** This suggested disparate feature weights between cancer-types; an indication of transcriptomic-dysregulation in GBM-CD8^+^T cells. Interestingly, ML-prediction performance also arranged the 6 TCGA-cancers as a continuum **(Fig.1h),** comparable with the gradient of tumour mutational-burden (TMB) (color-scale) and objective response-rates toward anti-PD(L)1-immunotherapy (trend-line),^8^ from an independent clinical-trial meta-analysis.^8^ Altogether, this emphasised a continuum of CD8^+^T cell-states across different tumour-types, compatible with their immunogenic/antigenic-profiles; with high (SKCM) and low (GBM) ICB-responsive cancers^5,8^ occupying the opposite-ends.

Owing to above phenotypes, we prioritised the SKCM vs. GBM comparison to “enrich” for the most prototypical versions of exhausted (but partially-functional) vs. (severely) dysfunctional CD8^+^T cell-populations, respectively; with periodic substantiating assessments in LUAD/BLCA/BRCA/OV. Consistent with these expectations, GBM exhibited diminished CD8^+^T cell-fractions **(Fig.1i),** and lymphocyte-infiltrates **(Fig.1j),** compared to SKCM. These trends weren’t transcriptome-specific, since GBM also exhibited reduction in lymphocyte-specific protein tyrosine-kinase, LCK^22^ (the only lymphocyte-specific protein within TCGA’s reverse-phase protein array) **(Fig.1k).** Curiously, the genes within csCD8^+^T cell-signature showed high inter-correlations across SKCM-tumours (i.e., “concordance”, a sign of stable association with a unified transcriptomic-entity) **(Fig.1l);** whereas they exhibited lower inter-correlations (“discordance”) across GBM **(Fig.1m).** Across GBM-tumours, discordance was especially apparent for effector-function genes (red dotted-boxes). Also, *IL2, IFNG, TNF* (but not *CD28-CD80/86*), correlated considerably less with their cognate receptor-coding genes in GBM **(Fig.1n).** Accordingly, genetic-signatures for IL2-signalling **(Fig.S1c),** IFNγ-linked GAS/ISRE-signalling **(Fig.S1d)** and TNF-induced NFkB-signalling **(Fig.S1e)** (but not CD28-signalling **(Fig.S1f))** were reduced in GBM. These results indicated a clear contrast in CD8^+^T cell-states between GBM and SKCM.

Lastly several, possibly ICB-nonresponding, SKCM-patients may enrich for late/terminal-CD8^+^T_EXT_.^2,14^ Accordingly, as CD8^+^T cell-enrichment decreased across patient-subgroups (from top-25% to bottom-25%), the gap between SKCM-patients and GBM-patients also decreased for the prognostic impact of CD8^+^T cell-fractions **(Fig.S1g).** Nevertheless, even amongst the least CD8^+^T cell-enriching patient-subgroup, SKCM had significantly higher OS-advantage over GBM, thereby emphasizing the distinct GBM-CD8^+^T cell-dysfunction **(Fig.S1g).** Overall, this indicates that GBM might enrich CD8^+^T_SDF_-ike immune-disparities, whereas SKCM enriches partially-functional CD8^+^T_EXT_ features.

### Organ-specific immunomodulatory niches “imprint” the tumoral lymphocytic continuum

Chronic viral-infection induced CD8^+^T_EXT_ adopt organ-specific effector-functions.^23^ Considering the above continuum, we wondered whether distinct organ-of-origin/localization for above cancers influenced lymphocytic effector/co-stimulation markers. To address this, we procured bulk-RNAseq data for normal-organ/tissue counterparts for above cancer-types from Genotype-Tissue Expression (GTEx)-dataset.^24,25^ Interestingly, an intriguing continuum of CD8^+^T cell-fractions and *IFNG-IL2-CD28* indeed existed across different normal-organs, ranging from high/medium (lung/breast>bladder/skin) to low-enrichment (ovary/brain) **(Fig.2a-c; Fig.S2a).** Since brain/ovary are “immune-privileged” with proficient immunotolerance,^26,27^ we wondered how this aligned with GBM/OV. We compared TCGA data from primary-tumours or metastases (wherever available) vs. GTEx normal-organ counterparts. Fascinatingly, the primary tumour-samples largely reflected the trends for their corresponding normal-tissue’s CD8^+^T cell-fractions, *IFNG* or *IL2*; albeit at somewhat lower intensities, possibly resulting from malignant transformation **(Fig.2a-c; Fig.S2a).** Herein, primary-SKCM showed increased CD8^+^T cell-fractions and *IFNG-CD28,* beyond normal-skin levels **(Fig.2a,b).** Notably, while melanocyte content of benign nevi resembles SKCM better than normal-skin, yet, expression-patterns for primary-SKCM vs. benign nevi comparison resembled primary-SKCM vs. skin **(Fig.2b,c; Fig.S2a,b).** Prominently, brain::GBM or ovary::OV combinations showed higher sparsity of CD8^+^T cell-fractions and *IL2* **(Fig.2a-c).** Thus, organ-specific niches can imprint differential CD8^+^T cell-enrichment across human cancers.

**Figure 2.**
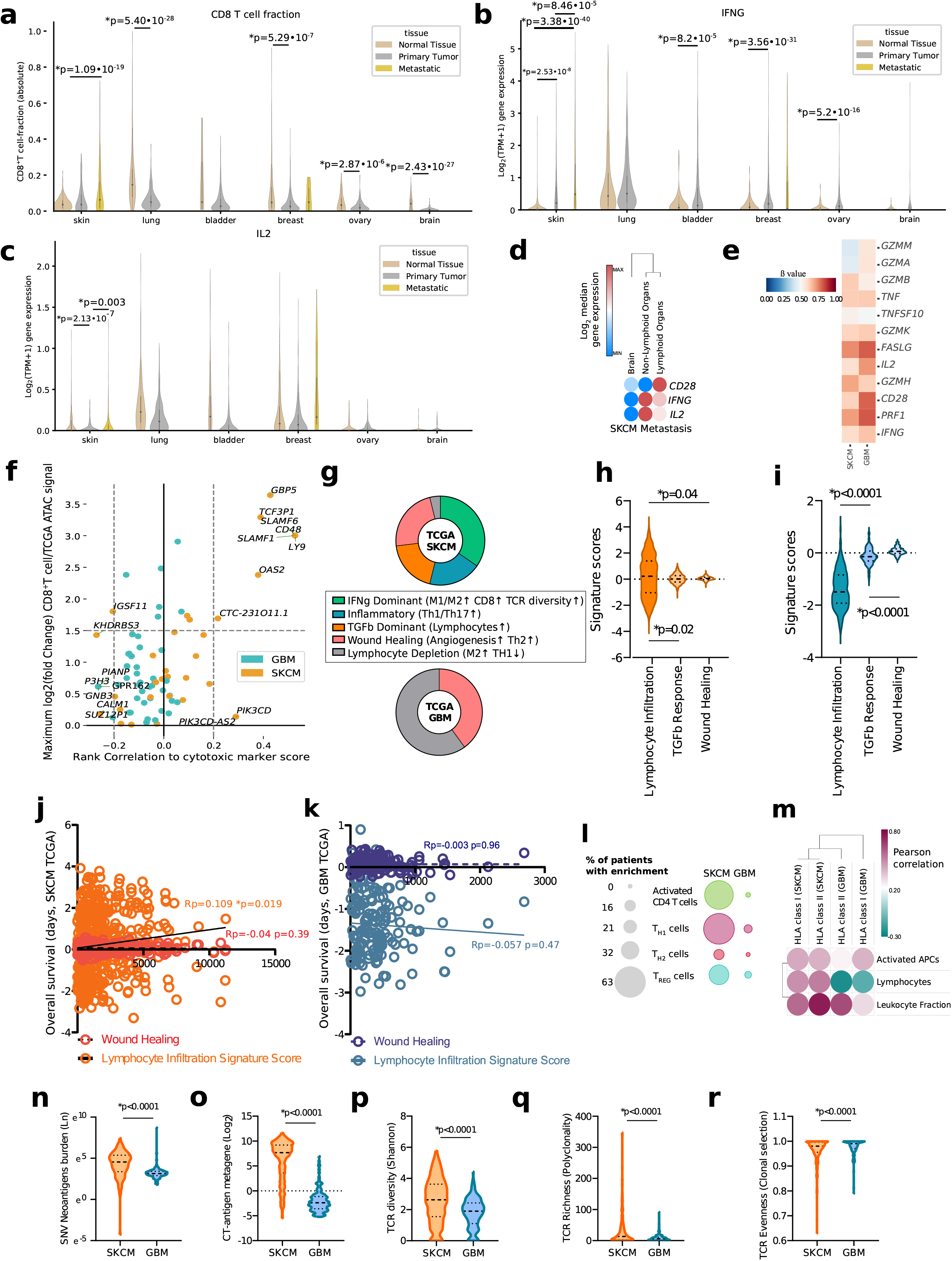
Comparative-immunogenomics, epigenomics and tumour/antigenic landscape-mapping. Violin-plots comparing CD8^+^T cell-fractions **(a)** (CIBERSORT-derived), or *IFNG* **(b),** *IL2* **(c)** expression between TCGA primary tumour/metastasis bulk-RNAseq vs. Genotype-Tissue Expression (GTEx) normal organ/tissue bulk-RNAseq datasets (Welch’s t-test). Brain (normal-tissue, n=207; primary-tumour, n=153), bladder (normal-tissue, n=9; primary-tumour, n=407), breast (normal-tissue, n = 179; primary-tumour, n = 1092; metastatic-tissue, n=7), lung (normal-tissue, n=288; primary-tumour, n=513), ovary (normal-tissue, n=88; primary-tumour, n=419), skin (normal-tissue, n=557; primary-tumour, n = 102; metastatic-tissue, n=366). (d) Dot-plot of indicated genes’ expressions in SKCM-metastases within lymphoid (n=18), non-lymphoid (n=22) organs, or brain (n=29). **(e)** Heatmap for DNA-methylation patterns (β-value), of indicated genes in TCGA-SKCM (n=475) or TCGA-GBM (n=115). (f) Dot-plot showing ATACseq peak-to-gene links predicted to associate with CD8^+^T cells within TCGA-SKCM (n=13) or TCGA-GBM (n=9) datasets. Each dot represents an individual predicted-gene peak, further Pearson-correlation to cytolytic-marker score (*GZMA/PRF1-*based). GO-BP enrichment was done for top gene-hits (>1.5 log_2_fold-change; >0.2 Pearson’s-correlation). **(g)** Extracellular network-analyses using interactions between CD8^+^T cell-fractions and *IFNG-IL2-TNF-CD28* to enrich tumour immune-landscape features across TCGA-SKCM/GBM-datasets. **(h-i)** Violin-plots of indicated signature-score’s levels in TCGA-SKCM (h) or TCGA-GBM (i) (Repeated measures ANOVA with Holm-Sidak’s multiple comparisons test). (j,k) Pearson’s correlation between indicated signature-scores and patient-overall survival (OS, in days) in the TCGA-SKCM (**j**) and TCGA-GBM (**k**) patient-cohorts. (**l**) GSEA for indicated CD4^+^T cell-signatures in TCGA-SKCM/GBM-patients (g-value<0.05). (m) Pearson-correlation between HLA class-I/II signatures and indicated immune-fractions for TCGA-SKCM/GBM. Violin-plots for singlenucleotide variation (SNV)-derived neoantigens **(n),** cancer-testis (CT)-antigen signature **(o),** T cell-receptor (TCR) Shannon-diversity/richness/evenness metrics **(p-r)** in TCGA-SKCM/GBM datasets (Mann-Whitney test). Statistical-significance threshold: *p<0.05. For different gene-signatures, see Supplementary-Table.2.

Notably, there were prominent differences between primary-SKCM and metastatic-SKCM, with latter showing higher pro-lymphocytic orientation **(Fig.2a-c).** SKCM frequently metastasises far from skin-locales, toward distant lymphoid/non-lymphoid niches including brain.^28^ Thus, we wondered how effector-function was shaped within SKCM brain-metastasis. Interestingly, in an independent cohort (since, TCGA-dataset lacks intra-cranial samples),^29^ the effector-function genes showed relatively lower expression in SKCM brain-metastasis, compared to extra-cranial SKCM-metastasis **(Fig.2d).** This suggested that localization within immuno-tolerant/privileged organs may dictate lymphocytic effector-function of otherwise pro-lymphocytic, melanocytes-derived, SKCM. To further substantiate this, we performed a comparison with primary uveal melanoma/UVM, a melanocytes-derived malignancy located in eyes, another immune-privileged organ. Remarkably UVM resembled GBM, rather than SKCM, in exhibiting depletion of CD8^+^T cell-fractions or *IFNG-IL2-CD28* **(Fig.S2c,d).** Thus, some tumours align with organ-specific immunotolerant niches in reducing CD8^+^T cells and lymphocytic effector-functions.

### Epigenetic defects in effector/cytolytic modules distinguish CD8^+^T_SDF_-enriching tumour

Transcriptional and epigenetic dysregulations together cause permanent suppression of lymphocytic effector/cytolytic functions.^3,12,16^ Considering GBM-associated lymphocytic disparities, we pursued epigenomics analyses (DNA-methylation) within TCGA-datasets. Interestingly, GBM enriched for higher DNA-methylation (vs. SKCM), of effector-function (*IL2*-*CD28*) and cytolytic activity (*PRF1, FASLG-GZMM-GZMA*) genetic-modules **(Fig.2e),** which further associated with their transcriptional down-regulation **(Fig.S3a).** Since, DNA-methylation reduces transcription by limiting chromatin-accessibility;^13^ we wondered if above results indicated restricted accessibility for GBM-CD8^+^T cells-associated effector/activation genetic modules. We addressed this via TCGA-derived, assay for transposase-accessible chromatin using sequencing (ATACseg)-data.^30^ Strikingly, across the entire TCGA ATACseq-dataset, only for SKCM-patients (but not GBM) there were above-threshold CD8^+^T cells-linked genes with increased chromatin-accessibility and positive-correlation with cytolytic-activity score **(Fig.2f).** These genes were associated with IFN-response (*OAS2-GBP5*) and lymphocyte effector function-modulation (*SLAMF6-SLAMF1-LY9-CD48*) **(Fig.2f).** Altogether, this indicated epigenetic-dysregulation in GBM-CD8^+^T_SDF_, a sign of putative permanent-dysfunction.^10^

### Defects in antigen-priming and CD4^+^T cell enrichment distinguish the immunosuppressive CD8^+^T_SDF_-landscape

There are CD8^+^T cell-autonomous (chronic/suboptimal antigen-priming) and non-autonomous reasons (defective CD4^+^T cell-compartment and T cell-suppressive tumour micro-environment) behind CD8^+^T_EXT_/T_SDF_-features.^10,31^ To better understand this, we pursued tumour immune-landscape mapping.

A pan-cancer immunogenomic analyses (> 10,000 patients across 33 TCGA-cancers) had identified six broad tumour immune-landscapes i.e., wound healing, IFNγ-dominant, inflammatory, lymphocyte-depleted, immunologically-quiet, or TGFβ-dominant.^25^ We pursued an extracellular network-analysis,^25^ to decipher which of these immune-landscapes were enriched during effector-function genes’ (*TNF-IL2-IFNG-CD28*) “computational interactions” with CD8^+^T cell-fractions, within TCGA-SKCM/GBM datasets. Most SKCM-tumours had IFNγ-dominant or inflammatory landscapes, followed by those with immuno-suppressive TGFβ-response/wound healing **(Fig.2g; Fig.S3b)**.^25^ However, GBM-patients exclusively enriched for lymphocyte-depletion or wound healing **(Fig.2g; Fig.S3b)**.^25^ Accordingly, lymphocyte-infiltration significantly exceeded TGFβ-response/wound healing in SKCM **(Fig.2h).** While, wound healing or TGFβ-response dominated GBM-tumours, that were devoid of lymphocyte-infiltrates **(Fig.2i).**

Lymphocyte depletion and wound healing accentuate tumoural-macrophage infiltrates with primarily immuno-suppressive (M2)-orientation.^25,32^ Accordingly, GBM enriched more M2 macrophages than SKCM **(Fig.S3c).**^32^ Operationally, in SKCM, lymphocyte-infiltration (but not wound healing) positively correlated with patient-OS **(Fig.2j)** whereas in GBM, both showed null-to-negative correlation **(Fig.2k).** The lymphocyte-depleted GBM made us curious about the status of CD4-helper T (T_H_)-cells, since CD4^+^::CD8^+^T cell-crosstalk helps avoid CD8^+^T cell-dysfunction.^2,10,31^ Strikingly, compared to SKCM, GBM-tumours: (I) poorly enriched for all major T_H_-subsets (T_H1_/T_H2_/T_REG_; estimated via gene-set enrichment analyses/GSEA using established pan-TCGA signatures^33^) **(Fig.2l),** and (II) lacked co-clustering between CD8^+^T cells and other anticancer lymphocyte-fractions (T_H1_, activated-NK cells, memory-B cells, activated-memory T_H_), or activated (M1)-macrophages **(Fig.S4a,b).**

Severe dysfunction of CD4^+^/CD8^+^T cell-compartments within GBM, made us wonder whether we were dealing with “canonical” (chronic antigen-stimulation induced pre→early→late exhaustion) or “non-canonical” CD8^+^T cell-states (suboptimal antigen-priming facilitated dysfunction).^14,31,34^ Hence, we analysed the antigenicity of TCGA-GBM, vs. TCGA-SKCM, on the levels of antigen-enrichment, antigen-presenting molecules (HLA-family), and T-cell receptor (TCR) diversity-metrics. GBM exhibited inconsistent to negative correlations across HLA-I/II genes, vs. activated antigen-presenting cell (APC)-fractions, lymphocyte-fractions or leukocyte-fractions **(Fig.2m);** paralleled by diminished SNV-neoantigens and reduced cancer-testis (CT)-antigen’s gene-expression **(Fig.2n-o).** This associated with severe reduction in TCR-diversity within GBM (vs. SKCM) **(Fig.2p),** in terms of both polyclonality (low TCR-richness=low polyclonality) **(Fig.2q)** and clonal selection (higher TCR-evenness=lower clonal-selection) **(Fig.2r).** Interestingly, in SKCM, SNV-neoantigens (but not CT-antigens) positively correlated with HLA-I genes/TCR-richness **(Fig.S4c,d)** whereas no such correlations existed in GBM **(Fig.S4e,f).** Thus, CD8^+^T_SDF_ might be products of suboptimal antigen-priming (in absence of CD4^+^T cells);^31^ embedded within an immuno-suppressive wound healing/TGFβ-enriched^34^ landscape. Contrastingly, CD8^+^T_EXT_ are products of chronic antigen-stimulation based exhaustion. These observations predicted contrasting cell-states/trajectories for CD8^+^T_EXT_/T_SDF_ due to distinct developmental cues.

### SKCM and GBM CD4^+^/CD8^+^T cell-populations exhibit highly contrasting single-cell states

We wanted to confirm our immunogenomics-derived predictions at single-cell resolution for CD4^+^/CD8^+^T cells. Hence, we shortlisted existing human pan-tumour GBM/SKCM single-cell RNAseq (scRNAseq)-maps with lymphocytes data (6xSKCM, 4xGBM; **Fig.S5a)**;^35^ and further selected high-quality datasets on the basis of qualitative (high gene-capture for csCD8^+^T cell-signature within CD8^+^T single-cells; **Fig.S5a)** and quantitative criteria (representativeness of T cell-counts; **Fig.S5b).** Qualitatively, Smart-Seq2 tumour-datasets (3xSKCM, 1xGBM) had the highest gene-capture **(Fig.S5a).** Between these, a sizable difference in CD8^+^T cell-counts (SKCM>>GBM) existed. To allow representative comparison, we selected SKCM/GBM dataset-pairs based on how closely SKCM-to-GBM ratios (forT cell-counts), per scRNAseq dataset-pair, overlapped with similar ratios for TCGA-derived lymphocyte/CD8^+^T cell/T_REG_ CIBERSORT-fractions **(Fig.S5b).** Through these criteria, we short-listed GBM **(Fig.S5c)**^36^ and SKCM **(Fig.S5d)**^37^ dataset-pairs with high gene-capture depth (>9000-genes) per CD4^+^/CD8^+^T cells **(Fig.S5e).**

GBM/SKCM-CD8^+^T cells enriched for csCD8^+^T cell-signature **(Fig.3a-b);** but SKCM-CD8^+^T cells exhibited higher co-expression of the signature’s genes per cell (thus better co-clustering, causing higher nodal-distance from centre), than GBM (more “fragmented” clusters, thus shorter nodal-distance from centre) **(Fig.3c).** Altogether, SKCM-T cells better co-expressed effector/cytolytic-function genes relative to *CD8A/CD8B,* as indicated by the larger radar-plot area (vs. GBM) **(Fig.3d; Fig.S5f-g).**

**Figure 3.**
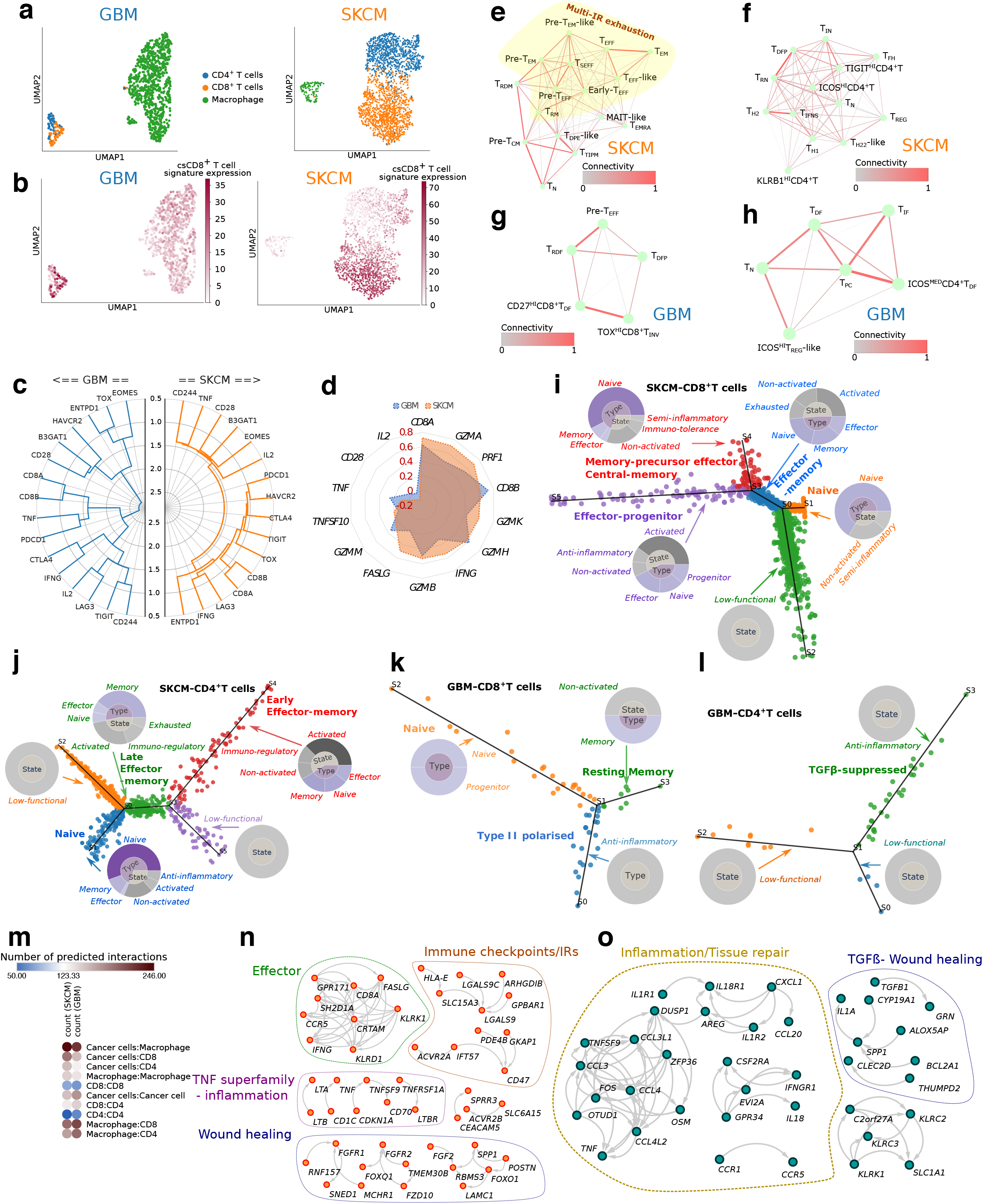
Single-cell (sc)RNAseq-analyses of tumour-tissue derived cancer or immune cells. **(a,b)** UMAP-visualizations of indicated immune cell-types, based on CIBERSORT-LM22 gene-signatures, in SKCM (19-patients: 126 macrophages, 2069 T-cells) or GBM (28-patients: 754 macrophages, 94 T cells) patient-cohorts **(a).** These immune cells were furthered coloured for csCD8^+^T cell-signature expression-levels **(b). (c)** Tree-wise comparison of csCD8^+^T cell-signature’s co-clustering tendency across GBM/SKCM-CD8^+^T single-cells (closeness of dendrogram nodes to the outer-edge indicates higher correlations between indicated genes per T-cell). **(d)** Nearest-neighbour (spearman’s rank-correlation) analyses of indicated genes across all SKCM/GBM-T single-cells (z-score row centring), **(e-h)** Partition-based graph abstraction (PAGA)-derived T single cell-subpopulation connectivity-networks for SKCM-CD8^+^T cells **(e),** SKCM-CD4^+^T cells **(f),** GBM-CD8^+^T cells **(g),** GBM-CD4^+^T cells **(h).** T-cells expressing multiple-IRs based general-exhaustion are demarcated **(e).** Herein: T_EM_, effector-memory; T_EFF_, effector; T_RM_, resting-memory; T_CM_, central-memory; T_EMRA_, effector-memory re-expressing CD45RA; T_SEFF_, suppressed-effector; T_DPE_, double-positive effector; MAIT, mucosal-associated invariant-T; T_N_, naive; T-_TIPM_, tumour-infiltrating peripheral-memory; T_REG_, regulatory; T_RN_, resting-naive; T_FH_, follicular-helper; T_IFNS_, interferon-stimulated; T_H2_, type-II; T_H1_, type-I; T_H22_, type-22; T_DFP_, dysfunctional-precursor; T_IN_, inflamed-naive; T_DF_, dysfunctional; T_IF_, inflammatory; Tpc, precursor; T_RDF_, resident-dysfunctional; T_INV_, invariant, **(i-l)** Single-cell STREAM-based trajectories annotated via our FIAT-approach for SKCM-CD8^+^T cells **(i),** SKCM-CD4^+^T cells **(j),** GBM-CD8^+^T cells (k) orGBM-CD4^+^T cells (I). Doughnut-plots represent the branch-level FIAT-annotations derived via branch-specific GSEA to enrich ImmuneSigDB-datasets for final-annotation (Supplementary-Table.3) of respective T cell’s functional-phenotypes/states. **(m)** Dotplot depicting CellPhoneDB-computed number of unique interactions between indicated cell-type combinations in GBM/SKCM-scRNAseq datasets, **(n,o)** Network topology-analyses based prioritisation of SKCM/GBM scRNAseq-based CellPhoneDB interaction-predictions (Fig.S7a-d) within TCGA-SKCM/GBM tumour bulk-RNAseq network-context (threshold: FDR adjusted p-value<0.05).

Comprehensive T cell-subpopulation annotation (see **Supplementary-Box.1** for details) coupled with trajectory networks, revealed that SKCM enriches diverse CD4^+^/CD8^+^T cell-subpopulations with full-spectra of distinct functional-states **(Fig.3e-f; Fig.S6a-b).** These subpopulations exhibited dense interconnectivity between naive, resident/central memory, pre-effector/pre-memory, effector/effector-memory, and anti-inflammatory/immunoregulatory transcriptional-states;^2^ majority of which showed multiple-IR based exhaustion-characteristics. Herein, effector/effector-memory CD8^+^T cells (T_EFF_/T_EM_) exhibited CD8^+^T_EXT_-like transcriptional-features **(Fig.3e; Fig.S6a),** suggestive of an ICB-responsive population.^2,10^ Likewise, SKCM-CD4^+^T cells showed diverse polarization-states: type-I (T_H1_), type-II (T_H2_), immunoregulatory (T_REG_), or IFN-inflamed/inflammatory **(Fig.3f).** In-contrast, GBM exclusively enriched for sparsely interconnected dysfunctional (T_DF_/T_RDF_/T_DEP_), or immunosuppressive/invariant (*TOX*^HI^CD8^+^T_INV_, /COS^HI^T_REG_-like) CD4^+^/CD8^+^T cell-subsets **(Fig.3g-h; Fig.S6c-d);** thus, representing CD8^+^T_SDF_ population. Anticipatingly, GBM-CD8^+^T cells didn’t show typical characteristics of chronic antigen-stimulation, unlike SKCM-CD8^+^T cells e.g., simultaneous high expression/population-coverage for multiple-IRs (*PDCD1, HAVCR2, CTLA4, ENTPD1, TIGIT, LAG3*)^10^ and detectable terminal differentiation-marker (*EOMES*)^38^ **(Fig.3e-h; Fig.S6a-d).** Altogether, these analyses confirm our immunogenomics-driven conclusions at single-cell resolution, establishing functional-proficiency of SKCM-CD8^+^T_EXT_/CD4^+^T cells and severe-dysfunction in GBM-CD8^+^T_SDF_/CD4^+^T cells.

### CD8^+^T_SDF_ exhibit dysfunctional single-cell trajectories lacking effector/memory phenotypes

Above cell state-trajectories indicated distinct, non-overlapping, developmental-patterns for CD8^+^T_EXT_/T_SDF_. To confirm this, we pursued pseudotime-trajectory analyses aimed at deciphering functional immune-annotations. T cell-annotations for standard pseudotime-trajectories don’t align properly with experimental/mechanistic evidence.^2^ Therefore, we used a customised “functional immune-annotation for trajectories (FIAT)” approach. We first used STREAM^39^ to generate standard CD4^+^/CD8^+^T cell-trajectories. Thereafter, rather than annotating trajectory-branches via few distinguishing genes, we took an unbiased approach for annotation through GSEA per trajectory-branch, enriching for ImmuneSigDB’s^21^ experimental-datasets on immune-phenotypes/states. Herein, we depicted the STREAM treeplots **(Fig.3i-l),** and adjacent to each trajectory-branch, we placed doughnut-plots portraying the topmost lymphocytic phenotype (blue-color), or state (grey-color) associated ImmuneSigDB-datasets enriched by that branch (see **Supplementary-Table.3** for full gene/dataset-lists). These phenotypes/states helped define the final functional immune-annotations per trajectory-branch (see, **Methods** for details).

SKCM-CD47CD8^+^T cells presented typical effector/memory trajectories **(Fig.3i-j).**^12^ For CD8^+^T cells, naive-trajectory connected with effector-memory trajectory showing partial activation/exhaustion **(Fig.3i),** which further connected to: (I) effector-progenitor trajectory^12,40^ (enriching *TCF3/7,* multiple-IRs) showing partial-activation **(Fig.S6e);** and, (II) memory precursor-effector like (immuno-regulatory) trajectory,^41^ also exhibiting mixed central-memory (*SELL^HIGH^CD27^HIGH^IL7R^HIGH^*) features **(Fig.S6f; Fig.3i)**^42^ This was complemented by CD4^+^T cells showing early (activated-immunoregulatory) to late (activated-immunoregulatory-exhausted) effector-memory trajectories **(Fig.3j).** There were also some low-functional CD4^+^/CD8^+^T cells, possibly representing late-exhausted trajectory, co-existing with above pre/early-exhausted trajectories **(Fig.3i-j).** These patterns resemble consensus T cell-differentiation during chronic antigen-stimulation induced exhaustion.^2,12^

Conversely, GBM-CD4^+^/CD8^+^T cells displayed exclusively low-functional or anti-inflammatory trajectories **(Fig.3k),** and TGFβ-suppression (which facilitates T cell-quiescence)^43^ **(Fig.3l).** Although the number of GBM-CD4^+^/CD8^+^T cells were limited, this didn’t compromise our trajectory-analyses since, we had significant gene-depth (owing to Smart-Seq2) to create statistically-stable trajectories (see **Supplementary-Table.3).** Importantly, there were no significant differences in gene drop-outs, for the top 3566 most-variable genes shared by SKCM/GBM CD8^+^T cells. Thus, low GBM-CD8^+^T cell-counts didn’t create gene-sparsity issues that can destabilize trajectory-plotting. Moreover, our single-cell and immunogenomic estimations were compatible. In conclusion, single-cell trajectories of SKCM-CD8^+^T_EXT_ completely contrasted from GBM-CD8^+^T_SDF_, thereby suggesting divergent, tumour-specific, origins of CD8^+^T cell-dysfunction.

### TGFβ/wound healing and tissue repair-inflammation define T cell-cancer cell interactions in CD8^+^T_SDF_-enriching tumour

To substantiate TCGA immune-landscaping on single-cell level, we explored computational-predictions of receptor-ligand interactions between cancer cells-CD4^+^/CD8^+^T cells using CellPhoneDB.^44^ Quantitatively, CD4^+^/CD8^+^T cells had higher predicted interaction-counts with SKCM-cancer cells, than GBM-cancer cells **(Fig.3m).** Qualitatively, several cancer-type specific and overlapping/shared receptor-ligand interactions were predicted between T cells and SKCM/GBM-cancer cells **(Fig.S7a-d).** However, for context-driven prioritisation, we filtered the predicted CD8^+^::CD4^+^T-cell and CD4^+^/CD8^+^T-cell::cancer-cell interactions via SKCM/GBM TCGA-based networking-approach. SKCM-associated interactions signified co-existence of effector-signalling, IRs, TNF superfamily-associated inflammation and wound healing **(Fig.3n).** Conversely, in GBM, these interactions were skewed toward tissue repair-inflammation and TGFβ/wound healing responses **(Fig.3o).** Thus, tissue repair/healing-like responses dominated CD47^+^CD8^+^T-cell::cancer-cell interactions in the CD8^+^T_SDF_-context whereas CD8^+^T_EXT_-context exhibited effector/immuno-regulatory interactions.

### CD8^+^T_SDF_ exhibit IL2-withdrawal linked cell-cycling defects

GBM-CD8^+^T_SDF_ trajectory suggested cell biological anomalies. Usually IL2-signalling drives cell-cycling of antigen-primed pre/early-CD8^+^T_EXT_.^45^ CD8^+^T cells with suboptimal TCR-signalling are particularly dependent on IL2-stimulation.^46^ Since *IL2* was depleted in brain, and GBM exhibited low-antigenicity, we wondered if this led to cell-cycling defects in CD8^+^T_SDF_.

Cell-cycle (phase) scores computed from scRNAseq-data revealed that GBM-CD8^+^T cells exhibit significantly lower S/G2M-scores and higher G1-score (vs. SKCM-CD8^+^T cells) **(Fig.4a-b).** GBM-CD4^+^T cells exhibited relatively better cell-cycle scores albeit with similar “substandard” patterns **(Fig.4a-b).** Within IFNγ/IL2-signalling signatures **(Supplementary-Table.2),** JAK3 (downstream of IL2/γc-family cytokines) and STAT1 (downstream of type-I/II IFNs) are well-established positive-regulators of T cel I-cycling.^10,47,48^ Remarkably, GBM-CD8^+^/CD4^+^T cells showed significant *JAK3/STAT1* depletion across cell-cycle phases **(Fig.4c-f; Fig.S8a-d).** *JAK3* disparity appeared more severe than *STAT1* in GBM-CD8^+^T cells thereby highlighting IL2-dysregulation **(Fig.4e-f).** T cells undergoing cell-cycle exit induced by IL2-withdrawal (before apoptosis/quiescence), exhibit a specific genetic-signature, as published previously.^49^ Interestingly, GBM-CD8^+^/CD4^+^T cells showed higher association with this signature,^49^ than SKCM-CD8^+^/CD4^+^T cells **(Fig.4g).** This indicated G1 arrest-like phenotype in CD8^+^T_SDF_.

**Figure 4.**
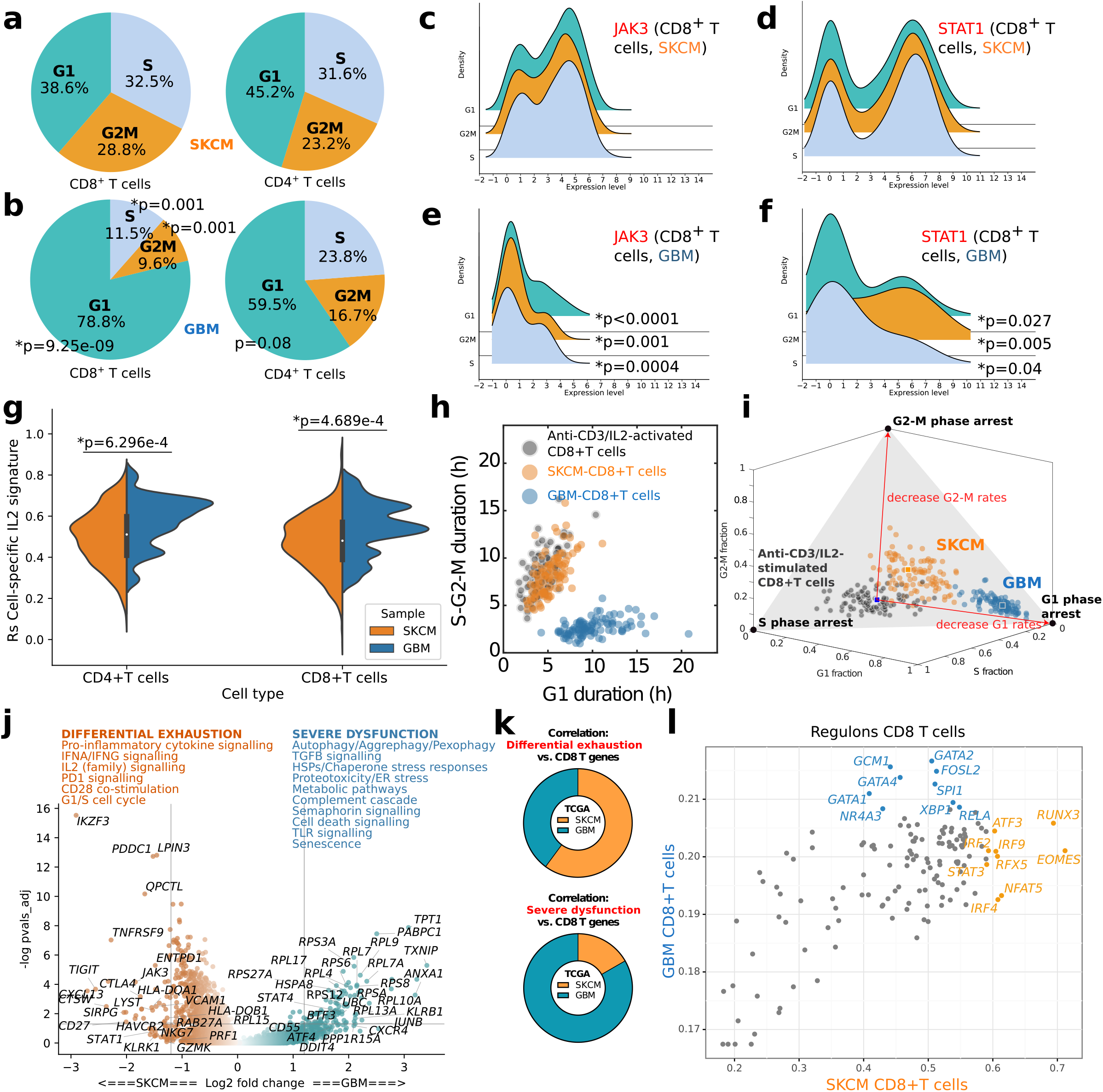
The scRNAseq-driven mapping of tumoral T cell-cycle and differential exhaustion-states in melanoma and glioblastoma. **(a,b)** Pie-charts representing the distribution of CD8^+^/CD4^+^T single-cells per cell-cycle phase scores (i.e., G1, S or G2M) for SKCM **(a)** or GBM **(b)** tumoral scRNAseq-datasets (Fisher-exact test-based statistics was estimated for SKCM vs. GBM comparison per cell-cycle phase score), **(c-f)** Ridge-plots illustrating the distribution of *JAK3/STAT1* expression in SKCM-CD8^+^T single-cells **(c,d)** and GBM-CD8^+^T single-cells **(e,f)** (statistics was estimated for SKCM vs. GBM comparison, for each gene per cell-cycle phase score; area-under-the-curve method combined with ANCOVA-analyses). **(g)** Double-sided violin-plot comparing the Spearman Rho-value distribution (Y-axis) for each single-cell in GBM/SKCM T-cell populations (X-axis) to the T-cell specific IL2-withdrawal genetic-signature (independent t-test). Distribution of cell-cycle phase pseudo-durations for 100 CD8^+^T cell-cycle iterations using stretched cell-cycle model **(h)** or Erlang cellcycle model (i). Red-arrows (i) indicate underlying mathematical reaction-rates corresponding to each (adjustable) cell-cycle phase pseudo-durations, **(j)** Volcano-plot comparing differential gene-expression between GBM/SKCM CD8^+^T-cells. Log_2_fold-change between both cancertypes is plotted on the X-axis while Benjamini Hochberg (BH)-adjusted p-values occupy Y-axis. Major/redundant REACTOME-pathways enriched for each cancer-type are highlighted (for full-analyses, see Fig.S8e). (k) Correlation of Fig.4j-derived “differential-exhaustion” or “severe-dysfunction” signatures with CD8^+^T genes (*CD8A/CD8B*) within TCGA-SKCM/GBM tumour bulk-RNAseq-datasets. **(l,m)** Scatterplot comparing the regulon-specificity scores for GBM/SKCM-CD8^+^T-cells. Statistical-significance threshold was *p<0.05. For different gene-signatures, see Supplementary-Table.2.

To further explore this, we used cell-cycle mathematical modelling for CD8^+^T cells proliferating after anti-CD3 (TCR-stimulus) and IL2 stimuli.^50^ We adjusted this model’s parameters to reflect the cell-cycle scores computed from CD8^+^T cells’ scRNAseq-data **(Fig.4a-b),** and used experimentally-verified kinetics of anti-CD3/IL2-stimulated CD8^+^T cells as reference-point for normal activated-T cell-cycle.^50^ Indeed, SKCM-CD8^+^T cells co-clustered closely with this reference-point in showing high S/G2M-cycling whereas GBM-CD8^+^T cells failed to co-cluster and showed higher G1-persistence **(Fig.4h).** Using another cell-cycle model that allows better appraisal of cell phase-distributions,^51^ we observed that changing G1 phase-specific reaction-rates specifically predisposed GBM-CD8^+^T cells toward G1-arrest **(Fig.4i).** Thus, CD8^+^T_SDF_ suffer from IL2-withdrawal induced cell-cycle defects, exemplifying their suboptimal antigen-priming and localization in immuno-tolerant niche.

### Maladaptive pro-death stress responses and immunosuppressive gene-regulatory networks distinguish CD8^+^T_SDF_

Next, we decided to decipher specific cell biological characteristics (not biased by our signature-based approaches), differentially enriched between SKCM-CD8^+^T_EXT_ vs. GBM-CD8^+^T_SDF_.

Volcano-plots coupled to REACTOME pathway-enrichment on scRNAseq-data **(Fig.4j; Fig.S8e)** indicated that CD8^+^T_EXT_ enriched for effector-function (IFNγ/CD28/IL2-signalling), cell-cycling and pro-inflammatory/PD1-signalling. Contrastingly, CD8^+^T_SDF_ enriched for ribosome biogenesis-stress (*RPL/RPS* genes),^52^ maladaptive stress-responses (proteotoxicity/ER-stress, chaperone-stress, autophagy, senescence), pro-death signalling and tissue-repair/immunotolerance (TGFβ/Complement/Semaphorin-signalling).^2,10,12,14^ CD4^+^T cells showed similar patterns **(Fig.S8f-g).** Interestingly, genetic-signatures of “exhaustion” or “severe-dysfunction”, based on major differentially-expressed genes in CD8^+^T_EXT_/T_SDF_ volcanoplots **(Fig.4j),** showed (contrasting) higher positive-correlations with *CD8A/B* in TCGA-SKCM or TCGA-GBM, respectively **(Fig.4k).** This confirms SKCM/GBM as prototypes for CD8^+^T_EXT_/T_SDF_.

Next, we mapped differently-enriched, transcription factors-driven, gene-regulatory networks (GRNs) between SKCM-CD8^+^T_EXT_ (orange-dots) and GBM-CD8^+^T_SDF_ (blue-dots), in scRNAseq-data **(Fig.4l)** with SCENIC. Again, CD8^+^T_EXT_ enriched for effector-function (*STAT3, IRF2/4/9*)^9^ memory-differentiation (*RUNX3, EOMES*),^2^ or antigen-stimulation (*NFAT5, ATF3*) GRNs;^10^ but CD8^+^T_SDF_ enriched for T cell-suppression (*GATA1/2/4, NR4A3*),^53^ exhaustion-modulation (*FOSL2*)^54^ and ER-stress (*XBP1*)^55^ **(Fig.4l).** Altogether, divergent from CD8^+^T_EXT_, CD8^+^T_SDF_ exhibit fatal maladaptive-stress responses and immunosuppressive GRNs thereby providing “mechanistic view” for lymphocyte-depletion in GBM.

### Spatial gradient of CD8^+^T cell exhaustion radiates from tumoural blood vessels

Most tumoural-CD8^+^T cell studies are non-spatial analyses.^2,14^ Thus, spatial configurations of CD8^+^T_EXT_/T_SDF_ remain unaddressed. Henceforth, we pursued a two-step approach: we started with protein-level, discovery-analyses, at single-cell resolution via multiplex immunohistochemical-probing of tumour-tissue (MILAN-method)^19^ in our own SKCM/GBM patient-cohorts. This represented our own original and novel observations, which we further (independently) validated via IVY-GAP,^56^ a GBM tissue-anatomy driven transcriptomic-dataset.

The MILAN-method mapped CD4^+^/CD8^+^T single-cells, within tumour-tissue from SKCM and GBM-patients **(Supplementary-Table.4),** embedded inside various anatomical-niches, including (i) tumoural, peri-tumoural and non-tumoural areas; (II) non-vascular, peri-vascular and vascular-zones within the tumoural-area; and (III) (GBM-specific) haemorrhagic-zones with vascular-leakage **(Fig.5a-h; FigS9a-d).** Expectedly, SKCM exhibited higher CD8^+^T cell-density, than GBM **(Fig.5i).** Within SKCM (but not GBM), there was an enrichment of CD8^+^/CD4^+^T cells in peri- and non-tumoral areas, while SKCM-tumoral areas mainly enriched CD8^+^T cells **(Fig.5j, Fig.S10a-c).** Within tumoural areas, SKCM and GBM CD4^+^/CD8^+^T cells particularly preferred peri-vascular zones **(Fig.5j; Fig.S10a).** GBM-CD4^+^/CD8^+^T cells also enriched within dysregulated hemorrhagic-zones **(Fig.S10d-e).** Interestingly, CD4^+^T cells exhibited relatively better spatial-proximity to CD8^+^T cells in SKCM, rather than GBM, in almost all anatomical-locations **(Fig.S10f).**

**Figure 5.**
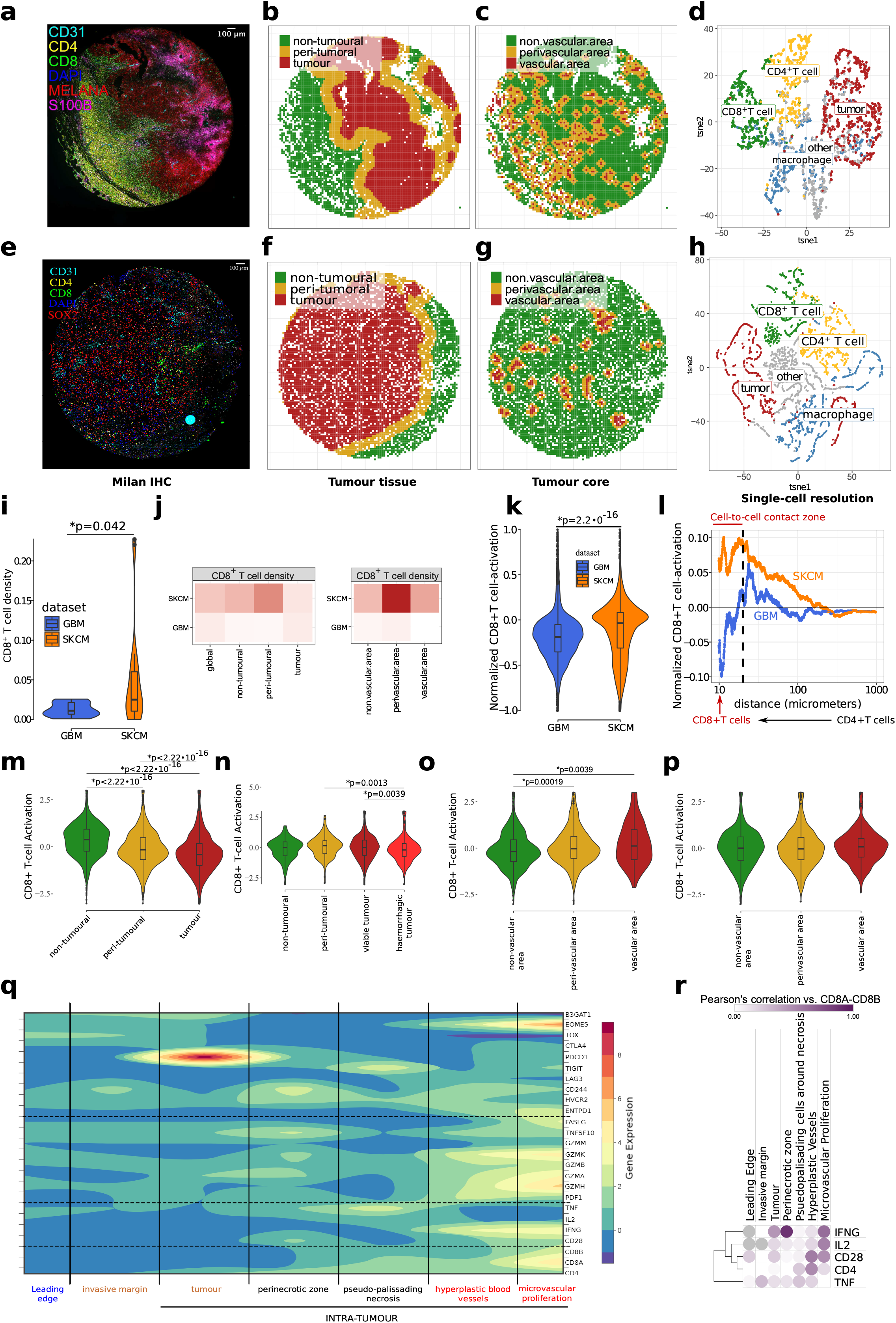
Spatial immuno-mapping of melanoma or glioblastoma tumour-tissue. **(a,e)** Multiple Iterative-Labelling by Antibody-Neodeposition (MILAN)-method derived representative immunofluorescent overlay-images of indicated immune/cancer-cell markers in SKCM **(a),** or GBM **(e)** tumour tissue sections. Digital representations of indicated tumour-tissue anatomical areas **(b,f)** or vasculature areas **(c,g)** in SKCM **(b,c)** or GBM **(f,g),** based on annotation marker expression **(a/e). (d,h)** tSNE-map of a random subset of main annotated cell types across analysed tumour tissue samples in SKCM (n=10-samples across 10-patients) **(d)** and GBM (n = 11-samples across 8-patients) **(h),** using the same colour representations as previously **(a,e). (i,j)** Density analysis of CD8^+^T cells in GBM/SKCM: Violin plot indicates CD8^+^T cells density cross the entire tissue section (i), while heatmaps show relative distribution across indicated areas within each tumour type **(j). (k-p)** Activation scores were calculated for each CD8^+^T cell based on CD69/OX40 vs. LAG3/TIM3 expression levels (Fig.S11a-f). Violin plot **(k)** depicts overall activation of all CD8^+^T cells across all sections for SKCM/GBM. Line plot **(l)** shows median activation of each CD8^+^T cell, relative to its distance from nearest CD4^+^T cell in SKCM/GBM (for statistics, see Fig.S11j). Finally, activation scores of CD8^+^T cells were defined across various tumour anatomical-regions defined in SKCM **(m,o)** or GBM **(n,p). (q)** Contourplot for z-score expression profile of indicated genes across the 7 (indicated) GBM-tumour micro-dissected anatomical sections from IVY GBM Atlas-Project (IVY-GAP) GBM patient dataset (n=270 anatomical sections from 36 GBM-patients). **(r)** Pearson’s correlation of indicated genes with CD8^+^T-signature (*CD8A/CD8B*) within the IVY-GAP GBM-patient dataset. In **i, k, m-p,** Mann-Whitney test with Holm-method for multiple-testing comparison was applied (FDR-adjusted p-values are indicated). Statistical significance threshold was *p<0.05. In **j, o-p,** vasculature analyses were restricted to only regions within tumour “core” (non/peri-tumoural zones were excluded).

Next, we investigated CD8^+^T cell-activation relative to their spatial-distribution. Since tissue-level cytokine-labelling is unfeasible, we opted for a previously established CD69-OX40 (CD8^+^T cell-activation markers^2,10,13,14,31^) vs. TIM3-LAG3 (IRs^2,10^) model of phenotypic-activation/exhaustion **(Fig.S11a-f)**.^19^ Again, SKCM-CD8^+^T cells showed higher phenotypic-activation (CD69-OX40>TIM3-LAG3) than GBM-CD8^+^T cells **(Fig.5k),** irrespective of anatomical-locations **(Fig.S11g-h).** Intriguingly, spatial-proximity of CD4^+^::CD8^+^T cells (with <20μm relative-distance representing cell-to-cell contact-range^55^) significantly amplified phenotypic-activation of SKCM-CD8^+^T cells, but not GBM-CD8^+^T cells **(Fig.5l; Fig.S11i-j).** Herein, the farthest intra-tumoural CD8^+^T cells exhibited phenotypic-exhaustion (TIM3-LAG3 ≥CD69-OX40) **(Fig.5l; Fig.S11i-j).** Spatially, SKCM-CD8^+^T cells were more phenotypically-activated in non- and peri-tumoral areas, but phenotypically-exhausted within the tumour **(Fig.5m):** a well-documented phenomenon.^14^ However, for GBM-CD8^+^T cells, such trends weren’t readily visible **(Fig.5n).** Interestingly, significantly in SKCM and marginally in GBM, vascular-zones had relatively higher CD8^+^T cell phenotypic-activation **(Fig.5o-p).** Importantly in SKCM/GBM scRNAseq-data, *CD69*^HIGH^*HAVCR2*^LOW^ and *HAVCR2*^*HIGH*/+^*CD69*^LOW^ CD4^+^/CD8^+^T cells, primarily exhibited IL2-enriched and IL2-depleted profiles (especially in GBM), respectively **(Fig.S12a).** Altogether, this suggested that despite GBM’s anti-lymphocytic niche, vascularised-zones might provide threshold support to (incoming) T cells.

To confirm this, we interrogated IVY-GAP **(Fig.S12b).** Consistent with above observations, GBM tumour-core (tumour÷necrotic zones), invasive-margin and leading-edge showed “depletion” of CD4^+^/CD8^+^T cell-markers **(Fig.5q).** Herein, CD8^+^T_SDF_-profile was clearly visible, as effector-function genes failed to uniformly positively-correlate with *CD8A/B* **(Fig.5r).** Interestingly, an appreciable proportion of GBM-CD4^+^/CD8^+^T cell-markers associated with vascular-zones **(Fig.5q).** Moreover, these vascular GBM-CD8^+^T cells co-associated with slightly CD8^+^T_EXT_-like phenotype and *CD4* **(Fig.5r).** Accordingly, these vascular-zones had relatively more *IFNG/IL2* **(Fig.S12c-d):** a sign of CD8^+^T_EXT_-supportive niche. Notably, while encouraging, these vascular *CD8A/CD8B, IFNG/IL2, CD4* expression-levels were marginally above zero, thereby signifying an unstable landscape **(Fig.S12e-f).** Thus, SKCM-tumours (but not GBM) have better CD4^+^/CD8^+^T cell-accessibility and higher probability of CD4^+^::CD8^+^T cell-to-cell contact supporting CD8^+^T cell-activation. Nevertheless, the vascularised-zones within GBM exhibited marginal CD8^+^T_EXT_-supportive niche.

### Anti-PD1 immunotherapy facilitates, rather than suppress, CD8^+^T_SDF_ landscape

We wanted to provide a “cause-effect” validation of our descriptive-insights on CD8^+^T_EXT_/T_SDF_. Hence, we pursued integrated (retrospective) analyses of published immuno-oncology clinical-trials profiling tumours, cross-sectionally (i.e., profiled before anti-PD(L)-1/CTLA4 ICBs, or combination thereof)^6,7,57–59^ or longitudinally (i.e., profiled before and after, anti-PD1 ICB).^60,61^

To validate organ-specific niches’ role in aiding/impeding anticancer immunity, we performed correlation-analyses of tumoral TMB-score (antigenicity-surrogate) assessed before/pre ICB-treatment, vs. patient-OS observed after/post ICB-treatment **(Fig.6a).** Importantly, while SKCM primary-tumours/metastasis in skin, lymph-node or non-lymphoid (extra-cranial) organs had a continuum of positive-correlations, yet GBM, UVM and SKCM brain-metastasis had negative-correlations **(Fig.6a)**.^7^ Accordingly, pre-ICB levels of various immuno-oncology biomarkers (*CD274, CD8A/B, IFNG*) and our csCD8^+^T cell-signature, predicted prolonged ICBs-driven patient-OS with significantly higher efficiency in SKCM^57,59^ or BLCA,^58^ compared to GBM^6^ **(Fig.S13a).** Of note, even in SKCM/BLCA, TCR/BCR-clonality were not as reliable predictive-biomarkers as TMB-score **(Fig.S13a).** Overall, this verifies organ-specific immuno-tolerance, especially in brain, co-opted by malignancies to suppress immunogenicity.

**Figure 6.**
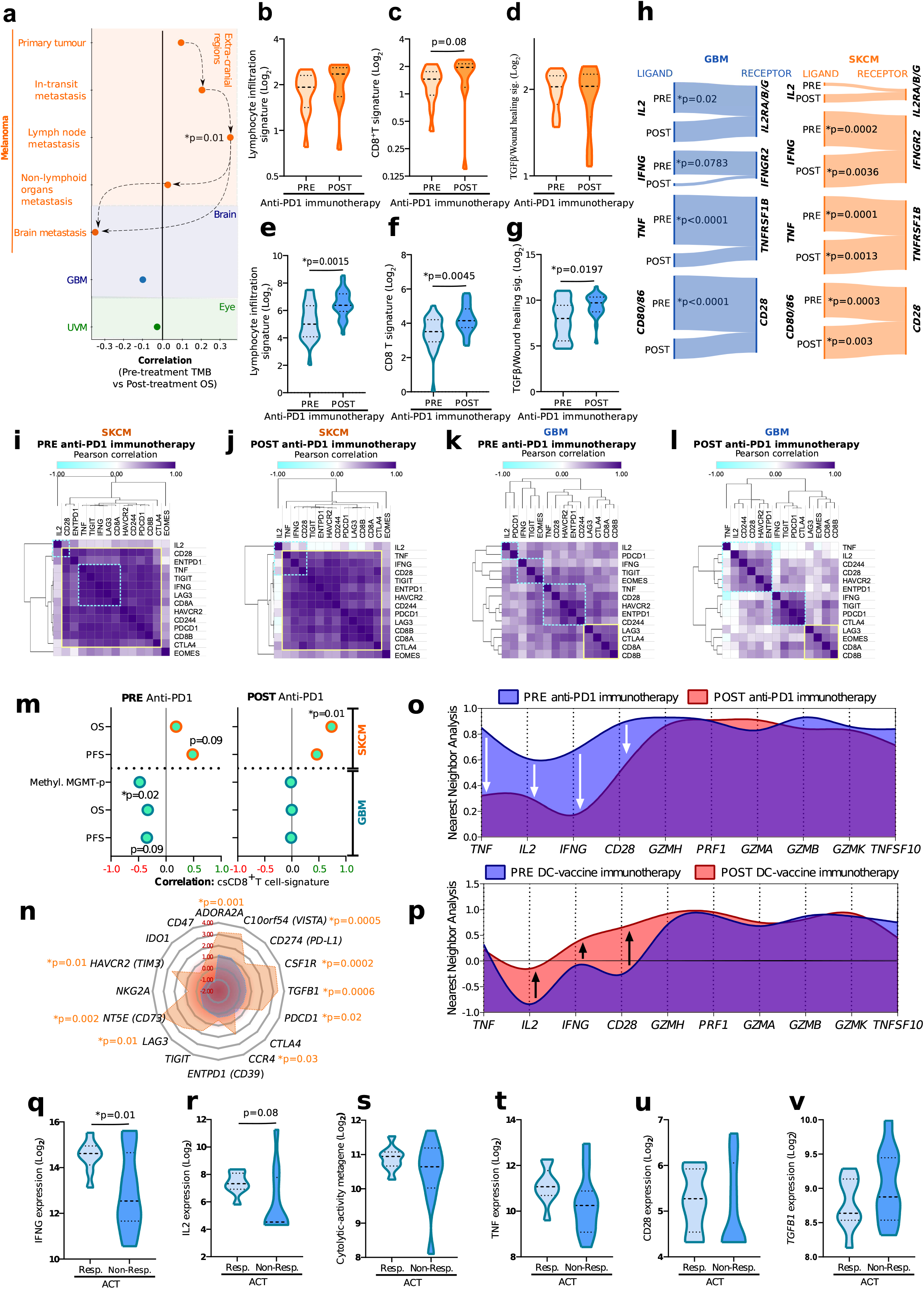
Immunogenomics in immuno-oncology clinical-trials. **(a)** Pearson’s-correlation between tumour mutational burden (TMB)-score in SKCM primary-tumours (n=20), SKCM-metastasis (in-transit, n = 18; lymph-nodes/LNs, n=50; non-lymphoid/extra-cranial organs, n=71; brain, n=15), GBM (n=82) or metastatic-UVM (n=20), before immune-checkpoint blocking (ICBs)-immunotherapy (anti-PD(L)l, anti-CTLA4 or anti-PD(L)l+anti-CTLA4), vs. patient overall-survival (OS) after ICB-immunotherapy. Clinical trials involving pre/post-ICB (anti-PD1) treatment derived tumour-tissue from SKCM (Pre, n=13; Post, n=11) or GBM (Pre, n=24; Post, n=25) patients were used for creating: **(b-g)** violin-plots for indicated signaturelevels (Mann-Whitney test); **(h)** Sankey-plots indicating Spearman-correlations between indicated ligand-receptor genes; **(i-l)** Correlation-matrices for csCD8^+^T cell-signature; **(m)** Pearson’s-correlation between csCD8^+^T cell-signature vs. patient-level positive clinico-pathological prognosticators (OS; progression-free survival, PFS; O6-methylguanine-DNA methyltransferase, MGMT promoter-methylation), **(n)** Radar-plot of TIDE-prediction z-scores for clinically-relevant immune-checkpoint/IRs coding-genes in TCGA-SKCM (n=413) or TCGA-GBM (n=158) (higher z-score and *p<0.05 means patient-subgroup with low target-gene expression shows significantly prolonged-overall-survival/OS for high-expression of CD8^+^T cell-signature at median-expression cut-off). Clinical-trial involving pre/post-dendritic cell (DC)-vaccination derived tumour-tissue from GBM (Pre, n=6; Post, n=6) patients was used for creating cubic-spline representation of nearest-neighbour (Pearson’s-correlation) analyses for indicated genes within the GBM anti-PD1 immunotherapy **(o)** or DC-vaccination clinical-trials **(p)** (1000-permetuations). Clinical-trial involving adoptive T-cell therapy (ACT) administered to GBM-patients, with gene-expression profiling of ACT-products, was used for creating, violin-plots for indicated genes/signature-levels **(q-v),** in ACT-responding (Resp., n=9) or nonresponding (Non-Resp., n = 11) GBM-patients (unpaired t-test with Welch’s correction). Herein, Resp. have >8-months’ time-to-progression (TTP) or no-progression (post-ACT), whereas Non-Resp. have <8-months TTP. Statistical-significance threshold was *p<0.05. For different genesignatures, see Supplementary-Table.2.

Interestingly, within longitudinal-samples around (neo)adjuvant anti-PD1-immunotherapy,^60,61^ while both SKCM and GBM experienced an increase in signatures of lymphocytes-infiltration and *CD8A/CD8B,* after anti-PD1-immunotherapy, yet only GBM (but not SKCM) significantly enriched for TGFβ/wound healing **(Fig.6b-g).** Qualitatively, while in SKCM, *IL2, IFNG, TNF* and *CD80/86* improved/maintained correlations with their cognate receptor-coding genes, after anti-PD1-immunotherapy; yet in GBM, these correlations were consistently disrupted **(Fig.6h).** Quantitatively, anti-PD1-immunotherapy moderately increased *CD28-IFNG* expression in SKCM, while in GBM it mainly improved *TNF* **(Fig.S13b-c).** Accordingly, csCD8^+^T cell-signature’s genes exhibited high correlations/concordance in SKCM, both before/after anti-PD1-immunotherapy **(Fig.6i-j);** whereas in GBM, anti-PD1-immunotherapy reduced, rather than improved, pre-existing low correlations/discordance **(Fig.6k-l).** Operationally, while csCD8^+^T cell-signature consistently positively-correlated with progression-free survival (PFS) and OS in SKCM-patients, especially after anti-PD1-immunotherapy **(Fig.6m)**;^61^ yet in GBM-patients, csCD8^+^T cell-signature exhibited negative (before) or null (after) correlations with PFS/OS or MGMT promoter-methylation **(Fig.6m)**.^60^ These patterns were consistent with TCGA-immunogenomics and cross-sectional trials’ observations, thus compensating for statistical constrains of these trial’s small cohort-sizes. In-conclusion, CD8^+^T_EXT_ are primed for ICB-responsiveness, while ICBs rather than reinvigorating CD8^+^T_SDF_-landscape, instead facilitate lymphocytic effector-dysfunction and TGFβ/wound healing.

### IL2/IFNγ-integrating cellular immunotherapy can amend pro-CD8^+^T_SDF_ landscape

Our observations indicated general non-responsiveness of CD8^+^T_SDF_ to ICBs (even beyond PD1-axis). To confirm this, we applied Tumour Immune Dysfunction and Exclusion (TIDE)-model, which predicts therapeutic-feasibility of ICB-targets by mathematically estimating if lowering tumoural IRs gene-expression will result in CD8^+^T cell-markers predicting prolonged patient-OS (in TCGA-dataset).^11^ Indeed, most ICB-targets **(Fig.6n)** were predicted to sustain significant therapeutic-feasibility in SKCM, but not in GBM. Interestingly in GBM, the top-5 predicted ICB-targets beyond *PDCD1* were “positioned” within TGFβ/wound healing or macrophage-pathways, thus verifying our immunogenomic/single-cell mapping. Still, this forecasted a relatively low-feasibility for ICBs in reinvigorating GBM-CD8^+^T_SDF_.

Our results indicated that low antigenicity and pro-CD8^+^T_SDF_ landscape impeded anti-GBM efficacy of ICBs. A possible solution to these hurdles is cellular immunotherapies, that can augment TCR-diversity and antigen-specific anticancer immunity. Indeed sporadic success has been reported for cellular immunotherapies like dendritic cell (DC)-vaccines or adoptive T cell-therapies (ACT) against GBM.^62,63^ Henceforth, we wondered which CD8^+^T_SDF_ landscape-disparities were amended (or not), by these. In DC-vaccines trial with longitudinal-sampling,^64^ GBM-patients experienced palpable (but insignificant) increase in signatures of lymphocytes-infiltration and *CD8A/CD8B* **(Fig.S13d-e),** but without increase in TGFβ/wound healing **(Fig.S13f);** a relatively good sign. Interestingly, compared to pre vs. post anti-PD1 immunotherapy in GBM, the pre vs. post DC-vaccination correlation for effector-function (*IFNG-IL2-TNF-CD28*) genes was significantly higher **(Fig.S13g).** Consequently, DC-vaccines increased co-association of *IFNG-CD28* (but not IL2) with cytolytic-activity genes, better than anti-PD1-immunotherapy, which reduced these co-associations **(Fig.6o-p),** thus highlighting IL2 as a DC-vaccine limiting-factor.

Although promising immunological/clinical-responses were reported for DC-vaccine trials,^64^ from which above samples originated, yet matched patient prognostic data weren’t verifiably available. We therefore pursued therapeutic dissection of anti-GBM adoptive T cell-therapy (ACT). We examined the expression-profiles of (PBMCs-derived) autologous cytomegalovirus (CMV)-specific^65^ ACT-products, applied in a recent clinical-trial reporting positive therapeutic-efficacy in GBM-patients.^63^ We compared expression of various genes across ACT-products associated with pathologically responding (≥8-months time-to-progression, or no-progression) and non-responding GBM-patients (<8-months time-to-progression). The threshold of 8-months was based on the 75th percentile cut-off for the entire dataset. Remarkably, ACT-products of responding-patients had higher *IFNG, IL2* **(Fig.6q-r)** and elevated *TNF,* cytolytic-function genes or *CD28* **(Fig.6s-u),** compared to non-responders. Conversely, non-responders’ ACT-products had elevated *TGFB1-levels* **(Fig.6v).** Altogether, this indicated that cellular immunotherapies have higher probability than ICBs in (partly) overcoming pro-CD8^+^T_SDF_ landscape, with IFNγ-IL2 and TGFβ/wound healing being critical efficacy-determinants.

## DISCUSSION

IRs-centric interpretation of CD8^+^T cell-exhaustion gives an ambiguous impression of pancancer CD8^+^T_EXT_ enrichment, despite contrasting ICB-responsiveness across tumours.^2,9,12,14^ Our deep-dissection of clinical CD8^+^T cell-states revealed tumour-specific landscapes and organ-specific niches as important players in shaping CD8^+^T_EXT_/CD8^+^T_SDF_; with these pre-existing CD8^+^T cell-states determining immunotherapy-responsiveness.^2^ We observed that critical disparities on levels of lymphocytic IFNγ/IL2 and TGFβ/wound healing in immunotolerant organs-associated malignant lesions (e.g., brain-GBM, SKCM brain-metastasis, eye-UVM, ovary-OV)^26^ can facilitate immunotherapy-resistant CD8^+^T cell-state. Contrastingly, CD8^+^T_EXT_-enriching tumours (e.g., SKCM/LUAD/BLCA) enjoy threshold immunological support from specific organ-niches, allowing endurance of effector-functionalities. Since T cell exhaustion/dysfunction evolved to avoid autoimmunity,^10,66^ the increase in TGFβ/wound healing following anti-PD1-immunotherapy in GBM could be an attempt at avoiding brain-associated autoimmunity.^10,27^ These conclusions align with: symptomatic SKCM brain-metastasis showing ICB-hyporesponsiveness;^28^ TGFβ-signalling, low-antigenicity, and strong lymphocyte auto-regulation maintaining immune-privilege;^27^ TGFβ’s pro-exhaustion/quiescence activity via IL2-downregulation;^10,43^ ICB non-responsiveness of OV/UVM;^5,8^ and exhausted-CD8^+^T cells adopting organ-specific effector-profiles during chronic viral-infection.^23^ Lastly, our observations of organ-specific imprinting on tumoural lymphocytic-features can also be ascribed to other confounding factors e.g., bacteriome/virome.^67^

We interpreted CD8^+^T_EXT_/CD8^+^T_SDF_ as population-level phenomenon, rather than specific CD8^+^T cell-subpopulations/lineages, to allow cross-omics translation of CD8^+^T cell-states.^2,3,12,14,38,45^ Canonical CD8^+^T_EXT_-trajectory proceeds from pre/early-exhaustion (ICB-responsive) to late-exhaustion (ICB-hyporesponsive) under chronic antigen-exposure.^10,12^ While this applied to CD8^+^T_EXT_ in immunogenic/antigenic SKCM, yet in low-immunogenic/antigenic GBM, CD8^+^T_SDF_ didn’t imitate late-CD8^+^T_EXT_, possibly because former develop from suboptimally antigen-primed anergic/senescent CD8^+^T cells. This is substantiated by low TCR-diversity in GBM, coupled with effector/cytolytic (transcriptomic/epigenomic) dysfunctions, maladaptive pro-death/senescent stress and cell-cycle defects in CD8^+^T_SDF_, without signs of chronic antigen-exposure (multiple-IRs or memory-differentiation). Lastly, ICB hyporesponsiveness in various SKCM-patients can be attributed to defective late-CD8^+^T_EXT_.^2,12,14^ In our settings, this is supported by presence of low-functional SKCM-CD8^+^/CD4^+^T single-cells, TGFβ/wound healing in SKCM patient-subsets, and decrease in TMB-score vs. patient-OS correlation, after ICB-immunotherapy, in non-lymphoid organ-associated SKCM-metastasis.

Beyond non-spatial, we also revealed spatial CD8^+^T cell-characteristics – a novel feature of our study. Pro-CD8^+^T_EXT_ tumour (SKCM) had more T cells-enriched normal-adjacent or peri-tumoral tissue, than pro-CD8^+^T_SDF_ (GBM); further highlighting the role of organ-specific niches in better lymphocyte-accessibility. This aligns with high CD8^+^T cells at tumour invasive-margins predicting better ICB-responses.^14^ Curiously, tumour-vasculature supported CD8^+^T_EXT_-like features across both SKCM/GBM, while non-vascularised zones enforced tumour-specific CD8^+^T_EXT_/T_SDF_ states. Remarkably, spatial-proximity of CD4^+^::CD8^+^T cells (especially in cell-to-cell contact-range) facilitated better CD8^+^T cell-activation in SKCM, but not GBM, thereby confirming dysfunctional lymphocytic-interactions in GBM.

Our observations positioned CD8^+^T_SDF_ beyond the therapeutic “window-of-opportunity” for (neo)adjuvant ICB-immunotherapy. Instead we documented that cellular immunotherapies (DC-vaccines or ACT),^62,68^ may partially “overcome” IFNγ or IFNγ/IL2-disparities in GBM, respectively. Nevertheless, either of these approaches alone may fail to ensure recurrence-free GBM-patient survival.^68^ Thus, a multi-modal immunotherapeutic-regimen integrating nextgeneration versions of these cellular immunotherapies with proficient IL2-production,^4^ might be needed to overcome GBM. Concurrent combination with anti-CSF1R or anti-TGFβ ICBs is also worth testing.^69^

Our study also delivers some valuable resources/methods for differentiating CD8^+^T_EXT_ from CD8^+^T_SDF_ in clinical omics-datasets e.g., csCD8^+^T cell-signature, “first-in-class” ML prediction model, novel single-cell immune-trajectory annotation approach (FIAT), mathematical CD8^+^T single cell-cycle modelling, and CD4^+^::CD8^+^T-cells’ spatial-proximity based activation modelling. However, while a specific caveat of our study-design is the lack of patient matching across all omics-platforms; yet, our approach does provide cross-cohort/cross-platform validation. Nevertheless, it would be desirable to reproduce our observations via multi-omics spatial biomarker-screening in a prospective clinical-trial.

Altogether, we believe our study provides key resources and a novel reference-map for clinical CD8^+^T cell-exhaustion/dysfunction in cancer, and a critical overview of immunotherapy-deterministic factors. Overall, this can guide the design of innovative immunotherapy-regimen for hard-to-treat cancers.

## Supporting information

Supplemental box 1

## Acknowledgements

We acknowledge Dr. Alexander Huang and Dr. Tara Mitchell (UPenn, USA) for helping us in accessing the patient survival/prognostic data from their SKCM longitudinal clinical trial with (neo)adjuvant anti-PD1-immunotherapy. We acknowledge Dr. Corey Smith, Dr. Sriganesh Srihari, and Dr. Rajiv Khanna (QIMR Berghofer Medical Research Institute, Australia) for helping us in accessing the gene-expression data and patient pathological response data from their GBM clinical trial with ACT-immunotherapy. We thank Dr. Nicholas Van Baren & Dr. Pierre Coulie (UCL, Belgium), Dr. Patrizia Agostinis & Yichao Hua (KU Leuven-VIB, Belgium) for attentively reading the manuscript and providing valuable suggestions and feedback. This study is supported by Research Foundation Flanders (FWO) (Fundamental Research Grant, G0B4620N to ADG; Excellence of Science/EOS grant, 30837538, for ‘DECODE’ consortium, for ADG, BJVDE), KU Leuven (C1 grant, C14/19/098 and POR award funds, POR/16/040 to ADG), and Kom op Tegen Kanker (KOTK/2018/11509/1 to SDV, ADG, FDS; and KOTK/2019/11955/1 to ADG). IV is supported by FWO-SB PhD Fellowship (1S06821N). DMB is supported by the Belgian Federation against Cancer grant nos. 2018-127 and 2016-133 and by a grant from Fondation Roi-Baudouin to ST. ST is further supported by a Senior Clinical Investigator award of FWO. This work is also supported by KU Leuven grant C14/17/084 to FDS and C3/19/053 to FDS and FMB, FWO grant G0I1118N to FDS, the Leuven Kankerinstituut (LKI), and an FWO fellowship to JM (1156520N). AMA is supported by the Opening The Future Foundation and LKI. The authors also thank the multiplexed immunohistochemistry platform of KU Leuven.

## METHODS

### 1. Study Design

The design and workflow of this study is described in **Fig.1a.** We started with a comprehensive systematic literature meta-analyses, involving PubMed-derived research publications, to construct a consensus csCD8^+^T cell-signature, in order to qualitatively drive our immunogenomics, single-cell as well as spatial tumour-tissue analyses. Immunogenomics analyses was based on large-scale publicly-available human cancer patients-derived transcriptomic (tumour bulk-RNAseq or bulk-microarray), proteomic (tumour bulk reverse-phase protein array/RPPA) or epigenomics (tumour bulk DNA-methylation or bulk Assay for transposase-accessible chromatin using sequencing ATACseq) datasets derived from The Cancer Genome Atlas (TCGA), The Cancer Proteome Atlas (TCPA), or Gene Expression Omnibus (GEO) databases. This was complemented by similar analyses of large-scale publicly-available human (healthy) subjects-derived normal-organ/tissue transcriptomics datasets from Genotype-Tissue Expression (GTEx) database. This immunogenomic initiative involved genetic-signatures analyses, immune-deconvolution, network analyses, mathematical modelling and/or GSEA, across >3300 patients spanning 7 different cancer-types or >1300 subjects spanning 6 different organ/tissue-types. This was followed by hypothesis-driven validation in single-cell (sc)RNAseq datasets across 47 patients spanning SKCM/GBM, spatial multiplex-immunohistochemistry analyses of human tumour tissue via MILAN method across 18 patients spanning SKCM/GBM and spatial bulk-RNAseq analyses of GBM tumour anatomical-sites isolated via laser-assisted dissection across 36 GBM-patients. Finally, we validated our descriptive analyses derived conclusions in immuno-oncology clinical trials: cross-sectional (i.e., tumours profiled pre-treatment) clinical trials applying ICBs across 674 patients spanning 4 cancer-types and longitudinal (i.e., tumours profiled both pre/post-treatment) clinical trials applying cancer immunotherapy (anti-PD1 ICB, DC-vaccines or ACT) across 64 patients spanning SKCM/GBM. All analyses were integrated via consistent qualitative analyses of csCD8^+^T cell signature-associated genes further complemented by state-of-the-art computational immunology and mathematical modelling.

### 2. Systematic literature meta-analyses to construct csCD8^+^T cell-signature

PubMed was searched on 14/09/2019 for applicable studies conducted in human cancer patients without restriction on timeline (studies were automatically retrieved between 1986 to 2019). The following search string was used (with “Human” species filter): (“cd8 T” OR “cd8+T” OR “cd8+” OR cd8) AND (exhaustion OR exhausted OR anergy OR anergic) AND (cancer OR tumour OR tumor OR neoplasm). To distinguish potentially relevant studies, the catalogue of articles identified in the earlier search, were also scanned manually. Studies within the catalogue were considered qualified if they met all of the subsequent criteria: involved experimental/direct analyses of human cancer patient-derived lymphocytes, *in vitro/ex vivo* human patient-derived T cell analyses, and bulk-RNAseq, microarray, single-cell (sc)-RNASeq or other genetic/omics analyses of tumour-isolated lymphocytes. Studies were excluded because of following reasons: (1) non-experimental genetic signature analyses of lymphocytes, (2) analyses with animal/rodent models (syngeneic, allogeneic or xenografts), (3) studies with virus-induced exhaustion, (4) human lymphocytes analyses not focussing on exhaustion/anergy, (5) inconclusive T cell analyses, (6) tumour bulk-RNASeq/microarray analyses or (7) letters, reviews, commentary, perspectives, case reports, conference abstracts, editorials or expert opinion. Overall, these search criteria helped short-list 96 research studies annotated in **Supplementary Table 1.** Herein, only those biomarkers were considered for annotation of T cell exhaustion phenotype (i.e., dysfunction, senescence, differential exhaustion, or early-exhaustion/polyclonality) for which a consistent and distinct expression profile (i.e., either High or Low, but not both or simply expression positivity) was reported across ≥ 3 research studies. For the chronically-sensitised (cs)CD8^+^T cell signature **(Supplementary Table 2),** only biomarkers associated with dysfunction/senescence or differential T cell exhaustion were considered. Early-exhaustion/polyclonality biomarkers were not considered for this signature since they could have skewed the analyses in favour of early/mid-effector or pre-exhausted T cell-phenotype, which is beyond the focus of this study.

### 3. Immune-MSigDB datasets’ network for csCD8^+^T cell signature

The ShinyGO v0.61 workflow^70^ (http://bioinformatics.sdstate.edu/go/) was utilised to enrich experimental-dataset hits from the Immune-MSigDB compendium,^71^ using genes within the csCD8^+^T cell signature. The enrichment exercise was restricted to Human datasets, and top-20 most significant terms with a false-discovery rate (FDR) p-value cut-off of 0.05. For the network an edge cut-off of 0.3 was applied. The resulting network depicted relationships between different datasets (nodes), with connection between two datasets depicting sharing of ≥20% genes (thickness of edges was proportional to the extent of overlapping genes). Moreover, nodes with darker color depicted significant enrichment of gene sets whereas the size of nodes was proportional to the number of genes.

### 4. Immuno-transcriptomic analyses for human peripheral CD4^+^/CD8^+^ T cells

RNA-sequencing (RNAseq)-based gene-expression data for human peripheral unstimulated or naive, and stimulated or activated, CD4^+^T cells or CD8^+^T cells was derived from an existing dataset (available as “Celltypescore” module within DICE tools workflow: https://tools.dice-database.org/celltypescore/) as a transcripts-per-million (TPM)-score and further log_2_ transformed.^20^ Briefly, the original investigators^20^ had FACS-sorted these human CD4^+^/CD8^+^T cells from peripheral blood mononuclear cells (PBMC) and either left them untreated (hence naive) or activated them ex *vivo* with Dynabeads Human T-Activator CD3/CD28 (ThermoFisher Scientific), wherein the bead-to-cell ratio was 1:1. This activation protocol was carried out for 4 h at 37°C following by preparation of samples for further RNAseq analyses. For further details relevant for this work-flow please refer to the previous publication.^20^ For this study, we analysed the expression of csCD8^+^T cell signature genes **(Supplementary Table 2)** in this dataset.

### 5. Meta-analyses of anti-PD(L)l immunotherapy efficacy in cancer patient clinical trials

Forest plot-derived clinical (pooled) objective response rates (ORRs) from largest clinical trials (at least >10 patients; 2012-19) applying anti-PD1 or anti-PDL1 mono-immunotherapy (without combining with other anticancer therapies and without any biomarkers-based patient pre-selection) in melanoma (NCT01866319, NCT01844505), non-small cell lung cancer (NSCLC; NCT01295827, NCT01642004, NCT01772004, NCT02008227, NCT01673867), bladder cancer (NCT02256436, NCT02387996, NCT01693562, NCT01772004, NCT02108652), breast carcinoma (NCT01772004, NCT01375842), ovarian cancer (NCT02674061, UMIN000005714, NCT00729664, NCT01772004) or glioblastoma (NCT02017717, NCT02336165) were derived from a recently published PRISMA-guidelines adherent literature meta-regression/meta-analyses (accessed from supplementary document: 236800_3_supp_6337488_qbcxtt.docx).^8^ Corresponding median tumour mutational burden (TMB) scores were also derived from the same source.

### 6. Immuno-transcriptomic analysis with The Cancer Genome Atlas (TCGA) datasets

We accessed patient overall-survival (OS) durations and tumour gene expression profiles (bulk-tumour RNAseq-based) for skin cutaneous melanoma (SKCM, n=470 patients), lung adenocarcinoma (LUAD, n = 574), bladder cancer (BLCA, n=426), breast cancer (BRCA, n = 1212), ovarian cancer (OV, n=427), uveal melanoma (UVM, n=79) or glioblastoma (GBM, n = 171) patients, from Xena^72^ (http://xena.ucsc.edu), which analyzes TCGA and GTEx with a similar workflow and thus reduces technical variation. Survival analyses were performed using the Lifelines Python package (https://zenodo.org/record/4136578#.X8fe52hKiM8). Additional phenotypic data was obtained from the FireBrowse portal (a Broad Institute GDAC Firehose analyses pipeline: http://firebrowse.org/),^73^ or TCGA PanCancerAtlas Immune Response Working Group’s Cancer Research Institute (CRI) iAtlas Explorer.^25^ Normalizations/data corrections were done where needed and specified in the analyses as applicable. Raw data associated with CRI iAtlas Explorer was accessed from the dedicated GDC portal (https://gdc.cancer.gov/about-data/publications/panimmune) or accessed from supplementary documents accompanying published research on CRI iAtlas (mmc2.xlsx or PanImmune_GeneSet_Definitions.xlsx).^25^ Publication sources for different TCGA datasets are accessible on this webpage: https://www.cancer.gov/about-nci/organization/ccg/research/structural-genomics/tcga/publications. All the different immunogenetic signalling or pathway signatures used for interrogation of TCGA datasets (and their respective sources) are elaborated in **Supplementary Table 2.** Of note, the number of patients for which expression or clinical data was drawn from above sources was largely consistent. Minor variations in patient number occurred, due to differential qualitative thresholds or, analyses-specific qualitative cut-off constraints. These variations created no biologically/clinically-relevant limitations or inconsistencies within our study. Alternative TCGA-based workflows (e.g., The Cancer Immunome Atlas or TCIA,^33^ and Tumor Immune Estimation Resource or TIMER^74^) were utilised to cross-confirm the consistency of observations derived from above workflows (data not shown), to ensure cross-platform uniformity. Almost all observations derived from TCGA datasets were verified in independent tumour scRNAseq datasets, spatial-transcriptomics profiling dataset as well as bulk-tumour gene expression profiles from immuno-oncology clinical trials.

#### a. General gene-expression analyses

Expression analyses (IFNG, TNF, IL2, CD28) and correlation analyses (correlation to CIBERSORT CD8^+^ T cell fractions, see below) were performed based on the Toil-recompute RSEM TPM data present in Xena. Values were log_2_(TPM + 1) transformed from the original log_2_ transformation with different pseudo-count, to be consistent with other analyses like SKCM vs. GBM signature-expression or single geneexpression comparative-analyses (for csCD8^+^T cell-signature, CIBERSORT-based CD8^+^T cell-signature, IL2, IFNγ, TNFJ. Correlation matrices were constructed using the similarity function in Pandas. Sankey-plots depicting the correlations between ligand-receptor coding genes were constructed using the SankeyMATIC portal (http://sankeymatic.com/). P-values were calculated with a Welch’s t-test (α=0.05).

#### b. Immune-deconvolution and immuno-genetic signatures/scores

The relative proportions of various immune cell-types estimated via Cell-type identification by estimating relative subsets of RNA transcripts (CIBERSORT-LM22)^75^ method (and multiplied by leukocyte-fraction estimates to derive overall fractions per tissue; e.g., CD8^+^T cells, M0/M1/M2 macrophages), aggregated combinations of these immune-fractions (e.g., lymphocytes, macrophages or CD4^+^T cells), or specific immune-cell signature scores (e.g., lymphocyte infiltration signature-score, TGFβ response-score, IFNγ-response score, or wound healing score) were accessed through the CRI iAtlas workflow (herein, GBM = 153). In order to create the CD8^+^T cell fractions’ quadrant analysis, all quartiles were split of based on the CD8^+^T cell fraction from lowest to highest 25% inferred by CIBERSORT per cancer type individually. Per quartile, the Kaplan-Meier curves were plotted with lifelines. The CoxPh hazard ratios for GBM vs SKCM were estimated with the same package, correcting for age, tumour stage and gender. Features were z-score standardized. Correlation matrices for 25 immune-deconvoluted immune cell fractions were constructed using the similarity matrix function available within the Broad Institute’s Morpheus portal (https://software.broadinstitute.org/morpheus). Lastly, in order to account for interference due to gene over-representation affecting immune-deconvolution for CD4^+^T cell subpopulations, we utilised single sample gene set enrichment analysis (ssGSEA) data delineating enrichment of T_H1_, T_H2_ or T_REG_ per patient across the entire TCGA datasets available from TCIA workflow (herein GBM, n=160).^33^ Within this workflow we used a normalized enrichment score (NES) cut-off >1, and FDR (q-value) <0.05.

#### c. Extracellular network analyses

Extracellular network analyses based on TCGA-SKCM or TCGA-GBM datasets was performed using the FANTOM5-database driven Extracellular Networks module within CRI iAtlas Explorer workflow.^25^ This module uses FANTOM5 database of established ligand-receptor, cell-receptor or cell-ligand pairings to extrapolate networks for specific immune-genes or immune cell-fractions relative to tumour immune landscape pathways.^25^ We applied this module at 45% abundance-threshold and 1.5 concordance threshold while selecting specific genes-of-interest (*IFNG, IL2, TNF* or *CD28*) and cell-of-interest (CD8^+^T cell-fractions), with dagre-layout and connectivity-edges based on tumour immune-landscape pathways. The output was represented as original extracellular network or pie-charts summarizing the number of times connectivity-edges enriched specific tumour immune-landscape pathways.

#### d. Antigen-landscape and T cell receptor (TCR) diversity analyses

Pre-computed TCR diversity metrics and single-nucleotide variations (SNVs)-derived neoantigen levels were accessed from TCGA’s GDC portal (https://gdc.cancer.gov/about-data/publications/panimmune) or, supplementary documents accompanying published research on CRI iAtlas (mmc2.xlsx or PanImmune_GeneSet_Definitions.xlsx).^25^ Briefly, the TCGA investigators delineated neo-antigenic peptide predictions via NetMHCpan v3.0-driven workflow as described previously.^25^ As a quality cut-off, only those patient data points were retained wherein predicted peptides presented with a mutated position, sufficient predicted binding to autologous HLA, and palpable gene expression (threshold: 1.6 TPM), which finally resulted in 103 SKCM-patients and 148 GBM-patients passing the relevant quality thresholds enforced by the TCGA consortium.^25^ Cancer-testis (CT) antigens signature **(Supplementary Table 2)** was calculated with the same workflow as above (herein, SKCM, n=467; GBM, n = 157). Similarly, the TCGA investigators used MiTCR v1.0.3 to procure TCR CDR3 sequences from TCGA RNAseq datasets followed by further computational/mathematical processing as described previously,^25^ to derive TCR Shannon-diversity (SKCM, n=434; GBM, n = 142), TCR Richness (SKCM, n=468; GBM, n=153) or TCR Evenness (SKCM, n=397; GBM, n=126).

#### e. Survival analysis

Where applicable, hazard ratios and confidence intervals (α=0.05) were calculated using Coxphfitter in Lifelines. We compensated for age, gender and tumour stage, using the metadata from Firehose and Xena, using OS as event and OS.time as duration. Kaplan-Meier curves were generated using just OS and OS.time.

### 7. Immuno-transcriptomic comparative analyses

#### a. Tumour-tissue vs. Normal tissue

TCGA-derived tumour tissue (SKCM, LUAD, BLCA, BRCA, OV, GBM) vs. Genotype-Tissue Expression (GTEx: https://www.gtexportal.org/home/)^24^ derived corresponding normal tissue (skin, lung, bladder, breast, ovary or brain) based on the Toil-recompute project in Xena were used to directly compare TCGA and GTEx. UVM could not be compared due to the absence of corresponding normal tissue in the Xena GTEx dataset. Briefly, they re-computed the raw RNAseq data from TCGA or GTEx datasets derived from the UCSC Xena project with a uniform pipeline thereby allowing parallel comparative analyses for respective genes or gene signatures. All values were log_2_(TPM + 1) normalised after correcting for the original transformation and pseudocount in the RSEM TPM data. To compare CD8^+^T cell fractions between normal tissue and the corresponding cancers, both GTEx and TCGA were run through CIBERSORT in absolute mode. Absolute scores were rescaled to values between 0 and 1. In total, we compared above TCGA datasets with following normal organ/tissue counterparts: bladder, breast (mammary tissue), brain (frontal cortex/Ba9 and cortex), lung, ovary, skin (not sun exposed i.e., Suprapubic; and sun exposed i.e., lower leg. This resulted in the total number of samples for brain (Normal Tissue: 207, Primary Tumour: 153), bladder (Primary Tumour: 407, Normal Tissue: 9), breast (Primary Tumour: 1092, Normal Tissue: 179, Metastatic: 7), lung (Primary Tumour: 513, Normal Tissue: 288), ovary (Primary Tumour: 419, Normal Tissue: 88) and skin (Normal Tissue: 557, Metastatic: 366, Primary Tumour: 102) used in subsequent analyses.

#### b. GEO dataset analyses

Gene expression data for (bulk) intra-cranial or brain SKCM metastasis sub-group and (bulk) extra-cranial SKCM-metastasis, were derived from a previously published resource (GSE50496); similarly, gene expression data for (bulk) primary melanoma or benign nevi samples were derived from another published resource (GSE3189). These data were processed through a unified ShinyGEO workflow (https://gdancik.github.io/shinyGEO/).^76^ Herein, extra-cranial SKCM metastasis were sub-grouped as lymphoid (i.e., SKCM-metastasis originating from lymph-nodes or spleen) or nonlymphoid (i.e., SKCM-metastasis originating from adrenal glands, bowel, liver, lung, small intestine or soft tissue). All values were log_2_ transformed and median-values per subgroup were row-normalised for the heatmap representation via Broad Institute’s Morpheus portal (https://software.broadinstitute.org/morpheus).

### 8. Immuno-proteomic analysis with The Cancer Proteome Atlas (TCPA) datasets

We accessed tumour protein-level profiles (bulk-tumour reverse-phase protein arrays (RPPA)-based) for skin cutaneous melanoma (SKCM, n=354 patients), or glioblastoma (GBM, n=205) patients, originating from within the TCGA cohorts-derived TCPA initiative. These proteomic profiles were accessed from the TCPA portal^22^ (more specifically, TCGA Pan-Can 32-L4 dataset downloaded from https://tcpaportal.org/tcpa/index.html) and available proteins were checked for their relative specificity for marking CD4^+^/CD8^+^T cells and/or lymphocytes and accordingly LCK protein was selected for further differential protein-level analyses.

### 9. Epigenetic analyses within TCGA datasets

#### a. Methylation analyses

Methylation data from Illumina Human Methylation 450K arrays were downloaded from TCGA-database for a total of 115 GBM-patients and 475 SKCM-patients (based on matching for corresponding tumour bulk-RNASeq data in TCGA). Methylation of the following genes was investigated: *CD28, FASLG, GZMA, GZMB, GZMH, GZMK, GZMM, IFNG, IL2, PRF1, TNF* and *TNFSF10.* For each gene of interest, CpGs located 500 base-pair (bp) downstream the transcription start site (based on BDTSS definition)^77^ and 1500-bp upstream were selected. The analyses were performed using TCGAbiolinks (version 2.17.1)^78^ and the R package (version 3.6.3).

#### b. ATACseq analyses

TCGA ATACseq data for assorted SKCM or GBM patients was accessed from published resources of TCGA consortium (TCGA-ATAC_DataS9_ImmuneInfiltration_v3.xlsx).^30^ Mathematical approach utilised for estimating ATACseq-based immune-specific analysis, tailored for TCGA data, for peak-to-gene hits related to immune infiltrates was described previously.^30^ Briefly, the TCGA consortium had used published ATACseq data from the hematopoietic cells like CD8^+^T cells to delineate peak-to-gene links specific for only immune cells. They used this to “deconvolute” tumour-associated immune cell features while accounting for background features (i.e., ATACseq accessibility of the hematopoietic cell types like CD8^+^T cells vs. TCGA cohort’s accessibility for each peak). This allowed determination of putative immune-specific features for specific ATACseq peaks.^30^ To further tailor this to lymphocytic activity, correlative analyses was carried out between ATACseq peak accessibility and cytolytic activity-score (computed as the log-average of geometric mean of *GZMA* and *PRF1* TPM-expression),^30^ while plotting this correlation against the log_2_(fold-change) for putative immune-specific ATACseq peaks relative to TCGA-hits. We utilised this previous data for representation of only CD8^+^T cells-relevant gene-hits within SKCM or GBM TCGA-context.

### 10. Decision tree-based ensemble machine learning

The TCGA patient RNAseq data, processed by the TCGA PanCancerAtlas Immune Response Working Group^25^ (Kallisto, log-transformed TPM) was correlated to the CIBERSORT-LM22 CD8^+^T cell-fractions they predicted for the respective SKCM and GBM samples. Using the only the expression of csCD8^+^T cell-signature genes as input features, we predicted the CIBERSORT-LM22 CD8^+^T cell fractions as continuous response variable. To this end, we opted for a tree-based regression model based on gradient boosting, the XGBoost algorithm^79^. Performance estimation was performed using leave-one-out cross-validation (LOOCV) as evaluation. We used conservative parameters (learning rate=0.05, n_estimators=2000). Correlation values were obtained with a Pearson correlation between the predicted results and the reported CIBERSORT-LM22 CD8^+^T cell fraction computed from CRI iAtlas workflow.^25^ Models were trained for each cancer type separately. Afterwards, the full SKCM and GBM datasets were used to predict the CD8^+^T cell-fractions of the left-out cancer type (leave-one-group-out) to estimate how well the models extrapolate to the other cancer. The source code and brief explanation to reproduce the above machine learning exercise has been provided as **Supplementary-File.l.**

### 11. Tumour-derived single-cell RNA-sequencing (scRNAseq) mapping

#### a. Literature meta-analyses driven shortlisting of scRNAseq maps

To short-list suitable high-quality scRNAseq pan-tumour SKCM or GBM maps from existing peer-reviewed literature, we accessed the Tumor Immune Single Cell Hub (TISCH, http://tisch.comp-genomics.org),^35^ a large-scale curated database integrating single-cell transcriptomic data from >2 million single-cells (with a uniform/standardized workflow accounting for quality control and batch-effects) across >75 high-quality tumour-derived datasets (for 28 cancertypes). Herein, we used the Gene Exploration module of TISCH to short-list scRNASeq datasets with both cancer cells and innate/adaptive immune cell profiling based on both quantitative as well as qualitative coverage of the csCD8^+^T cell signature. This delineated 3 SKCM datasets (GSE115978, GSE72056, GSE120575) and only one GBM dataset (GSE131928). From this initial list of scRNAseq datasets it was amply clear that there was a big difference in CD8^+^T cells profiled from SKCM patients as compared to GBM patients across studies – resulting possibly from latter’s lymphocyte disparities. Henceforth, to allow at least some degree of comparative proportionality we decided to do further selection based on how closely SKCM-to-GBM ratios overlapped between the different scRNAseq datasets for CD8^+^T cells and TCGA datasets-derived immune-deconvolutions for lymphocytes, CD8^+^T cells and T_REG_. Based on this, we could delineate suitable SKCM^37^ and GBM^36^ scRNAseq datasets for further analyses and comparison.

#### b. Single-cell data pre-processing

Single-cell expression data (log_2_-transformed TPM counts, with a pseudocount of 1) obtained by SMART-Seq2, as well as the corresponding cell annotations were downloaded from the BROAD single-cell analysis data portal (https://singlecell.broadinstitute.org/single_cell) under identifiers SCP393 and SCP11 for GBM and SKCM, respectively. T cell labels were further enhanced using CIBERSORTx workflow^75^ (https://cibersort.stanford.edu) to distinguish between CD4^+^ and CD8^+^T cells. Correct annotation of these labels was checked using expression of standard FACS markers. All datasets received filtering for mitochondrial genes and quality trimming using the built-in function in STREAM^39^ (cal_qc()). Cells with less than 100 genes expressed were removed. Genes expressed in less than 10% of the cells per population were also discarded for further analysis. Post-filtering, 1180 CD8^+^ and 889 CD4^+^ T cells remained for SKCM respectively; while 52 and 42 cells remained for CD8^+^ and CD4^+^ T cells in GBM, respectively. To assess if both datasets were of similar quality, we calculated the QC metrics of the scRNAseq dataset with Scanpy and compared drop-outs (pct_dropout_by_counts) in the 3566 genes variable in both datasets. A Welch’s two-sided t-test indicated no biases in drop-out for these genes was detected (p=0.42).

#### c. Dimensionality reduction

The space was initialized using a two-dimensional PCA with only the LM22 genes as input. Manifold representations (UMAP^80^) and individual gene expression was plotted on top of this space for additional visual inspection of the dataset and the PCA separation of CIBERSORT labels.

#### d. The csCD8^+^T cell signature concordance/discordance analyses

CD8 T-cell normalized gene expression values were used to quantify the inference of the csCD8^+^T-cell signature in the expressed genes of the single cell data in GBM and SKCM. For each GBM and SKCM dataset, the Spearman correlation matrix was calculated with the “spearmanr” function from the Scipy Stats module, using the signature genes as the variables. The hierarchical clustering maps were created with Seaborn using Euclidean distance as metric. Underlying distances were transformed to a polar axis dendrogram with GBM and SKCM csCD8^+^T-Cell distance dendrogram plotted in opposite sites. For T cell-to-cell gene coverage/consistency analyses, nearest neighbourhood analysis (with Spearman rank correlation) was applied for applicable genes, using the analyses modules available within Broad Institute’s Single-Cell Portal workflow (https://singlecell.broadinstitute.org/single_cell).

#### e. T cells subpopulation annotation

We investigated the subpopulation characteristics of the T cells in SKCM and GBM datasets using SCANPY’s neighbourhood-based workflow. Neighbourhood graphs were defined with each cell having 10 neighbours. We applied PAGA^81^ to improve clustering results and then ran the recommended Leiden^82^ graph-clustering approach to aggregate individual T-cells in subpopulation structures. We observed more proficient connectivity between SKCM T cells than between GBM T cells, most likely due to limited diversity in the latter. The resulting graph structures were exported to JSON and further analysed with networkx^83^ and Cytoscape^84^. Since nomenclature of T cell subpopulations in scRNAseq datasets is somewhat of an issue, especially due to non-standardized usage of limited gene-sets for defining these subpopulations, we decided to take a comprehensive annotation approach accounting for well-established phenotypic as well as functional immunophenotyping markers (Publications;^41,42,85,86^ HCDM Database: http://www.hcdm.org/; Immuno-phenotyping Guide: https://assets.thermofisher.com/TFS-Assets/LSG/brochures/immune-cell-guide.pdf). Based on these sources, a comprehensive CD4^+^T cell or CD8^+^T cells-specific gene-sets **(Supplementary Table 2)** were uniformly applied across SKCM or GBM T single-cells and only the genes passing qualitative thresholds were used for final annotation of T cell subpopulations within the SCANPY Leiden-derived clusters (see **Supplementary Box 1** for final annotations).

#### f. Functional immune-annotation for trajectory (FIAT) approach for annotating T cells

Basic pseudo-time trajectories were inferred using the STREAM workflow. For STREAM pseudo-time trajectories, default package settings to generate the principal graphs and to analyse the different marker types (diverging, transition) were used. We used Z-scores to standardize gene expression of all genes identified by STREAM per branch, to the mean expression in the T-cell subpopulations. Z-scores were then ordered in descending order and used to run a pre-ranked GSEA with GSEApy (https://github.com/zqfang/GSEApy), using 2000 permutations for FDR estimation. Here, T cells-specific published immune-signatures from Immune-MSigDB datasets^21^ were used for GSEA. To annotate the branches, we filtered the Immune-MSigDB dataset signatures to include only signature sets with relevance for T-cell functional-phenotypes using keyword restrictions. Only datasets with combinations of terminologies like, “ACTIVATED”, “NAÏVE”, “EXHAUSTED”, “UNTREATED”, “RESTING”, “IL2”, “DELETIONAL_TOLERANCE”, “NONSUPPRESSIVE”, “STARVED”, “ACT”, “STIM”, “UNTREATED” on both sides of the signature descriptor, were the only T-cell signatures to be retained. Beyond this initial filtering, following 4 successive inclusion or exclusion criteria were ultimately applied for increased flexibility and clarity within the analyses workflow: (I) datasets with lack of experimental clarity on CD4^+^T cells or CD8^+^T cells were excluded → (II) only datasets with positive NES-values were selected (i.e., up-regulated genes enriching a dataset positively) → (III) only those datasets were selected that had FDR p-value <0.01, or <0.3, for SKCM or GBM respectively (in order to account for the disproportionalities in T cell-numbers between the two scRNAseq datasets) → (IV) lastly, only top-3 enriched datasets were selected from the list generated after step III to avoid unnecessary redundancies. Herein, branches that couldn’t meet any of these criteria and hence failed to enrich any Immune-MSigDB datasets were accordingly labelled “low functional”. Of note, both CD4^+^/CD8^+^T cell-specific datasets were indiscriminately used for annotations of either cell-type due to high functional overlap between these cell-types on almost every immunological level. The Immune-MSigDB datasets shortlisted at the end of step III are enlisted in Supplementary Table 3.

#### g. Ligand-receptor interactions

CellPhoneDB^44^ is a resource that contains a list of curated receptor-ligand interactions. Raw log_2_-normalized gene expression was used as input. We removed all cells that were not CD4^+^ or CD8^+^T-cells, macrophages or malignant/cancer cells. CellPhoneDB calculates the means for the interaction list and the p-values for the enrichment of ligand-receptor interaction pairs, based on the expression of the receptor in one cell type and the ligand in the other. Full CellPhoneDB hits are elaborated in Fig.S7. However, CellPhoneDB gives a rather large and redundant output that is pan-literature derived rather than tailored to a particular immunological context – altogether this reduces clarity to some extent. To overcome this hurdle we decided to apply a TCGA SKCM/GBM-driven systems biology-based prioritisation strategy to filter CellPhoneDB hits for CD4^+^::CD8^+^T cell interaction and CD4^+^/CD8^+^T cell::cancer cell interactions. More specifically, as initial input gene-list, we only selected those ligand or receptor-coding genes that were mentioned ≥2 times in CellPhoneDB hits unique to SKCM or GBM, and ≥3 times in CellPhoneDB hits shared between SKCM and GBM. The resultant gene input list was separately entered (for each cancer-type) into the WEB-based Gene SeT AnaLysis Toolkit (WebGestalt) work-flow,^87^ to execute human-specific network topology-based analyses (NTA) using TCGA-SKCM or TCGA-GBM RNAseq datasets, separately, for network building. The network was constructed in “network expansion” orientation (enriching top-25 ranking neighbors at cut-off of 0.05 FDR p-value). The resulting network file was processed for final representation via CytoScape workflow.

#### h. Cell cycle phase scoring

Cell phase labels were added with SCANPY’s score_genes_cell_cycle() function, which uses the 97 genes described in Tirosh et al., 2015.^37^ For each cell type, we calculated the proportions of cells in G1, G2M and S-phases. Significance of the cell cycle distribution was obtained by using Fisher-exact tests between the corresponding cell cycle phase proportions in GBM and SKCM with a 95% confidence interval. Ridge plots representing the expression of key genes, such as JAK3 and STAT1, in each of the cell cycle phases were generated using the JoyPy package (https://github.com/sbebo/joypy).

#### i. Difference in key gene expression

For statistical assessment of *JAK3* and *STAT1* in the cell-cycle, the log_2_-transformed TPM expression data from the BROAD institute were filtered to each cell population (CD4, CD8) per cancer type. We then filtered each subpopulation based on the suggested STREAM workflow (mitochondrial gene removal, quality calculation and sparsity cut-offs). Datasets were normalized, then merged with SCANPY’s ingest method. We then performed the construction of the elastic principal graph on the merged dataset. The root of the graphs was chosen as the point where GBM and SKCM cells showed most overlap, which was node S0. We then defined the pseudo-time as the distance from this tree root. To assess the difference in *JAK3* and *STAT1* expression, pseudotime from S0 was used for both the GBM and the SKCM samples. The pseudo-time was discretized in 10 equidistal bins and we then calculated the total area under the curve per bin, using trapezoid approximation. Co-variation was then tested with a linear model, using the time (as bin rank) and JAK3/STAT1 expression per bin and per cell cycle phase between both cancer types in R. Pairwise p-values were calculated using the emmeans R package^88^, based on least squares.

#### j. Significance of key regulators in exhaustion

Statistical significance of IL2, TNF, IFNG, CD28, PDCD1, TOX and EOMES between exhausted and activated states across cancer and cell types was assessed using the R package MAST^89^, which uses a custom hurdle model to compensate for drop-out biases. Statistical significance between cell cycle phases within a cancer type and cell type, regardless of pseudotime, were calculated using Welch’s two-sided t-test, as gene drop-out biases for a specific cell cycle phase may also hold information^90^. P-values in those cases were adjusted using Benjamini-Hochberg correction with alpha=0.05.

#### k. Expression profiling-based analysis of CD8^+^T cells

For the downstream analysis of CD8^+^T cell expression profiles, we normalized the dataset using a linear scaling factor normalization. CD8^+^ and CD4^+^T cells’ gene expression profiles from SKCM/GBM scRNAseq datasets were normalized by scaling factor of total cell counts (log_2_) with very highly expressed (> avg. 5%) genes excluded from the computation of the normalization factor (size factor) for each cell using SCANPY.

#### l. IL2 withdrawal-induced cell cycle exit analyses

The gene signature for “T cell-specific IL2-withdrawal induced cell-cycle exit” was derived from a published resource (Supplemental Table 2).^49^ The median of the gene expression averages for the genes in this signature for CD4^+^ and CD8^+^T cells in GBM and SKCM, was extracted and then used to calculate the Spearman correlation of each individual T cell in the dataset. The distribution of these Spearman correlation values was then used to perform the contrast between GBM and SKCM for their respective T cell types (CD4^+^ and CD8^+^T cells) using t-test.

#### m. Volcano plot and GSEA-based pathway enrichment

Using the CD8^+^T cell normalized expression data, differential gene-expression contrasts were performed between GBM and SKCM for each T cell type (CD8 and CD4) by t-test and Benjamini-Hochberg correction by multiple testing, and represented via volcano plots. Subsequently, using the GSEA API from BROAD Institute we calculated the enrichment of genetic sets and pathways for each differential gene-expression profile to enumerate REACTOME pathways. The normalized single cell expression matrix for each T cell type was used to perform the enrichment using the annotation of SCKM and GBM to calculate pathway enrichment scores using gseapy with default parameters and selecting for the ‘signal to noise’ method, and “phenotype” permutation type.

#### n. SCENIC-driven gene-regulatory network (GRN) analyses and integration with TCGA ATACseq

Initial regulatory network inference of transcription factor (TF)-gene targets was extracted using GRNBoost2 and Arboreto algorithms^91^ following pySCENIC suggested workflow with default parameters using feather file hg19-500bp-upstream-7species.mc9nr.feather, motifs file motifs-v9-nr.hgnc-m0.001-o0.0.tbl and a file of 1013 TFs (Supplementary Table 2).^92^ These results were used for regulon prediction using ‘pySCENIC CTX’ (cisTarget) to find enriched motifs for a gene signature and optionally prune targets from this signature based on cis-regulatory cues. Only regulons with TFs at 500-bp upstream of transcription start-site as well as with minimum 5 target genes and a NES of >2.5 were considered and their regulon specificity score (RSS; basis for final scatter plots) were calculated for the corresponding groups i.e., CD8^+^/CD4^+^T cells across SKCM/GBM scRNAseq-datasets using ‘pyscenic.rss’ function. For analyses exploiting overlap of TCGA ATACseq with scRNAseq-dataset derived regulons, the previous regulatory network inference of TF-gene targets inferred only for each independent CD8^+^ and CD4^+^T cells were used to quantify the genetic targeting. Then ATACseq identified genes from within the TCGA-dataset (see above) were used to filter the network target inferred TF-gene targets. Only TFs targeting 6-to-7 of the ATACseq target-genes were included in the final figure-representation. For each TF targeting multiple genes, only the highest inferred network importance score from the GRNBoost2 algorithm was used for colour-based scaling.

### 12. Mathematical modelling of CD8^+^T single-cell cycling

#### a. The stretched cell cycle model

To execute this workflow, we searched the literature for suitable high-quality research publications experimentally studying lymphocytee cell cycle kinetics and aiming to create mathematical models based on these characteristics. Herein, we found that Dowling *et al.* measured the duration of the different cell cycle phases for primary murine B and CD8^+^T lymphocytes, activated *in vitro* to mimic the immune responses,^50^ and based on these data they proposed the so-called *stretched cell cycle model,* which provides a mathematical description of cell cycle progression for proliferating lymphocytes. By stimulating CD8^+^T cells with the T-cell receptor (TCR)-stimulus αCD3 (145-2C11) and interleukin-2 (IL-2), they found that the cell cycle duration of dividing cells varies considerably, having a mean duration of approx. 13 h with a standard deviation of about 3 h.^50^ Using the data of these CD8^+^T cells, they were able to determine parameters for the *stretched cell cycle model* that accurately describe the experimental observations.^50^ Essentially, the model assumes that all cell cycle phases occupy approximately fixed proportions of the total cell division time. In other words, longer cell cycle durations are assumed to stretch the duration of all cell cycle phases.^50^ In the original publication, experimental data of total cell cycle duration was fitted using a lognormal distribution and then “stretched” to obtain the cell cycle phase distributions.^50^ The stretched cell cycle model consists of the following parameters:

*μ* = 12.95 *h*
*σ* = 3.46 *h*
*k_s_* = 0.51 + 0.05*ξ*
*k*_*G*2-*M*_ = 0.14 + 0.02*ξ*
*k*_*S–G*2-*M*_ = *k_s_* + *k*_*G*2-M_

Where *μ* represents the mean cell cycle duration, *σ* its standard deviation, *k_s_* the fraction of time spent in S phase, *k*_*G*2-*M*_ the fraction of time spent in G2-M phase, and f a random number drawn from a normal distribution. From these parameters, the total cell cycle duration and duration of the cell cycle phases are then determined as follows:

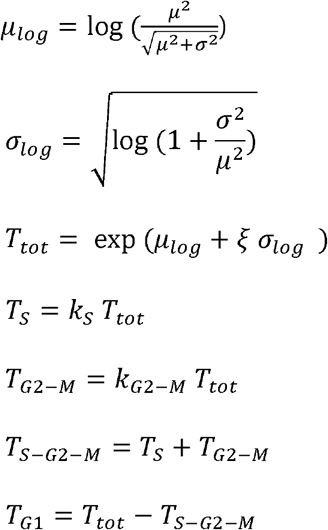

We used the scRNAseq data-derived cell cycle phase-scores for SKCM/GBM CD8^+^T cells (described above) as input for this stretched cell cycle model to derive pseudo-duration kinetics. More specifically, using this model, and the experimentally measured cell cycle durations for normally proliferating CD8^+^T cells,^50^ we adjusted two model parameters (*k_S_*, *k*_*G*2-*M*_) to reflect the cell cycle phase score-proportions computed for SKCM and GBM CD8^+^T single-cells. For SKCM CD8^+^T cells, we used (*k_S_*, *k*_*G*2-*M*_) = (0.325 + 0.05*ξ*, 0.288 + 0.02*ξ*), while for GBM cells we used (*k_S_*, *k*_*G*2-*M*_) = (0.115 + 0.05*ξ*, 0.096 + 0.02*ξ*). This resulted in cell cycle phase pseudo-duration estimations.

#### b. The Erlang cell cycle model

In the publication describing this model,^51^ the authors use quantitative measurements of cell cycle dynamics in human cells to propose a theoretical model whereby cell cycle progression in single cells is a succession of uncoupled, memoryless phases, each composed of a characteristic rate and number of steps.^51^ The authors note that the durations of G1, S, and G2 phases are each well captured by a so-called Erlang distribution, which describes a process that consists of a cascade of *k* independent reactions that occur at a rate λ.^51^ The durations of G1 and G2 phases were found to have a wider distribution consistent with a small amount of slower reactions, while S phase duration is characterized by a narrow distribution consistent with a large amount of faster reactions.^51^ Once the number of reactions *k* and the rate λ has been characterized, the model allows predicting distributions of cell cycle phase durations.

Chao *et al*. interpreted these reaction steps (*k*) and reaction rates (λ) in different ways:

Using a so-called *One for all* model,^51^ the reaction rates are determined by one general control system. This is like the stretched cell cycle model as one single control parameter will scale all cell cycle phase durations. We used this model to fit the reaction rates λ to the mean cell cycle phase duration of CD8^+^T cells from Dowling et al. paper.^50^ This gave us the following parameter set:

G1 phase: k = 10, λ = 2.22
S phase: k = 20, λ = 3
G2-M phase: k = 10, λ = 5.57

The results of this model using these parameters were depicted as gray dots in final figure, where the blue square indicates the mean of the distribution of cell cycle phase pseudo-durations.

Chao *et al.* also introduced a so-called *One for each* model, where rather than having one factor that controls all cell cycle phase durations, there are factors that specifically control the duration of specific cell cycle phases. Such a control system thus no longer ensures that all phase durations stretch by the same amount. One factor can thus only influence the reaction rates in one phase, changing the relative cell cycle phase distribution. The gray region in final figure shows all possible locations a cell can have in the phase space spanned by the fraction of time spent in G1, S and G2-M (the sum of the fractions needs to be 1). Starting from the parameter set obtained for the proficiently proliferating normal CD8^+^T cells from Dowling et al. paper,^50^ the cell cycle-phase scores for SKCM-CD8^+^T single-cells are obtained if the reaction rates λ in G2-M phase and in G1 phase are decreased by a factor of 2:

G1 phase: k = 10, λ = 1.11
S phase: k = 20, λ = 3
=G2-M phase: k = 10, λ = 2.785

Similarly, by gradually decreasing the reaction rates λ in G1 phase further, up to five-fold, a distribution is obtained that is consistent with the relative cell cycle phase pseudo-durations for GBM-CD8^+^T single-cells:

G1 phase: k = 10, λ = 1.11
S phase: k = 20, λ = 3
G2-M phase: k = 10, λ = 0.557

### 13. MILAN analyses of our SKCM and GBM patient cohorts

#### a. Tissue staining

Multiplex immunohistochemistry was performed according to the previously published method.^19,93^ Tissue sections of 3 μm were prepared out of formalin-fixed paraffin-embedded (FFPE) human SKCM and GBM tumour-samples (collected at the UZ Leuven/KU Leuven biobank according to the following protocols: S59804, S61081 and S62248). Tissue sections were dewaxed by sequential incubation steps in xylene and ethanol baths. Afterwards, antigen retrieval was performed in Tris buffer pH 8 with 10 mM EDTA (PT link, Agilent). Immunofluorescent staining was performed using the Bond RX Fully Automated Research Stainer (Leica biosystems) with the primary antibodies as indicated in Supplementary Table 5. The sections were incubated during 4 h with the primary antibodies, washed and visualized with secondary antibodies (30 min incubation; see Supplementary Table 5). A coverslip was placed on the slides with medium containing DAPI and slides were scanned in either a Zeiss Axio Scan Z.1 (Zeiss) with 10x magnification (GBM; 0.66 micrometers/pixel or μm/px) or a Hamamatsu NanoZoomer at 20x resolution (SKCM; 0.44 μm/px). After completion of the scanning procedure, the coverslips were manually removed, following by stripping of the antibodies in a buffer containing 1% SDS and β-mercaptoethanol during 30 min at 56°C. After this stripping process, the slides were washed in washing buffer for 45 min with frequent changes of the buffer. The staining procedure was thereafter repeated until all markers were stained and scanned on each of the slides.

#### b. Image processing

*For SKCM:* Image analysis was performed in Fiji/ImageJ following the procedure described previously.^19^ Briefly, images from sequential staining rounds were aligned using the DAPI channel. Tissue autofluorescence was acquired every round in a dedicated channel and subtracted from the measuring channels. TMA images were segmented into individual cores using a custom macro. Core segmentation was followed by cell segmentation and single-cell measurements which were performed using the EBImage R package.^94^ For every cell, the extracted features included: X/Y coordinates, nuclear size, and Mean Fluorescence Intensity (MFI) for all measured markers. *For GBM:* Images analysis was performed in R. Briefly, images from sequential staining rounds were aligned using the DAPI channel. A baseline to estimate tissue autofluorescence was acquired in between rounds and subtracted from the respective marker measurements. Scenes from sequential rounds were aligned using a custom pipeline. Cell segmentation and single-cell measurements were performed using the EBImage R package.^94^ For every cell, the extracted features included: X/Y coordinates, nuclear size, and Mean Fluorescence Intensity (MFI) for all measured markers.

#### c. Data analysis

*For SKCM:* MFI values were normalized to Z-scores as recommended by Caicedo JC, et al.^95^ Z-scores were trimmed in the [0, 5] range to avoid a strong influence of any possible outliers in downstream analyses. Single cells were mapped to known cell phenotypes using three different clustering methods: PhenoGraph,^96^ FlowSom,^97^ and KMeans. Clustering was performed exclusively in a sub*set* of the identified cells (25K) selected by stratified sampling and using the markers defined as phenotypic: CD3, CD4, CD68, CD8, MELANA, and S100B. For each clustering method, clusters were mapped to known cell phenotypes following manual annotation from domain experts (FMB, YVH, double blinded). If two or more clustering methods agreed on the assigned phenotype, the cell was annotated as such. If all three clustering methods disagreed on the assigned phenotype, the cell was annotated as “other”. For each phenotype, a fingerprint summarising the average expression of each marker for all the cells of the given phenotype was constructed. These fingerprints were used to predict the phenotype of all the cells included in the dataset (minimum of Euclidean distance). *For GBM:* MFI values were normalized to Z-scores as recommended by Caicedo JC, et al.^95^ Z-scores were trimmed in the [0, 5] range to avoid a strong influence of any possible outliers in downstream analyses. Single cells were mapped to known cell phenotypes using manual gating on the individual markers. *For all samples*: We further dissected the tissue samples in three different areas based on tumoral cells. To that end, tissues were fragmented into 50×50 px tiles (~22 sq μm). Tiles with at least one cell identified as tumour were initially defined as cellular tumour areas. To reduce the impact of potential outliers a median filter was applied to the obtained tumoral masks. The invasive-margin was defined at 300 px (~132 μm) from the cellular tumour edge. The rest of the tissue was defined as leading-edge (non-tumoral area). Similarly, we defined vascular areas. In this case however, given that CD31 (marker for endothelial cells) did not show a strong overlapping with DAPI (marker used to segment single-cells), we defined vascular areas using a CD31 mask. Then, we identified perivascular areas at 100 px (~44 μm) from vascular areas. The rest of the tissue was defined as a non-vascular area. For GBM specifically, necrotic areas were further dissected in haemorrhagic tumoral areas or viable tumoral areas based on manual annotation. Necrotic (dry-necrotic) areas were also manually annotated and excluded from the analysis.

#### d. T cell activation and nearest neighbour analysis

We evaluated the activation of each CD8^+^T cell (CD3+CD8+) in the tissue following the methodology described previously^19^ which assigns an activation score in the [-1, 1] range to each CD8^+^T cell. Resulting activation scores were normalized for each sample using a Z transformation. Z-scores were trimmed in the [-3, 3] range to avoid too strong influence from outliers. Nearest neighbour analysis was performed for the CD4^+^T cell and CD8^+^T cell populations by calculating the distance to the closest contrary cell, that is, the closest CD8^+^T cell for each CD4^+^T cell and vice-versa. We calculated how the distance to the closest CD4^+^T cell affects the phenotypic activation of CD8^+^T cells. To that end, we calculated the average phenotypic activation score of all CD8^+^T cells with a closest distance smaller than a given threshold. We spawned all thresholds between 10 and 1000 μm. At every given threshold, we also calculated the difference in phenotypic activation between CD8^+^T cells that have a CD4^+^T cell closer than the threshold versus the ones that do not have any, and assigned a significance value using a statistical t-test. To see the real significance of this approach we compared the obtained p-value curve with that obtained with random activation scores obtained from 100 random permutations of the real activation scores.

### 14. Spatial-transcriptomics analyses in IVY Glioblastoma Atlas Project (GAP) dataset

Gene expression data (z-score normalised) as well as the corresponding patient and tumour anatomical-zone annotations were accessed from the IVY-GAP data portal.^56^ The procured data was utilised to chart a contour plot using the Plotly Chart Studio (https://plotly.com/chart-studio/). Other analyses were executed in the same manner as mentioned above for TCGA-datasets.

### 15. Cross-sectional tumour analyses from immuno-oncology clinical trials

Cross-sectional tumour transcriptomic data from SKCM^57,59^, BLCA,^58^ and GBM^6^ immunooncology clinical trials’ data was accessed using a standardized TIDE data-portal.^11^ Briefly, we accessed the prognostic impact of specific biomarkers (detected in pre-treatment tumour-tissue) on patient OS, after immune-checkpoint blocking (ICB) immunotherapy. Herein, the prognostic impact was computed as z-score derived from Coxph statistical modelling associated with corresponding p-values.^11^ These values for the different clinical trials and biomarkers were pooled to create a dot-plot. In some cases, information for certain biomarkers wasn’t available within the clinical trial’s published data and hence couldn’t be included in this analysis. Of note, although the GBM^6^ immuno-oncology clinical trial used above did retrospective analyses of longitudinally collected tumour transcriptomic data, we only utilized pre-treatment tumour-tissue data herein, in order to compare with above SKCM^57,59^ or BLCA^58^ clinical trials (wherein post-treatment tumour transcriptomic data wasn’t available). This GBM trial wasn’t included in the below longitudinal tumour analyses since its trial design wasn’t based on neoadjuvant anti-PD1 immunotherapy in resectable GBM, and its overall cohort size was smaller than below longitudinal GBM trial. Also, the transcriptomic profiling platforms between these two GBM clinical trials were completely different thereby not allowing us to uniformly pool and normalize the data together.

The data for cross-sectional immuno-oncology clinical trial that detected non-synonymous somatic TMB in pre-treatment tumour tissue (filtered for: cutaneous melanoma, malignant uveal melanoma or glioblastoma multiforme) and followed-up on patient OS (in months) after ICB-immunotherapy, was accessed from the cBioPortal workflow (TMB and Immunotherapy, MSKCC, Nat Genet 2019 dataset accessible at the following link: https://www.cbioportal.org/study/summary?id=tmb_mskcc_2018).^7^ In case of SKCM, the patient data was subdivided into following subgroups, depending on the primary or metastatic-site of resection/biopsy: primary SKCM-tumour (sample type: primary), in-transit SKCM-metastasis, lymph nodes SKCM-metastasis (regional, non-regional or unspecified), nonlymphoid organ/site associated SKCM-metastasis (liver, muscle, pleura, parotid, adrenal gland, spine, ovary, lung, stomach, axilla, pelvis, abdomen, epidural, small bowel, soft tissue) and brain SKCM-metastasis. The Pearson correlation between overall survival and pre-treatment TMB was calculated per tissue location of the metastases for SKCM. For the other cancer types, tissue of origin was used when metastatic information was not present. UVM data was metastatic.

### 16. Longitudinal tumour analyses from immuno-oncology clinical trials

Gene-expression and patient survival or clinico-pathological data from longitudinal clinical trials involving resectable-SKCM or resectable-GBM patients treated with neoadjuvant anti-PD1 immunotherapy,^60,61^ were accessed from published sources. In case of SKCM, the patient survival data wasn’t publicly available and thus we were kindly provided access to this data by lead investigators of that study, Dr. Alexander Huang and Dr. Tara Mitchell (UPenn, USA). Since both these clinical trials utilized NanoString nCounter System for tumour transcriptome profiling, which is limited in terms of number of genes profiled as compared to tumour bulk-RNAseq, we pursued some adjustments for our genetic-signature analyses. This limitation did not affect analyses for lymphocytes infiltration signature or CD8^+^T cell-signature. However, for csCD8^+^T cell-signature, while expression of *B3GAT1* and *TOX* was available in SKCM-dataset yet these genes weren’t available in the GBM-dataset, hence for the sake of consistency, these two genes were excluded from the csCD8^+^T cell-signature. Despite this adjustment, most of our results were highly similar to results obtained with TCGA-dataset with the full gene set, thereby showing that the above adjustment didn’t compromise our workflows. Similarly, in order to derive a TGFβ/wound healing response signature tailored to these clinical datasets, we delineated the genes overlapping between the SKCM/GBM NanoString-datasets and the Gene-Ontology (GO) Wound Healing gene-set (GO:0042060), after that GO-term was significantly enriched by the full SKCM/GBM NanoString dataset-derived gene lists, using the STRING work-flow (for SKCM, 61 overlapping genes with *p=4.43·10-^13^; for GBM, 76 overlapping genes with *p=6.03·10-^23^).^98^ Thereafter we selected common genes between both SKCM and GBM datasets for the sake of analytical consistency (a total of 30 genes) to create the signature **(Supplementary Table 2).** This adjustment did not compromise the immunological veracity of this signature since several major wound healing-associated immunosuppressive or immunoregulatory protein-coding genes (e.g., *ADORA2A, AXL, MERTK, CD9, CEACAM1, CX3CL1, ENTPD1, GATA3, IL6*) and TGFβ-related gene (*TGFB1*) were in fact integrated in this signature^32^ thereby substantiating its use herein. All the analyses with these datasets, followed the same methodologies as for the TCGA datasets described above. Similar approaches (as applicable) were applied to longitudinal settings with DC-vaccines in GBM-patients,^64^ as well as GBM-patients infused with adoptive T cell therapy (ACT) products that underwent transcriptomic profiling.^63^ In the latter case, we defined GBM clinical responders to ACT as patients experiencing no tumour progression or time-to-tumour progression (TTP) of ≥8 months whereas GBM clinical non-responders to ACT were those experiencing TTP <8 months. The threshold of 8 months was defined based on a 75th quartile cut-off for the continuous distribution of all TTP values across the entire ACT clinical trial dataset. The data for this ACT clinical trial was kindly provided by the lead investigators of this study, i.e.: Dr. Corey Smith, Dr. Sriganesh Srihari, and Dr. Rajiv Khanna (QIMR Berghofer Medical Research Institute, Australia).

### 17. TIDE model-based immunotherapeutic predictions

For ICB-immunotherapy target predictions, we used the TIDE computational model^11^ on TCGA-SKCM and TCGA-GBM cohorts pre-loaded within the “Query gene” module of TIDE web portal. Through this module, we derived the TIDE score (also called T cell dysfunction score),^11^ which is basically a z score, whose positive value indicates a good ICB-immunotherapy (query gene) target i.e., significantly positive z-score for a queried gene indicates that when the queried gene’s expression is low (“mimicking” therapeutic blockade), this results in positive prognostic impact of CD8^+^T cell-signature whereas this pattern is disrupted or non-discernible when the queried gene’s expression is high.

### 18. Statistical analyses

The statistical details of all the analyses are reported in the figure legends and/or figures, including statistical analysis performed, statistical significance thresholds/values and in most cases the counts/number of data-points. In some cases, the counts/number of data-points are described in the methods section due to space limitations in figure legends. All the statistical tests used herein were always two-tailed unless otherwise explicitly mentioned. Gene signatures were estimated by considering the average expression of all the genes within that signature, unless otherwise mentioned. All statistical analyses or graphical representations were executed using Python version 3.7.3, R versions 4.0.1, 3.6.2, and 3.5.3 or GraphPad Prism version 8, as feasible or applicable. Different package versions used in this manuscript are detailed in **Supplementary Table 6.**

