## Supplemental figures for "Immunogenomic, single-cell and spatial dissection of CD8^+^T cell exhaustion reveals critical determinants of cancer immunotherapy"

[1]Ludwig Institute for Cancer Research, Brussels, Belgium; [2]De Duve Institute, UC Louvain, Brussels, Belgium; [3]Laboratory of Cell Stress & Immunity, Department of Cellular & Molecular Medicine, KU Leuven, Belgium; [4]Laboratory for Precision Cancer Medicine, Translational Cell and Tissue Research, Department of Imaging & Pathology, KU Leuven, Belgium; [5]Laboratory of Experimental Oncology, Department of Oncology, KU Leuven, Belgium; Department of General Medical Oncology, UZ Leuven, Belgium; [6]Laboratory of Dynamics in Biological Systems, Department of Cellular & Molecular Medicine, KU Leuven, Belgium; [7]Laboratory of Translational Genetics, Department of Human Genetics, KU Leuven, Belgium; [8]VIB Center for Cancer Biology, VIB, Leuven, Belgium; [9]Laboratory of Cell Death Research & Therapy, Department of Cellular & Molecular Medicine, KU Leuven, Belgium; [10]Laboratory of Tumor Microenvironment and Therapeutic Resistance, Department of Oncology, VIB-Center for Cancer Biology, KU Leuven, Leuven, Belgium; [11]Department of Neurological Surgery, UCSF Comprehensive Cancer Center, UCSF, San Francisco, CA, United States; [12]VIB Center for Brain and Disease Research, Leuven, Belgium; [13]Department of Microbiology and Immunology, KU Leuven, Belgium; [14]Laboratory of Lymphocyte Signalling and Development, The Babraham Institute, Cambridge, UK; [15]Department of Neurosurgery, University Hospitals Leuven, Leuven, Belgium; [16]Laboratory of Experimental Neurosurgery and Neuroanatomy, Department of Neurosciences, KU Leuven, Belgium; [17]Leuven Brain Institute (LBI), Leuven, Belgium; [18]Department of Immunology and Oncode Institute, Leiden University Medical Center, Leiden, Netherlands; [19]Translational Cell & Tissue Research, Department of Imaging & Pathology, KU Leuven, Belgium; [20]Laboratory for Molecular Digestive Oncology, Department of Oncology, KU Leuven, Belgium; [21]These authors had equal contributions;

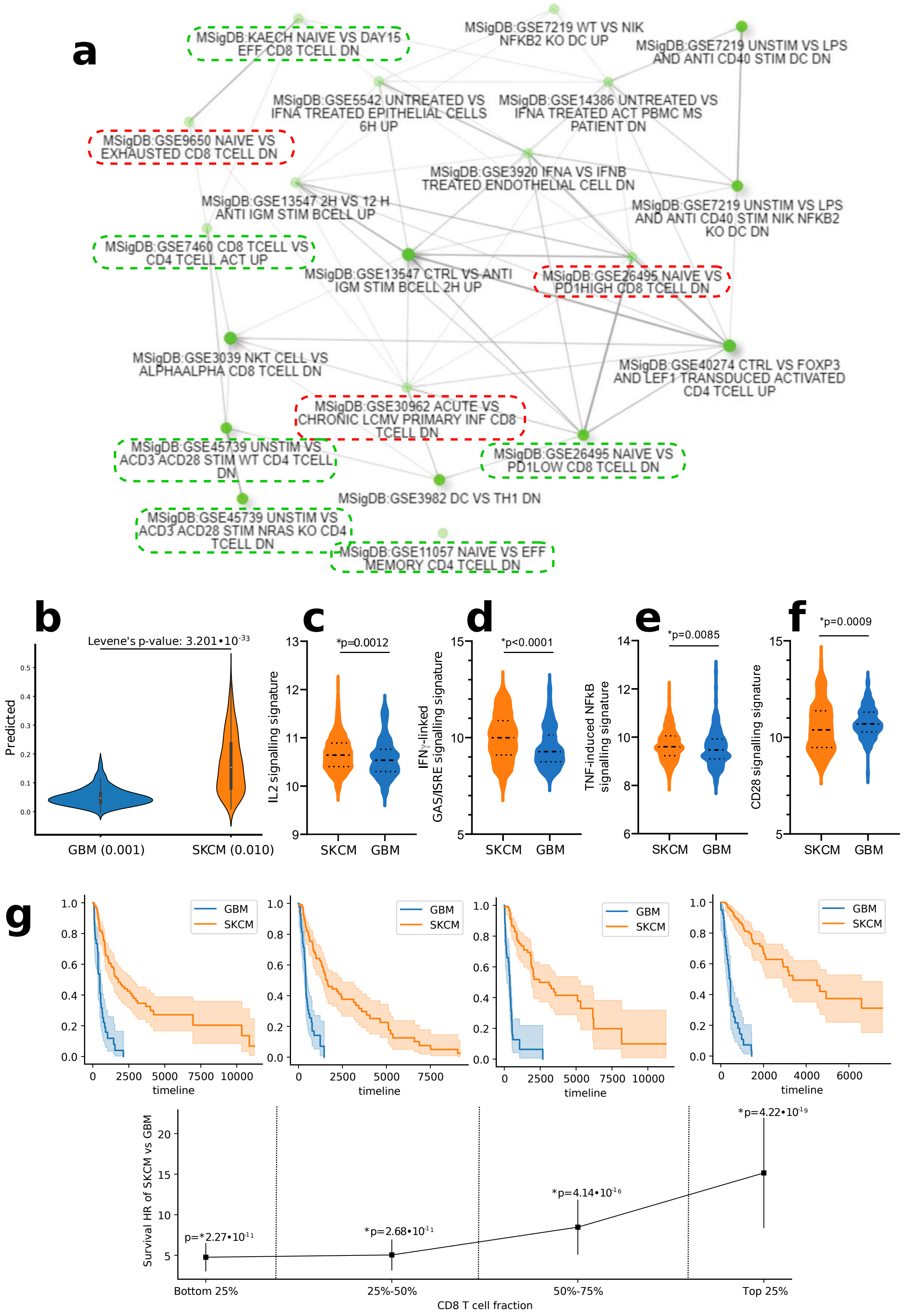

a
MSigDB:KAECH NAIVE VS DAY15 EFF CD8 TCELL DN
MSigDB:GSE7219 WT VS NIK NFKB2 KO DC UP
MSigDB:GSE7219 UNSTIM VS LPS AND ANTI CD40 STIM DC DN
MSigDB:GSE5542 UNTREATED VS IFNA TREATED EPITHELIAL CELLS 6H UP
MSigDB:GSE14386 UNTREATED VS IFNA TREATED ACT PBMC MS PATIENT DN
MSigDB:GSE9650 NAIVE VS EXHAUSTED CD8 TCELL DN
MSigDB:GSE3920 IFNA VS IFNB TREATED ENDOTHELIAL CELL DN
MSigDB:GSE13547 2H VS 12 H ANTI IGM STIM BCELL UP
MSigDB:GSE7219 UNSTIM VS LPS AND ANTI CD40 STIM NIK NFKB2 KO DC DN
MSigDB:GSE7460 CD8 TCELL VS CD4 TCELL ACT UP
MSigDB:GSE13547 CTRL VS ANTI IGM STIM BCELL 2H UP
MSigDB:GSE26495 NAIVE VS PD1HIGH CD8 TCELL DN
MSigDB:GSE3039 NKT CELL VS ALPHAALPHA CD8 TCELL DN
MSigDB:GSE40274 CTRL VS FOXP3 AND LEF1 TRANSDUCED ACTIVATED CD4 TCELL UP
MSigDB:GSE30962 ACUTE VS CHRONIC LCMV PRIMARY INF CD8 TCELL DN
MSigDB:GSE45739 UNSTIM VS ACD3 ACD28 STIM WT CD4 TCELL DN
MSigDB:GSE26495 NAIVE VS PD1LOW CD8 TCELL DN
MSigDB:GSE3982 DC VS TH1 DN
MSigDB:GSE45739 UNSTIM VS ACD3 ACD28 STIM NRAS KO CD4 TCELL DN
MSigDB:GSE11057 NAIVE VS EFF MEMORY CD4 TCELL DN
b
Levene's p-value: 3.201•10-33
Predicted
GBM (0.001)
SKCM (0.010)
c
*p=0.0012
IL2 signalling signature
SKCM GBM
d
*p<0.0001
IFNγ-linked GAS/ISRE signalling signature
SKCM GBM
e
*p=0.0085
TNF-induced NFkB signalling signature
SKCM GBM
f
*p=0.0009
CD28 signalling signature
SKCM GBM
g
GBM
SKCM
timeline
Survival HR of SKCM vs GBM
p=*2.27•10-11
*p=2.68•10-11
*p=4.14•10-16
*p=4.22•10-19
Bottom 25%
25%-50%
50%-75%
Top 25%
CD8 T cell fraction

**Supplementary Figure S1. (a)** A network depicting top 20 human Immune-MSigDB datasets enriched by the csCD8$^+$T cell-signature. The nodes connect the different datasets if ≥20% genes are shared between them. Darker nodes are more significantly enriched (FDR adjusted p-value<0.05), bigger nodes represent datasets with higher number of genes and thicker edges indicate higher overlapping genes. Green rectangles indicate effector/memory lymphocytes-related datasets and red rectangles indicate lymphocyte exhaustion-related datasets. **(b)** Distribution of AI/ML-model predicted values in TCGA-GBM and TCGA-SKCM datasets. The variance (between brackets) in GBM predictions is ten times less than in SKCM. This difference is statistically highly significant (Levene's test). These results connect to the plot in **Figure 1g**, wherein LOOCV performance indicates that the GBM-specific model is worse at predicting the trend (Pearson correlation between csCD8$^+$T-cell signature vs. CD8$^+$T cell-fractions predictions) in TCGA-GBM samples than the corresponding SKCM-specific model in TCGA-SKCM samples, exemplified by the substantially lower-variability in GBM-model. This is because, the low variance in GBM predictions as compared to SKCM means that the GBM-model is performing worse than SKCM; because an AI/ML-model that is successfully "learning" and generating reliable predictions tends to have higher variance owing to the natural variations of a dynamic patient population (e.g., SKCM) while lack of any meaningful predictions will lead to a minimally variable output (e.g., GBM). Feature weights also do not transfer well between both cancer types (**Figure 1g**, green-line). There is a statistically significant difference in mean prediction error (Welch t-test, p-value=$7.14*10^{-5}$), however, the effect size of this is small (Cohen's D, 0.310). The GBM model does not underfit the training data (training set Rp=0.99) and is responsive to changes in input features (test set Rp=0.03 using scrambled features). **(c-f)** Violin-plot of indicated signature's expression levels in TCGA-GBM and TCGA-SKCM datasets (Mann-Whitney test). (g) Analyses of CD8$^+$T cell-fractions' quadrant analysis: all quartiles were split off, based on the CD8$^+$T cell-fraction values arranged from lowest to highest 25% patient subgroups, inferred by CIBERSORT per cancer-type, individually. Per quartile, the Kaplan-Meier curves were plotted. The multi-variate CoxPh hazard ratios for GBM vs SKCM were estimated for each quartile of CD8$^+$T cell-fraction values. Features were z-score standardized. Statistical-significance threshold was *p<0.05. For different gene-signatures, see **Supplementary-Table.2**.

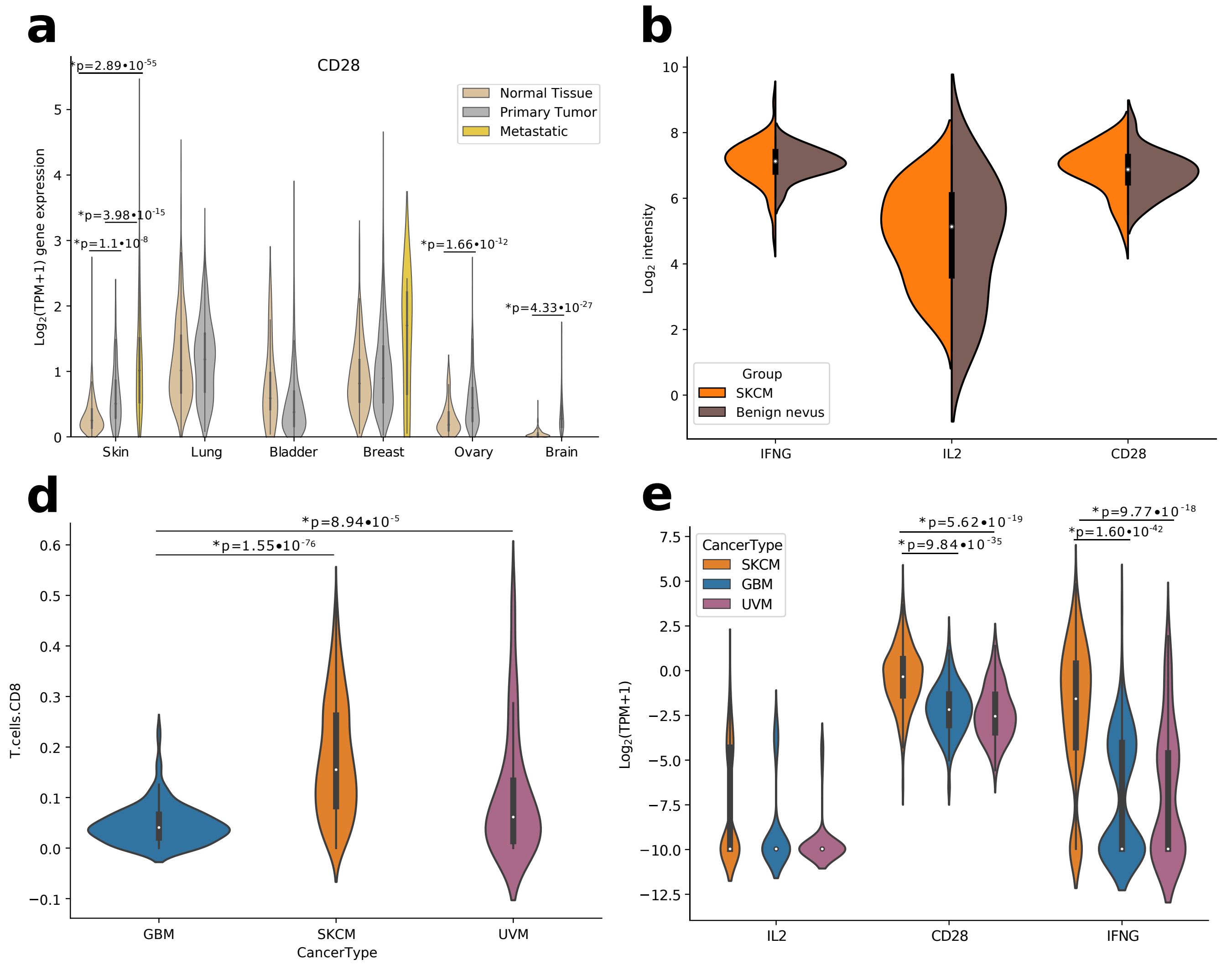


**Supplementary Figure S2. (a)** Violin-plots comparing *CD28* expression between TCGA primary-tumour/metastasis bulk-RNAseq vs. Genotype-Tissue Expression (GTEx) normal-organ/tissue bulk-RNAseq datasets (Welch's t-test). Brain (normal-tissue, n=207; primary-tumour, n=153), bladder (normal-tissue, n=9; primary-tumour, n=407), breast (normal-tissue, n=179; primary-tumour, n=1092; metastatic-tissue, n=7), lung (normal-tissue, n=288; primary-tumour, n=513), ovary (normal-tissue, n=88; primary-tumour, n=419), skin (normal-tissue, n=557; primary-tumour, n=102; metastatic-tissue, n=366). **(b)** Violin-plots comparing expression of indicated genes between primary-melanoma (n=45) and benign nevi (n=18). Violin-plots comparing CD8$^{+}$T cell-fractions **(c)**, *IFNG-IL2-CD28* **(d)** expressions between TCGA SKCM (n=470), GBM (n=171) and UVM (n=79) bulk-RNAseq datasets (Welch's t-test). Statistical-significance threshold was *$p<0.05$. For different gene-signatures, see **Supplementary-Table.2**.

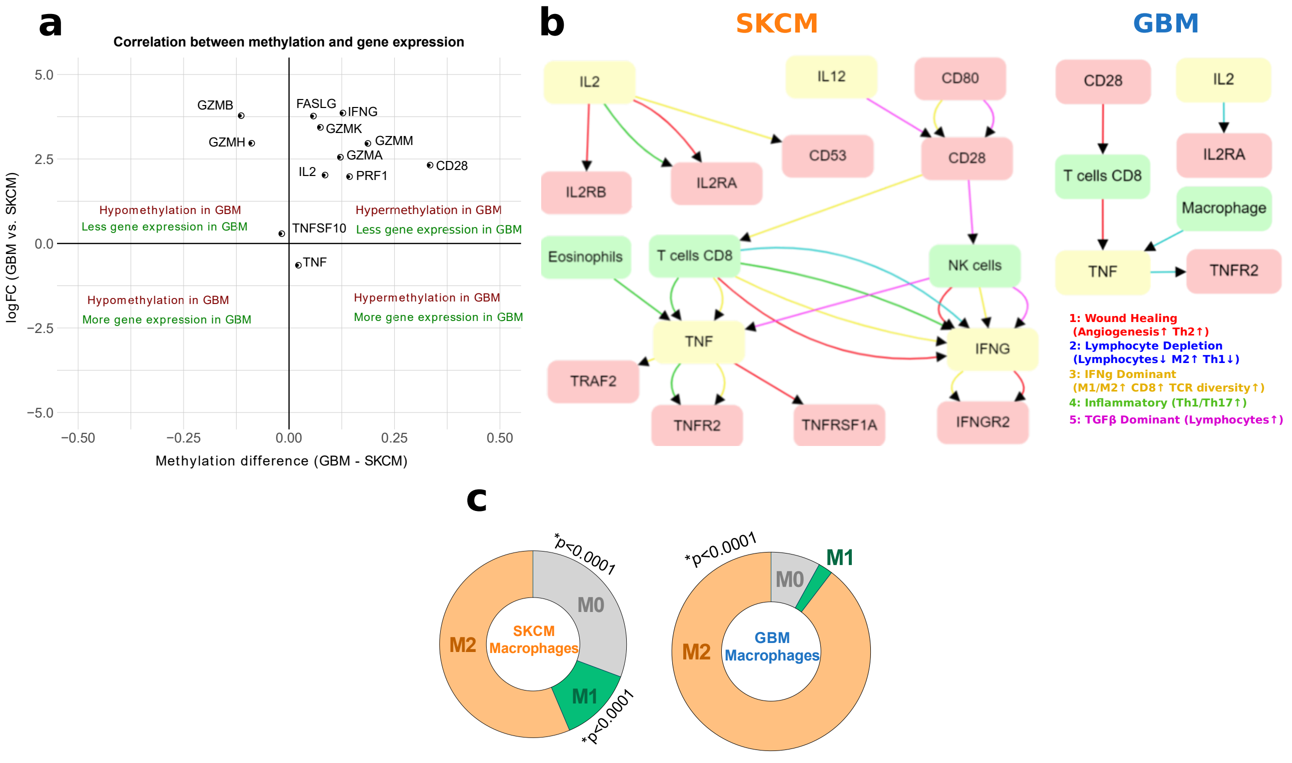


**Supplementary Figure S3. (a)** Scatterplot illustrating the relation between DNA methylation and gene-expression patterns for the indicated genes within TCGA-SKCM/GBM datasets. The X-axis shows the shift in methylation ß-value, while the Y-axis represents the expression-ratio between GBM and SKCM patients. **(b)** Extracellular network-analyses[1] driven by interaction between $CD8^+$T cell-fractions and *IFNG-IL2-CD28-TNF* depicting interactions between other "recruited" extracellular signalling ligands/cytokines and CIBERSORT-LM22 immune-deconvoluted immune cell fractions[1] within TCGA-SKCM or TCGA-GBM patient-datasets. Herein, the nodes indicate receptors, ligands or cell-types whereas the edges indicate specific enrichment of tumour immune-landscape features across specific patient subsets. Overall, these analyses indicate subset of patients enriching for particular tumour immune-landscape pathway and embedment of particular ligand-receptor-cell interactions within that landscape in those patient-subsets.[1] **(c)** Doughnut-plots representing average CIBERSORT-LM22 M0, M1 or M2 macrophage-fractions[1] in TCGA-SKCM or TCGA-GBM. Statistical estimate was done for SKCM vs. GBM comparison (Kruskal-Wallis ANOVA test with Dunn's multiple comparisons test). Statistical-significance threshold was $*p<0.05$. For different gene-signatures, see **Supplementary-Table.2**.

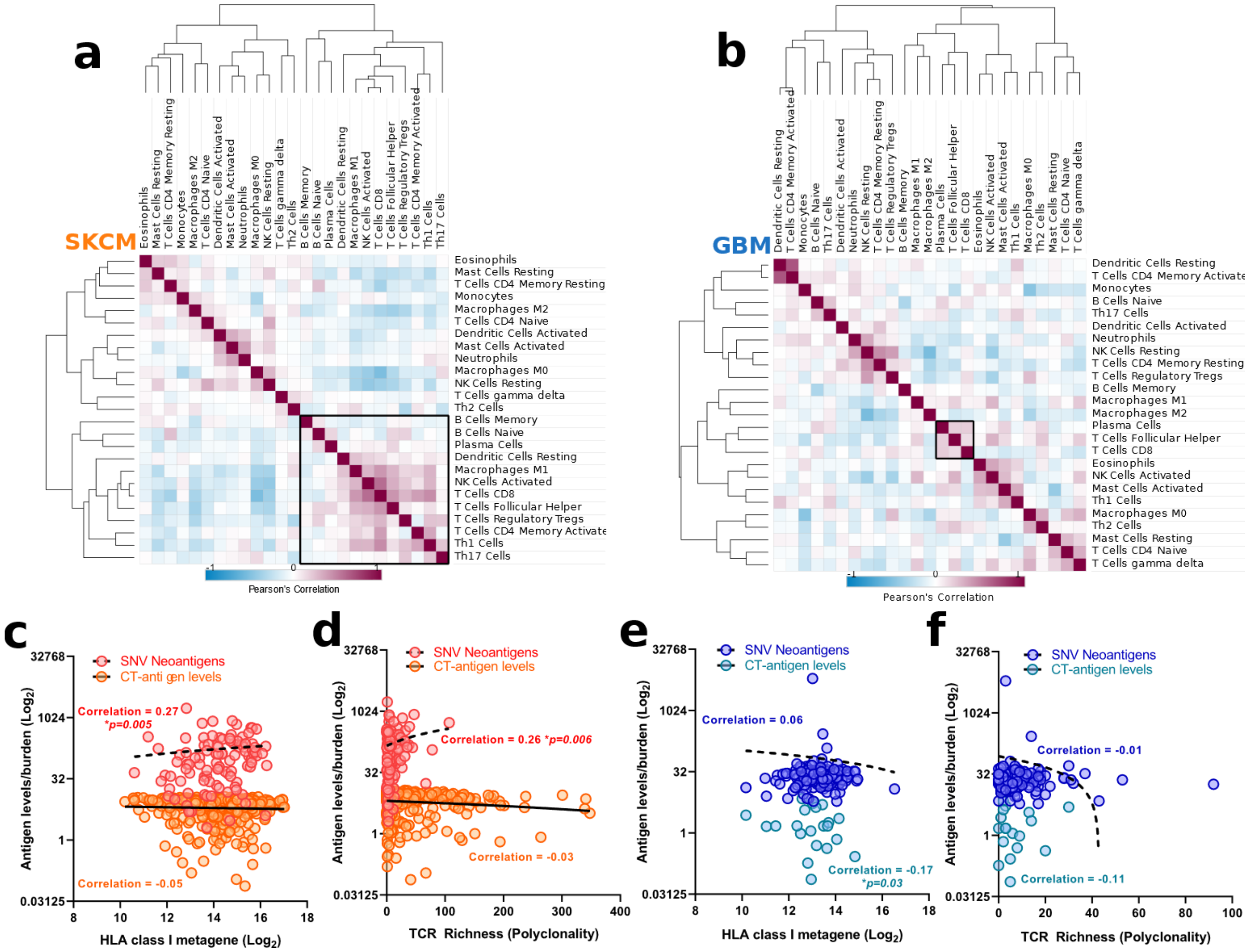


**Supplementary Figure S4. (a,b)** Correlation-matrices for different CIBERSORT-LM22 or CRI iATLAS based immune-deconvolution derived immune-cell fractions[1] across all TCGA-SKCM **(a)** or TCGA-GBM **(b)** patients. Spearman's correlation between indicated antigen-burdens (i.e., SNV-neoantigens or CT-antigen signature) and HLA class-I signature **(c,e)** or T-cell receptor (TCR) Richness **(d,f)** across TCGA-SKCM **(c,d)** or TCGA-GBM **(e,f)** patients. Statistical-significance threshold was *p<0.05. For different gene-signatures, see **Supplementary-Table.2**.

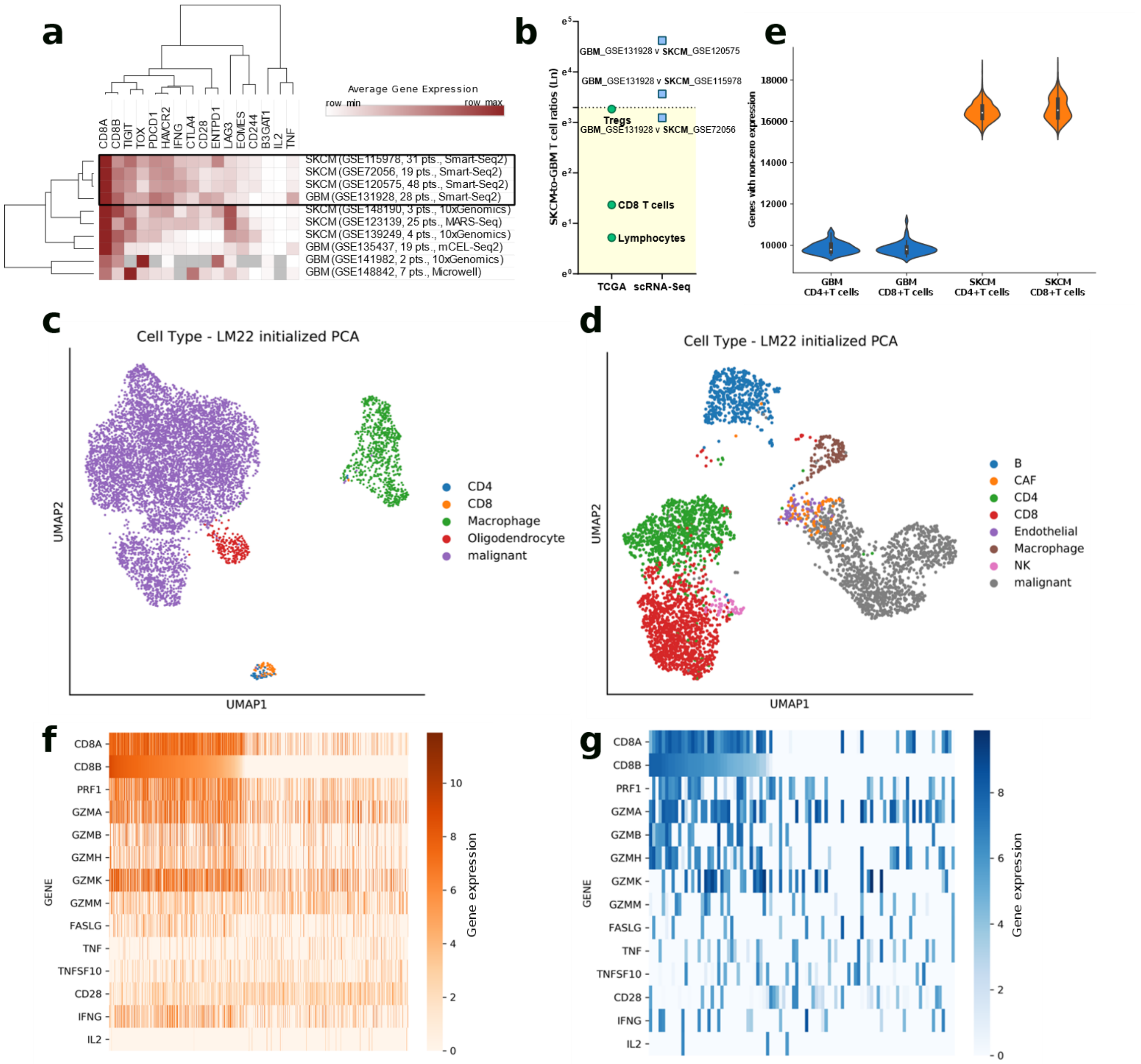


**Supplementary Figure S5. (a)** Results of a literature-wide meta-analysis for previously published single-cell (sc)RNASeq maps profiling SKCM or GBM patient-derived tumours for immune cell-compartment, especially T cells. This analysis was conducted via the Tumor Immune Single Cell Hub (TISCH) database,[2] a large-scale curated database integrating single-cell transcriptomics data from >2 million cells across at least 76 high-quality tumour-datasets from 28 cancer-types.[2] Heatmap indicates the coverage of csCD8$^{+}$T cell-signature (average gene-expression) within CD8$^{+}$T cells' single-cell transcriptomics data across all the scRNASeq-datasets. The black-outlined box indicates the scRNASeq-datasets with the best gene-by-gene coverage for csCD8$^{+}$T cell-signature across SKCM or GBM patient-cohorts. Grey box indicates lack of expression or gene "drop-outs". **(b)** SKCM-to-GBM ratios for indicated CIBERSORT-LM22/CRI-iATLAS derived immune-cell fractions within TCGA-datasets (green circles) or number of total T cells profiled across major scRNASeq datasets (blue squares) previously delineated (**Figure S5a**). **(c,d)** UMAP-visualizations of indicated immune cell-types, based on CIBERSORT-LM22 gene-signatures, in GBM **(c)** or SKCM **(d)** scRNASeq patient-cohorts.[3,4] Herein, for SKCM, scRNASeq profiling was done in 19-patients to derive 1252 melanoma-cells, 61 cancer-associated fibroblasts (CAFs), 2069 T-cells, 515 B-cells, 52 NK-cells, 126 macrophages and 65 endothelial cells.[3] For this profiling, the SKCM-tumour tissue originated from primary-melanoma (n=1) or metastasis procured from lymphoid tissue (n=9 from lymph-nodes, n=1 from spleen), subcutaneous/intramuscular tissue (n=5) or gastrointestinal tract (n=3).[3] Similarly, for GBM, scRNASeq profiling was done in 28-patients to derive 6863 GBM-cancer cells, 219 oligodendrocytes, 754 macrophages and 94 T-cells.[4] For this profiling, the GBM-tumour tissue originated from IDH-wild-type adult (n=20) and paediatric (n=8) GBM-patients.[4] **(e)** Visualization of gene-dropout. The number of genes with non-zero expression are shown for GBM and SKCM associated $CD4^{+}/CD8^{+}$ T single-cells. **(f,g)** Heatmap representation of indicated gene's expression levels across single T-cells (sorted by *CD8B*) within the SKCM **(f)** or GBM **(g)** scRNAseq patient-datasets.

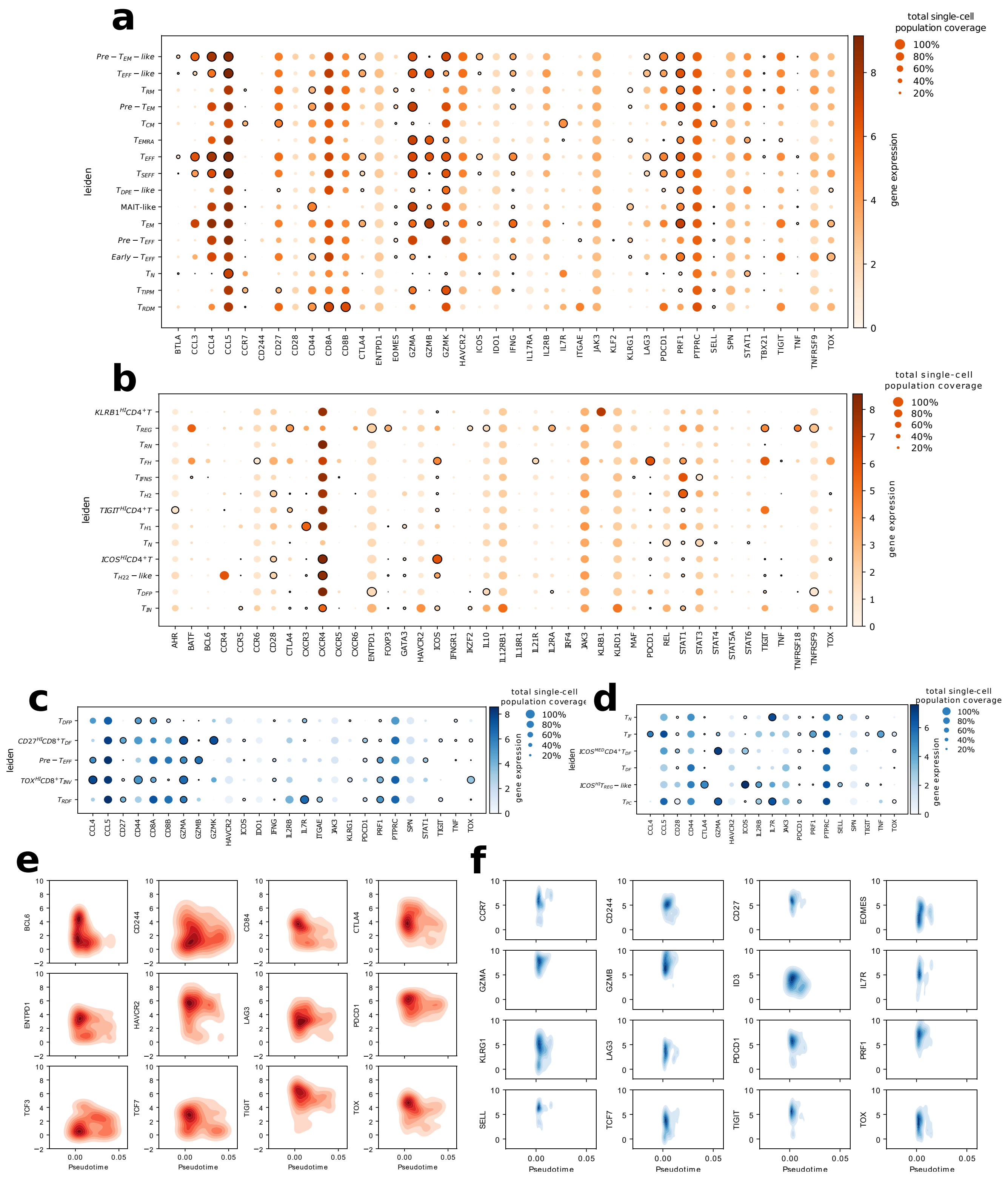


**Supplementary Figure S6. (a-d)** Dotplots highlighting differences in expression of indicated marker genes used for annotating the different $CD8^+$ **(a,c)** or $CD4^+$ **(b,d)** T-cell subpopulations (see **Supplementary Box.1**) as identified by the Leiden analysis in SKCM **(a,b)** or GBM **(c,d)** scRNASeq datasets.[3,4] Solid black-borders indicate gene majorly used for annotating the subpopulation, based on available literature. **(e,f)** Density plots for a panel of genes-of-interest using the scRNAseq data analysed with STREAM for deeper appraisal of the effector-progenitor trajectory[5,5,6] (**e**) or memory precursoreffector like trajectory[7] with central/resident-memory and immunotolerant-phenotypes (**f**).[8] The X-axis shows the inferred pseudotime, Y-axis represents the $\log_2$-transformed TPM expression.

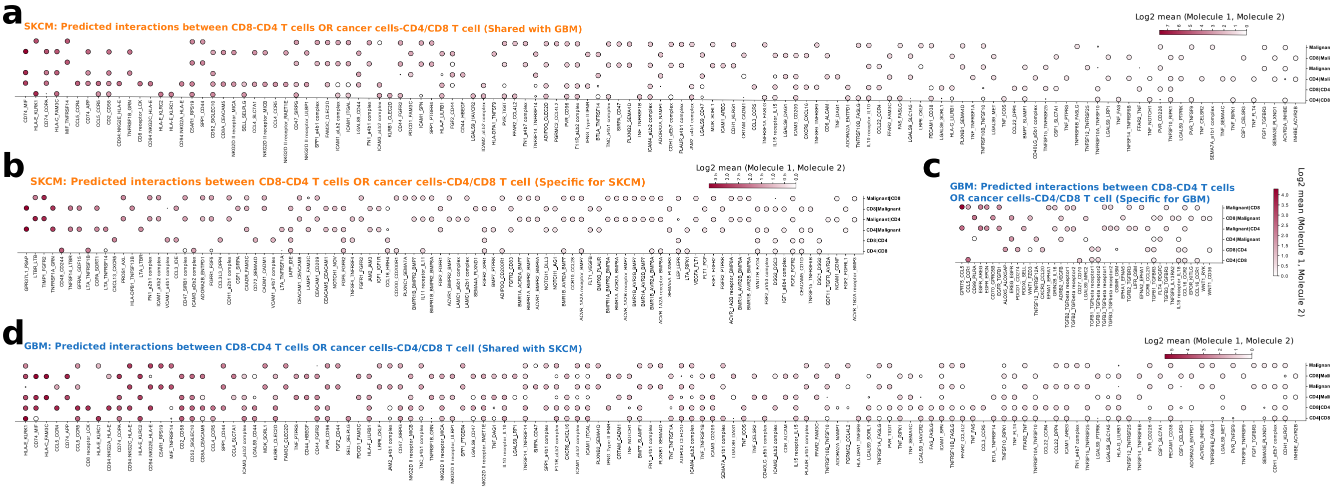


**Supplementary Figure S7.** Dotplots illustrating mean expression (colour) of the pairwise interaction and the probability (node size) of the interaction between ligand-receptor pairs (separated by "_") between indicated single-cell types (separated by "|") after running the CellPhoneDB pipeline.[9] Herein, we depict the interactions between $CD8^+$T cells::$CD4^+$T cells, cancer cells::$CD4^+$T cells or cancer cells::$CD8^+$T cells, specific for SKCM-patients **(a)**, specific for GBM-patients **(c)** or shared between SKCM and GBM patients **(b,d)** within the scRNASeq-dataset.[3,4]

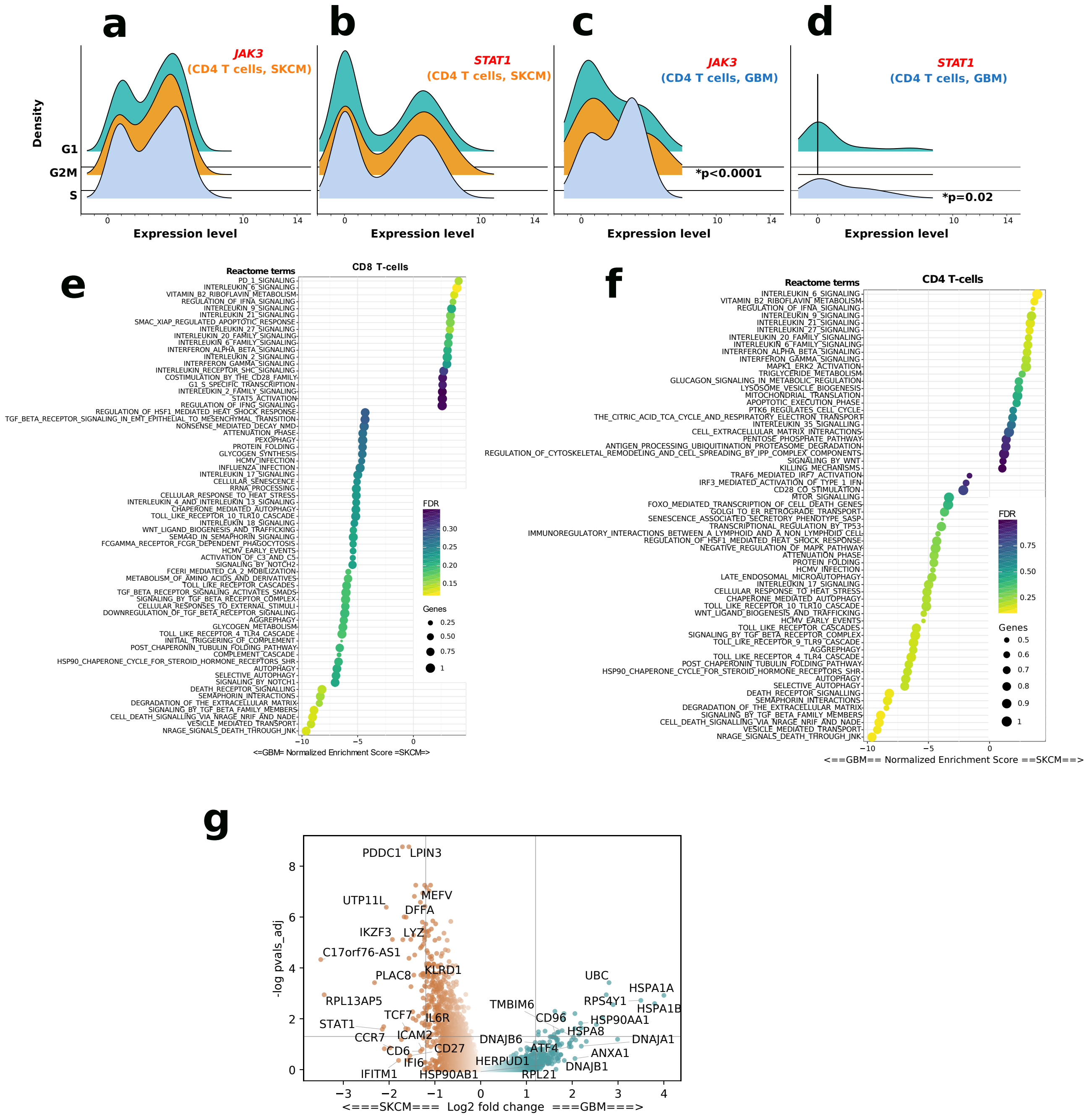


**Supplementary Figure S8. (a-d)** Ridge-plots illustrating the distribution of *JAK3/STAT1* expression in SKCM-CD4$^+$T single-cells **(a,b)** and GBM-CD4$^+$T single-cells **(c,d)** (statistics was estimated for SKCM vs. GBM comparison, for each gene per cell-cycle phase score; area-under-the-curve method combined with ANCOVA-analyses). **(e,g)** GSEA results for REACTOME pathways showing false-discovery rate (FDR) as colour intensity and normalized enrichment score on the X-axis between GBM and SKCM scRNASeq-data for CD8$^+$T single-cells **(e)** or CD4$^+$T single-cells **(g)**. Dot size represents the number of major contributor genes in the pathway-enrichment, scaled between 0 to 1. **(f)** Volcano plot comparing GBM-CD4$^+$T-cells to their SKCM counterparts, using scRNAseq data. P-values were Benjamini-Hochberg (BH)-adjusted. Statistical-significance threshold was *p<0.05. For different gene-signatures, see **Supplementary-Table.2**.

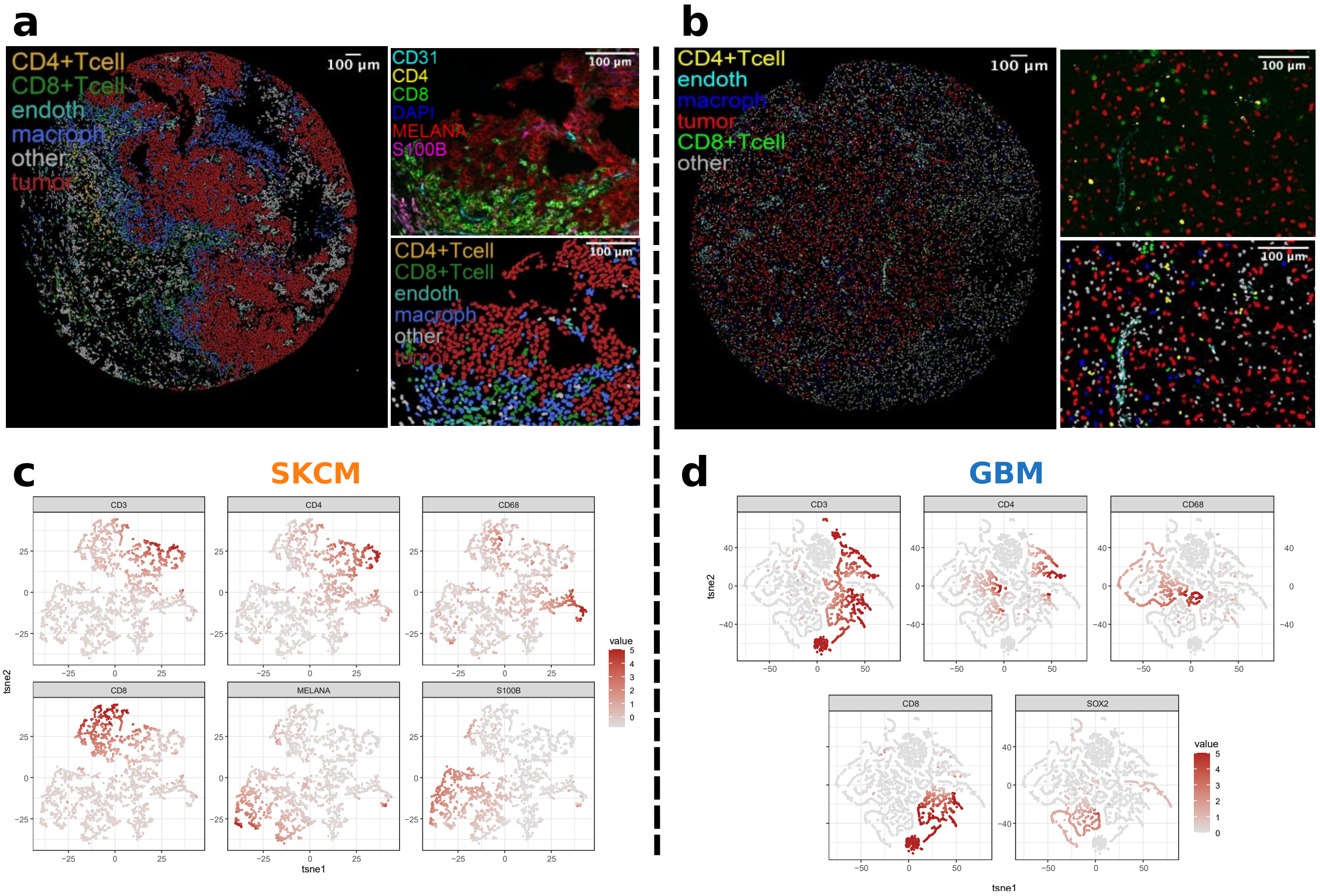


**Supplementary Figure S9. (a,b)** Digital representations of the indicated tissue sections from Figure 5a/5e, with the indication of the various cell-types. Inserts with magnified regions are shown on the right of each image, containing the actual overlayed images (top) and the digital representation (bottom). **(c,d)** tSNE plots corresponding to Figure 5d/5h in which the expression of the indicated markers are highlighted for SKCM (n=10-samples across 10-patients) or GBM (n=11-samples across 8-patients) patient-derived tumours.

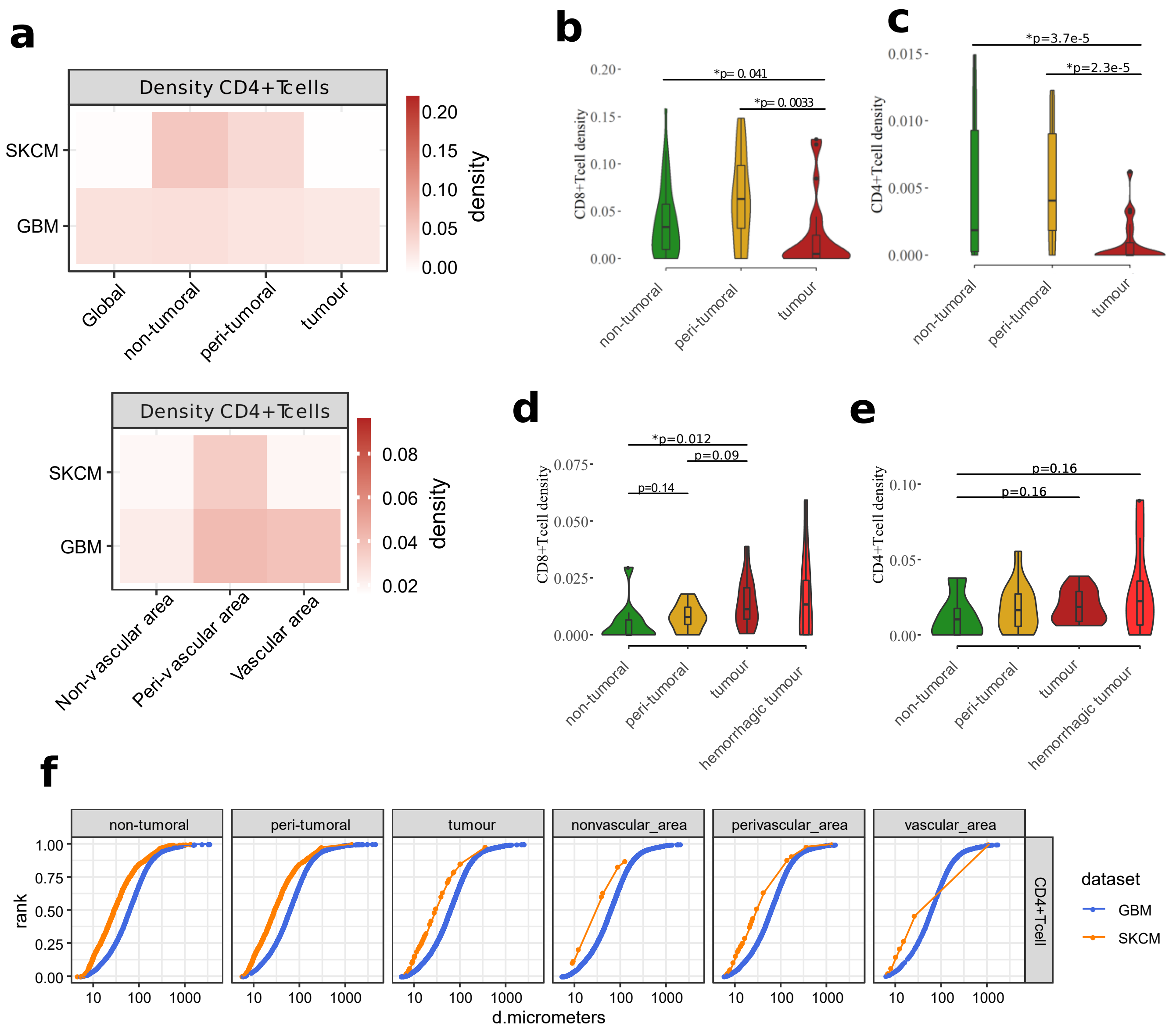


**Supplementary Figure S10. (a-e)** Density analysis of the CD4$^{+}$T cell and CD8$^{+}$T cell populations in GBM (n=11-samples across 8-patients) and SKCM (n=10-samples across 10-patients). Heat maps **(a)** show the relative distribution of CD4$^{+}$T-cells across the indicated areas within each tumour type. **(b-e)** Violin plots indicate the SKCM-CD8$^{+}$T cell **(b)**, SKCM-CD4$^{+}$T cell **(c)**, GBM-CD8$^{+}$T cell **(d)** or GBM-CD4$^{+}$T cell **(e)** density distribution across the indicated areas within each tumour-type per patient (Mann-Whitney test with Holm-method for multiple-testing comparison; FDR-adjusted p-values are indicated). **(f)** Density analysis defining the relative proportion of CD4$^{+}$T cells residing at the indicated distance (in micrometers/μm) from the nearest CD8$^{+}$T cell in SKCM or GBM tumours, across all analysed tissue-samples. Statistical-significance threshold was *p<0.05.

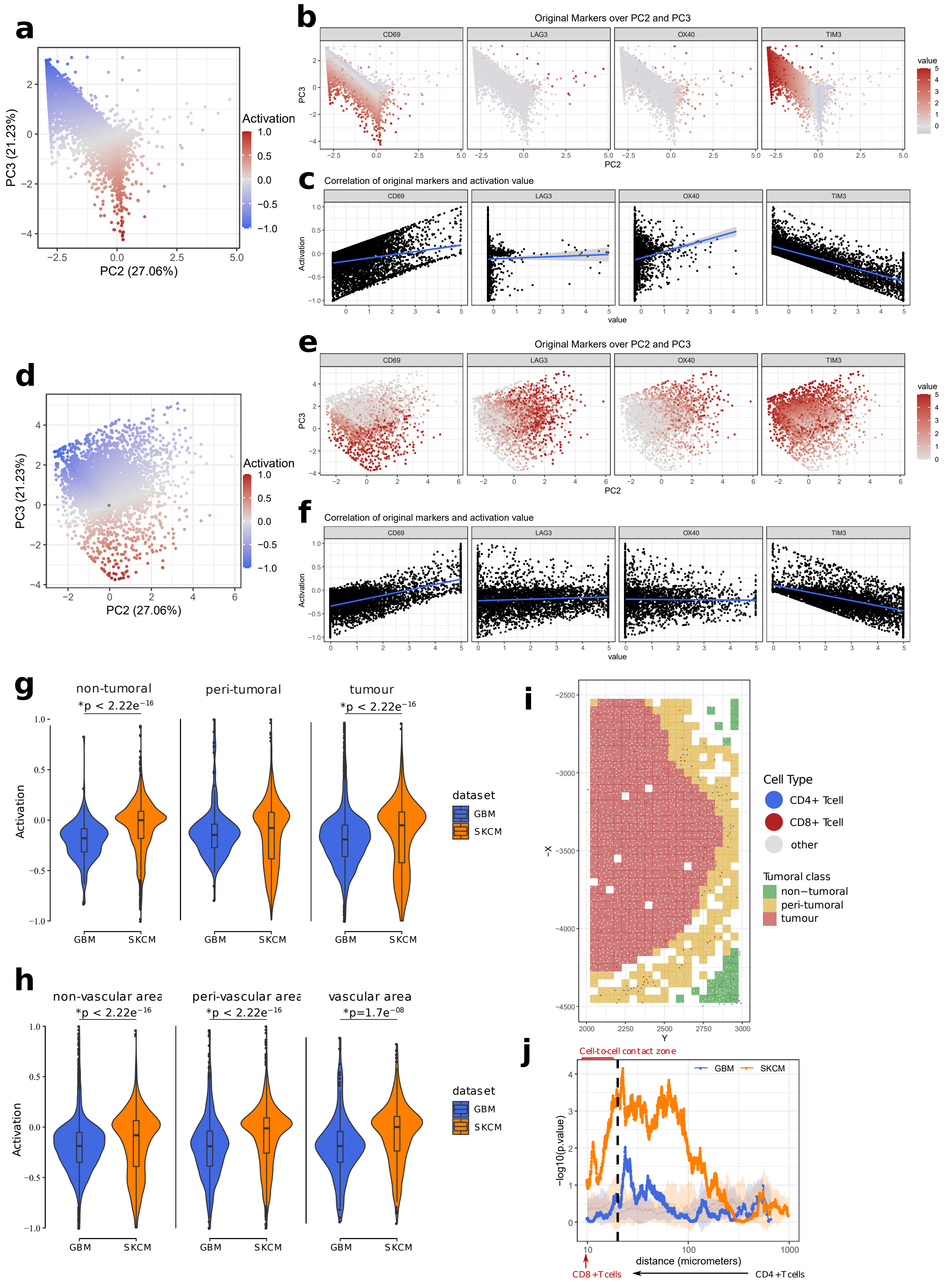


**Supplementary Figure S11. (a,d)** Dot plot representation showing the overall activation score distribution across all identified CD8$^{+}$T cells in SKCM (n=10-samples across 10-patients) **(a)** and GBM (n=11-samples across 8-patients) **(d)**. **(b,c,e,f)** Dot plot representation showing the break-up of the overall activation score across the different measured markers **(b, e)**, and their correlation to the overall activation score **(e,f)**. **(g,h)** Violin plots show the $CD8^{+}$T cell activation levels across the indicated areas within each cancer-type (Mann-Whitney test with Holm-method for multiple-testing comparison; FDR-adjusted p-values are indicated). **(i)** Schematic, digital representation of an SKCM tumour sample with the $CD4^{+}$ and $CD8^{+}$ T cell populations, and their relative distribution across the different areas, as indicated. **(j)** Line plots showing the corresponding p-values to the median activation score of each $CD8^{+}$T cell relative to the distance from the nearest $CD4^{+}$T cell in each indicated cancer-type (one-tailed t-test; FDR adjusted p-values are indicated) (corresponding to **Figure 5l**).

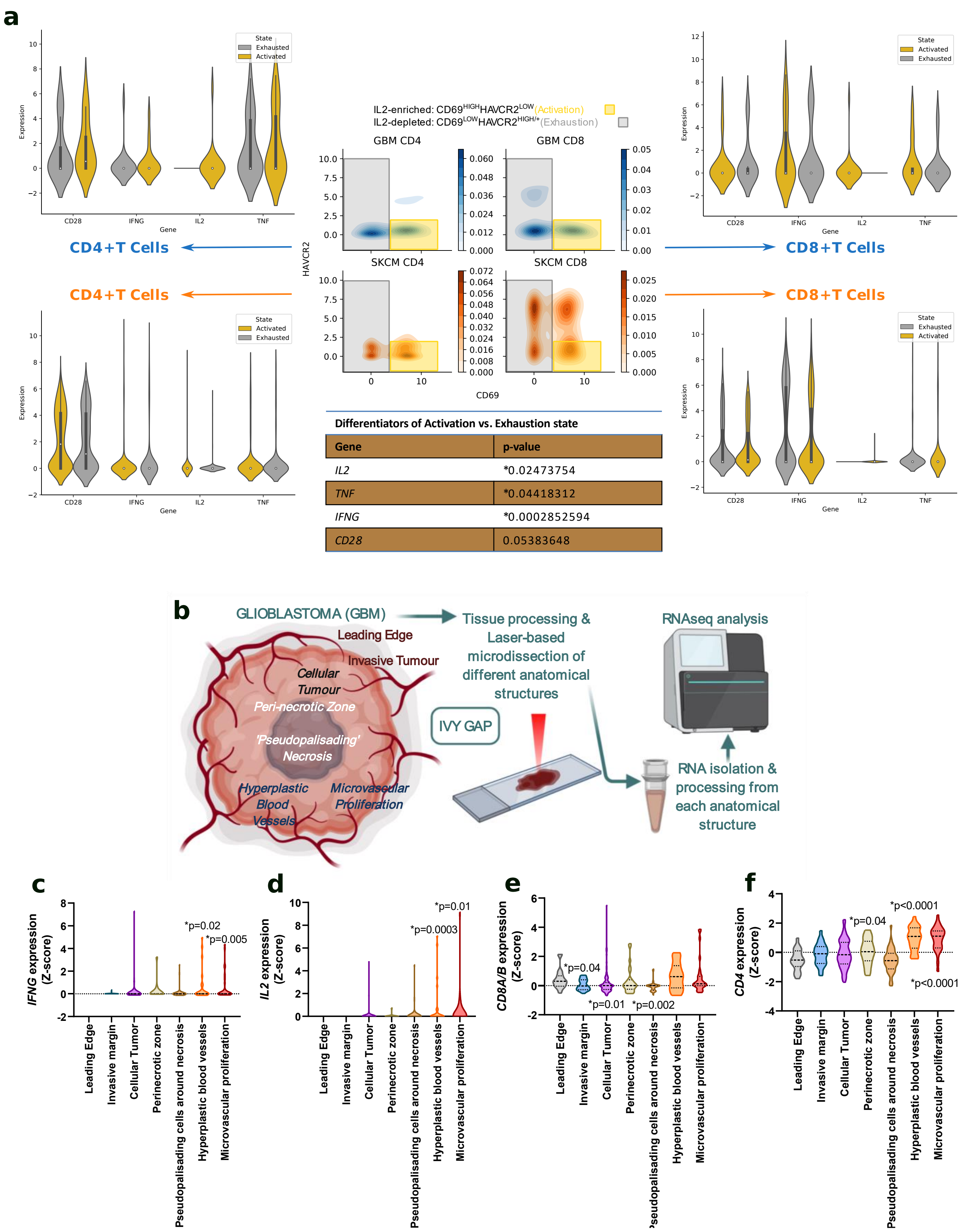


**Supplementary Figure S12. (a)** CD4$^+$/CD8$^+$T single-cell level appraisal of the *CD69* vs. *HAVCR2* activation/exhaustion-model. *CD69*$^{LOW}$*HAVCR2*$^{HIGH/+}$ and *CD69*$^{HIGH}$*HAVCR2*$^{LOW}$ status was annotated as IL2-exhausted and IL2-activated based on "targeted" analyses of *IL2*, *IFNG*, *CD28* or *TNF* in these subpopulations (see, gates in the density plots). P-values were obtained with MAST to estimate significance between activated and exhausted state. **(b)** Schematic overview of sampling, data procurement and analyses strategy of the IVY-GAP consortium's laser-assisted GBM tumour-tissue microdissection and anatomical sites-associated spatial transcriptomics.[10] **(c-f)** Distribution of the z-score standardized expression of indicated genes based on the anatomical location in the GBM tumour micro-environment as annotated by the IVY-GAP consortium[10] (Kruskal-Wallis ANOVA test with two-stage linear step-up procedure of Benjamini, Krieger and Yekutieli).

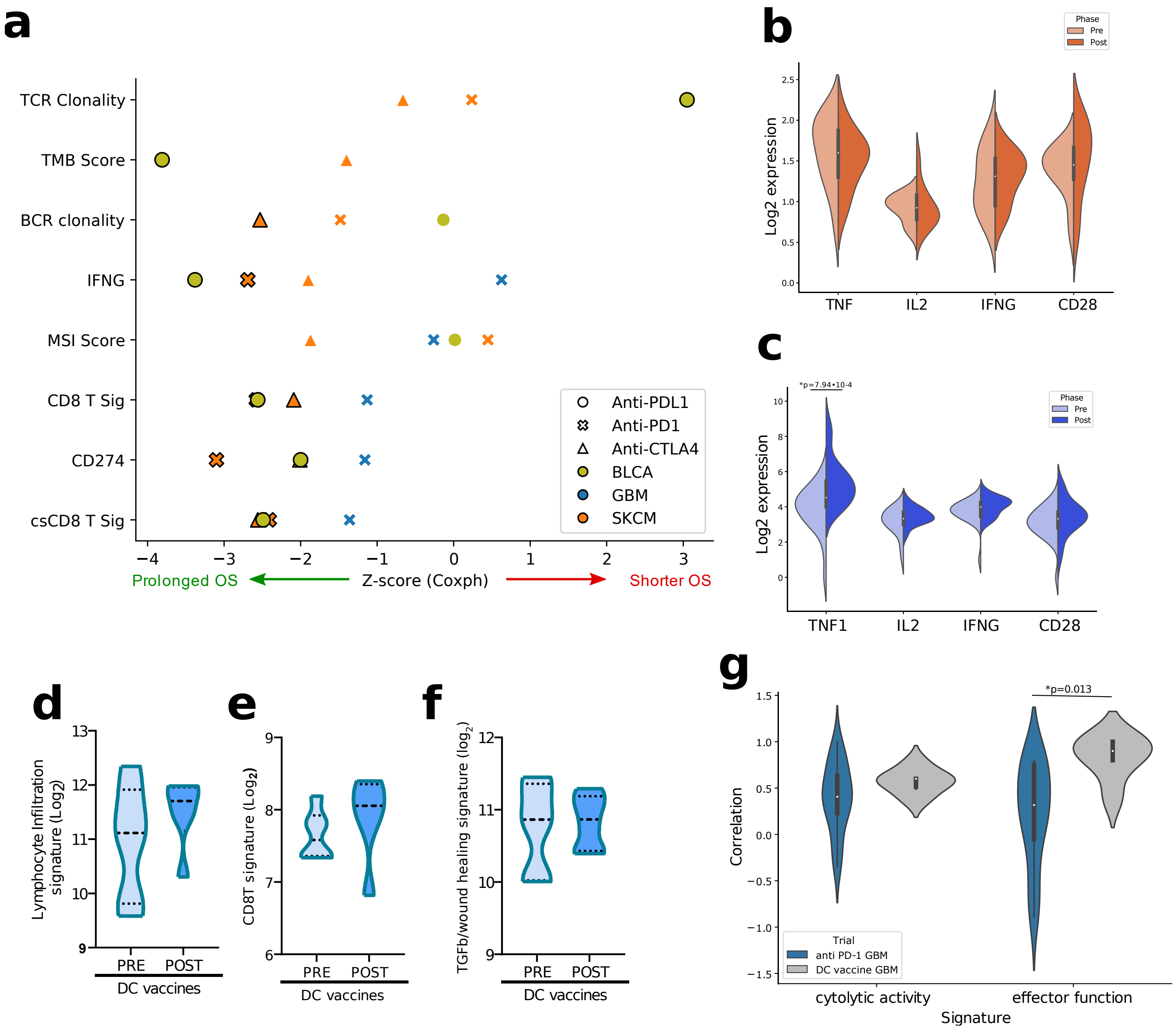


**Supplementary Figure S13. (a)** Clinical-trials involving pre-ICB (anti-PD1, anti-PDL1 or anti-CTLA4) treatment derived tumour-tissue from BLCA (n=298),[11] GBM (n=17)[12] or SKCM (anti-PD1 trial,[13] n=41; anti-CTLA4 trial,[14] n=42) patients were used for creating a comparison between the indicated biomarkers detected in pre-treatment tumour-tissue (Y-axis) and patient overall-survival (OS) estimated post-ICB treatment (OS represented as z-score in Coxph regression, X-axis) via the TIDE biomarker analyses platform.[15] Biomarker symbols with bold-outline indicates *p<0.05. The arrows at the bottom indicate prolonged or shortened OS, post-ICB treatment. Herein, TCR is T cell receptor, BCR is B cell receptor, TMB is tumour-mutational burden, MSI is micro-satellite instability. Clinical-trials involving pre/post-ICB (anti-PD1) treatment derived tumour-tissue from SKCM (Pre, n=13; Post, n=11)[16] or GBM (Pre, n=24; Post, n=25)[17] patients were used for creating violin-plots for indicated genes within SKCM **(b)** or GBM **(c)** datasets (Welch's t-test). Clinical-trial involving pre/post-dendritic cell (DC)-vaccination derived tumour-tissue from GBM (Pre, n=6; Post, n=6) patients was used for creating: **(d-f)** violin-plots for indicated signature-levels (Mann-Whitney test); and **(g)** Violin-plots for correlations between pre- and post- treatment levels of cytolytic activity signature (*PRF1, GZMA, GZMB, GZMH, GZMK, TNFSF10*) and effector function signature (*IFNG, IL2, CD28*) in above anti-PD1 immunotherapy or DC-vaccination clinical trials in GBM patients. Statistical-significance threshold was *p<0.05.
