## Supplemental box 1 for "Immunogenomic, single-cell and spatial dissection of CD8^+^T cell exhaustion reveals critical determinants of cancer immunotherapy"

**Supplementary BOX 1.** The annotation of CD4$^{+}$/CD8$^{+}$T cell sub-populations based on scRNASeq-data analyses.

**Abbreviations:** Overall expression representation of; HI, high-level; MED, medium-level; LO, low-level; +, minimal-level; MIN, minimal-to-negligible level; and NULL, negligible-to-zero level.

**SKCM-CD8$^{+}$T cells**

**Cluster 0 Pre-T$_{EM}$-like (Pre effector-memory like)** IFNG$^{MED}$, GZMA$^{HI}$, PRF1$^{HI}$, KLRG1$^{+}$, EOMES$^{+}$, TBX21$^{+}$, CCR7$^{NULL}$, SELL$^{MIN}$, CD44$^{MED}$

**Cluster 1 T$_{EFF}$-like (Effector-like)** TBX21$^{+}$, TNF$^{NULL}$, IFNG$^{MED}$, PRF1$^{HI}$, GZMA$^{HI}$, GZMB$^{HI}$, GZMK$^{MED}$, CCL5$^{HI}$, CCL3$^{LO}$, CCL4$^{MED}$, BTLA$^{+}$, CTLA4$^{MED}$, PDCD1$^{MED}$, ICOS$^{LO}$, LAG3$^{MED}$

**Cluster 2 T$_{RM}$ (Resting-memory)** IFNG$^{+}$, GZMB$^{MIN}$, GZMA$^{MIN}$, PRF1$^{HI}$, KLRG1$^{LO}$, TOX$^{MED}$, EOMES$^{LO}$, TBX21$^{MIN}$, TNF$^{NULL}$, CCR7$^{MIN}$, SELL$^{MIN}$, CD44$^{MED}$

**Cluster 3 Pre-T$_{EM}$ (Pre effector-memory)** IFNG$^{MED}$, GZMA$^{HI}$, PRF1$^{HI}$, KLRG1$^{LO}$, EOMES$^{MED}$, TBX21$^{NULL}$, TNF$^{NULL}$, CCR7$^{NULL}$, SELL$^{MIN}$, CD44$^{MED}$

**Cluster 4 T$_{CM}$ (Central-memory)** CCR7$^{MED}$, SELL$^{HI}$, IL7R$^{HI}$, CD27$^{HI}$, TBX21$^{NULL}$, EOMES$^{LO}$, PRF1$^{LO}$, GZMA/B$^{MIN}$

**Cluster 5 T$_{EMRA}$ (Effector memory re-expressing CD45RA)** CCR7$^{NULL}$, CD27$^{LO}$, GZMA/B$^{HI}$, GZMK$^{MED}$, PRF1$^{MED}$, IFNG$^{+}$, TNF$^{+}$, TOX$^{LO}$, PDCD1$^{MIN}$, CTLA4$^{MIN}$

**Cluster 6 T$_{EFF}$ (Effector)** TBX21$^{+}$, TNF$^{+}$, IFNG$^{HI}$, PRF1$^{HI}$, GZMA$^{HI}$, GZMB$^{HI}$, GZMK$^{HI}$, CCL5$^{HI}$, CCL3$^{HI}$, CCL4$^{HI}$, BTLA$^{+}$, CTLA4$^{HI}$, PDCD1$^{HI}$, ICOS$^{MED}$, LAG3$^{MED}$

**Cluster 7 T$_{SEFF}$ (Suppressed-effector)** TBX21$^{NULL}$, TNF$^{NULL}$, IFNG$^{+}$, STAT1$^{LO}$, PRF1$^{HI}$, GZMA$^{HI}$, GZMB$^{MIN}$, GZMK$^{HI}$, CCL5$^{HI}$, CCL3$^{MED}$, CCL4$^{HI}$, BTLA$^{NULL}$, CTLA4$^{LO}$, PDCD1$^{HI}$, ICOS$^{LO}$, LAG3$^{MED}$

**Cluster 8 T$_{DPE}$-like (Double-positive effector-like)** KLRG1$^{MED}$, IL7R$^{LO}$, STAT1$^{HI}$, TBX21$^{NULL}$, TIGIT$^{MED}$, PRF1$^{MED}$, PDCD1$^{LO}$, GZMA$^{LO}$, GZMB$^{NULL}$, GZMK$^{HI}$

**Cluster 9 MAIT-like (Mucosal associated invariant T cells)** CD8A$^{MIN}$, CD8B$^{NULL}$, CD44$^{HI}$, CCR7$^{NULL}$, SELL$^{MIN}$

**Cluster 10 T$_{EM}$ (Effector-memory)** IFNG$^{HI}$, GZMB$^{HI}$, GZMA$^{MED}$, PRF1$^{HI}$, KLRG1$^{LO}$, EOMES$^{+}$, TBX21$^{+}$, TNF$^{+}$, CCR7$^{NULL}$, SELL$^{NULL}$, CD44$^{MED}$

**Cluster 11 Pre-T$_{EFF}$ (Pre-effector)** TBX21$^{NULL}$, TNF$^{NULL}$, IFNG$^{MED}$, PRF1$^{MED}$, GZMA$^{HI}$, GZMB$^{NULL}$, GZMK$^{HI}$, CCL5$^{HI}$, CCL3$^{MIN}$, CCL4$^{HI}$, BTLA$^{+}$, CTLA4$^{MIN}$, PDCD1$^{LO}$, ICOS$^{LO}$, LAG3$^{LO}$

**Cluster 12 Early-T$_{EFF}$ (Early-effector)** TBX21$^{NULL}$, TNF$^{NULL}$, IFNG$^{MED}$, PRF1$^{HI}$, GZMA$^{LO}$, GZMB$^{MIN}$, GZMK$^{MIN}$, CCL5$^{HI}$, CCL3$^{LO}$, CCL4$^{HI}$, BTLA$^{+}$, CTLA4$^{LO}$, PDCD1$^{MED}$, ICOS$^{LO}$, LAG3$^{MED}$

**Cluster 13 T$_{N}$ (Naive)** CCR7$^{MED}$, IL7R$^{HI}$, SELL$^{MED}$, CTLA4$^{NULL}$, TOX$^{NULL}$, EOMES$^{NULL}$, GZMA/B/K$^{NULL}$, PRF1$^{LO}$, ICOS$^{MIN}$, IFNG$^{NULL}$, TNF$^{NULL}$

**Cluster 14 T$_{TIPM}$ (Tumour-infiltrating peripheral-memory)** SELL$^{LO}$, PRF1$^{LO}$, GZMA$^{MED}$, GZMB$^{NULL}$, GZMK$^{HI}$, IFNG$^{MIN}$, TNF$^{NULL}$, CCR7$^{MED}$, CD27$^{MED}$, TBX21$^{NULL}$

**Cluster 15 T$_{RDM}$ (Resident-memory)** ITGAE$^{HI}$, KLRG1$^{MIN}$, CCR7$^{NULL}$, SELL$^{MIN}$, KLF2$^{NULL}$, EOMES$^{NULL}$, TBX21$^{NULL}$

**SKCM-CD4$^{+}$T cells**

**Cluster 0 KLRB1$^{HI}$CD4$^{+}$T**

**Cluster 1 T$_{REG}$ (Regulatory)** FOXP3$^{HI}$, TIGIT$^{HI}$, IKZF2$^{MED}$, CTLA4$^{HI}$, IL2RA$^{MED}$, TNFRSF18$^{HI}$, TNFRSF9$^{MED}$, ENTPD1$^{MED}$, IL10$^{MED}$

**Cluster 2 T$_{RN}$ (Resting-Naive)** CXCR4$^{HI}$, CTLA4$^{NULL}$, CXCR3$^{NULL}$, CXCR5$^{NULL}$, CCR5$^{NULL}$, FOXP3$^{NULL}$, GATA3$^{MIN}$, ICOS$^{NULL}$, IKZF2$^{NULL}$, PDCD1$^{NULL}$, STAT1$^{LO}$

**Cluster 3 T$_{FH}$ (Follicular-helper)** BATF$^{HI}$, BCL6$^{MED}$, MAF$^{HI}$, PDCD1$^{HI}$, STAT3$^{MED}$, ICOS$^{HI}$, CXCR5$^{LO}$;

**Cluster 4 T$_{IFNS}$ (Interferon-stimulated)** STAT1$^{HI}$, STAT3$^{MED}$, STAT4$^{LO}$, STAT6$^{LO}$, PDCD1$^{NULL}$, CTLA4$^{MIN}$, REL$^{MED}$

**Cluster 5 T$_{H2}$ (type-II polarised)** GATA3$^{MED}$, FOXP3$^{NULL}$, CXCR3$^{NULL}$

**Cluster 6 TIGIT$^{HI}$CD4$^{+}$T**

**Cluster 7 T$_{H1}$ (Type-I polarised)** CXCR3$^{HI}$, CCR5$^{LO}$, CXCR6$^{+}$, ICOS$^{LO}$, TNF$^{+}$, CTLA4$^{LO}$, CD28$^{MED}$, TOX$^{MIN}$, STAT1$^{MED}$, PDCD1$^{LO}$

**Cluster 8 T$_{N}$ (Naive)** CXCR4$^{LO}$, CD28$^{LO}$, CCR5$^{NULL}$, CXCR3$^{NULL}$, FOXP3$^{NULL}$, GATA3$^{MIN}$, ICOS$^{MIN}$, PDCD1$^{NULL}$, STAT1$^{LO}$, TIGIT$^{MIN}$, TNF$^{NULL}$

**Cluster 9 ICOS$^{HI}$CD4$^{+}$T**

**Cluster 10 T$_{H22}$-like (Type-22 polarised-like)** CCR4$^{HI}$ AHR$^{MED}$ STAT1$^{LO}$, CTLA4$^{MIN}$, PDCD1$^{NULL}$, ICOS$^{MED}$

**Cluster 11 T$_{DFP}$ (Dysfunctional-precursor)** PDCD1$^{MED}$, CCR6$^{MED}$, STAT1$^{NULL}$, IL21R$^{LO}$, TNF$^{NULL}$, ICOS$^{NULL}$,

**Cluster 12 T$_{IN}$ (Inflammed-naive)** CXCR4$^{MED}$, CD28$^{LO}$, CCR5$^{LO}$, CXCR3$^{MIN}$, FOXP3$^{NULL}$, GATA3$^{LO}$, ICOS$^{MED}$, PDCD1$^{MIN}$, STAT1$^{MED}$, TIGIT$^{MED}$, TOX$^{MED}$, TNF$^{NULL}$

**GBM-CD8$^{+}$T cells**

**Cluster 0 T$_{DFP}$ (Dysfunctional-precursor)** CD8A$^{MED}$, CD8B$^{LO}$, IL7R$^{MED}$, KLRG1$^{LO}$, IFNG$^{NEG}$, GZMB$^{NULL}$, GZMA$^{NULL}$, PRF1$^{LO}$, TNF$^{LO}$, CD44$^{MED}$

**Cluster 1 CD27$^{HI}$CD8$^{+}$T$_{DF}$ (Dysfunctional)** IL7R$^{LO}$, CD27$^{HI}$, TOX$^{+}$, TIGIT$^{+}$, PDCD1$^{LO}$, GZMA$^{HI}$, GZMB$^{NEG}$, GZMK$^{HI}$, PRF1$^{LO}$, ICOS$^{LO}$, IFNG$^{NULL}$, TNF$^{LO}$

**Cluster 2 Pre-T$_{EFF}$ (Pre-effector)** TNF$^{NEG}$, CD27$^{NULL}$, ICOS$^{NEG}$, IFNG$^{LO}$, STAT1$^{LO}$, PRF1$^{MED}$, GZMA$^{MED}$, GZMB$^{HI}$, CCL4$^{MED}$, TOX$^{NULL}$, PDCD1$^{LO}$

**Cluster 3 TOX$^{HI}$CD8$^{+}$T$_{INV}$ (Invariant)** CD8A$^{LO}$, CD8B$^{LO}$, TOX$^{HI}$, CCL4$^{HI}$, CD27$^{LO}$, CD44$^{HI}$, IFNG$^{+}$, GZMB$^{NEG}$, GZMA$^{HI}$, PRF1$^{MED}$, KLRG1$^{MED}$, IL7R$^{NULL}$

**Cluster 4 T$_{RDF}$ (Resident-dysfunctional)** IL7R$^{HI}$, ITGAE$^{MED}$, KLRG1$^{NULL}$, CD27$^{MED}$, GZMA$^{MED}$, PRF1$^{MED}$, ICOS$^{LO}$, IFNG$^{NEG}$, STAT1$^{NULL}$, TNF$^{NULL}$, TOX$^{NEG}$, PDCD1$^{LO}$, TIGIT$^{LO}$

**GBM-CD4$^+$T cells**

**Cluster 0** **$T_N$ (Naive)** IL7R$^{HI}$, SELL$^{HI}$, CD28$^{LO}$, CTLA4$^{NEG}$, GZMA$^{NULL}$, PRF1$^{LO}$, ICOS$^{LO}$, PDCD1$^{LO}$, TIGIT$^{LO}$, TNF$^{NULL}$

**Cluster 1** **$T_{IF}$ (Inflammatory)** CCL4$^{HI}$, TNF$^{HI}$, CD28$^{LO}$, CTLA4$^{NEG}$, GZMA$^{LO}$, PRF1$^{HI}$, ICOS$^{NULL}$, PDCD1$^{LO}$, TIGIT$^{LO}$, TOX$^{MED}$

**Cluster 2** **ICOS$^{MED}$CD4$^+$T$_{DF}$ (Dysfunctional)** ICOS$^{MED}$, CTLA4$^{NEG}$, TOX$^{LO}$, CD28$^{LO}$, PDCD1$^{LO}$, GZMA$^{HI}$, PRF1$^{NEG}$

**Cluster 3** **$T_{DF}$ (Dysfunctional)** CTLA4$^{NEG}$, TOX$^{NEG}$, CD28$^{LO}$, ICOS$^{NULL}$, PDCD1$^{LO}$, GZMA$^{NULL}$, PRF1$^{NULL}$

**Cluster 4** **ICOS$^{HI}$T$_{REG}$-like (Regulatory-like)** ICOS$^{HI}$, CTLA4$^{HI}$, IL2RB$^{HI}$, TIGIT$^{LO}$, SELL$^{MED}$, TNF$^{NEG}$, PRF1$^{LO}$, GZMA$^{NULL}$, PDCD1$^{NEG}$

**Cluster 5** **$T_{PC}$ (Precursor)** IL7R$^{HI}$, IL2RB$^{MED}$, CTLA4$^{NULL}$, TOX$^{LO}$, CD28$^{MED}$, ICOS$^{NEG}$, PDCD1$^{LO}$, GZMA$^{HI}$, PRF1$^{NULL}$
